# Mechanical model of muscle contraction. 3. The orientation of the levers belonging to the myosin heads in working stroke follows the same uniform law in all half-sarcomeres of an isometrically stimulated fiber

**DOI:** 10.1101/2019.12.16.878835

**Authors:** S. Louvet

## Abstract

A myosin II head is modelled during the working stroke (WS) by three rigid segments articulated between them: the motor domain (S1a), the lever (S1b) and the rod (S2). Hypothesis 4 introduced in accompanying Paper 2 states that the lever of a WS head moves in a fixed plane where the position of S1b is characterized by the angle θ. This assumption allows the geometrization of a cross-bridge, i.e. the poly-articulated chain consisting of five rigid segments: the actin filament (Afil), S1a, S1b, S2, and the myosin filament (Mfil). The equations established in Paper 2 are operative to calculate the number of heads potentially in WS for a Mfil surrounded by six Afil. In addition, the value of the angles θ of the levers belonging to these WS heads is accessible. This census leads to an integer number (N_p_) of angular positions (θi) distributed discretely between θ_up_ and θ_down_, the two values that delimit θ during the WS. The number of Mfil per half-sarcomere (hs) is estimated between 400 and 2000 depending on the typology, figures that induce Gaussian variability for each of the N_p_ values θi calculated for a single Mfil. By summing the Gaussian N_p_ densities and after normalization, we obtain a probability density (d_G_) of the continuous variable θ between θ_up_ and θ_down_. The function d_G_ is calculated for a random length of a hs between 1 and 1.1 μm where the binding rate of the myosin heads is maximum. From this reference length, the hs is shortened 11 times with a step of 1 nm, i.e. a total of 11 nm. For each shortening, a count of the new θi positions is performed, which leads to a new probability density d_G_. The classic statistical law that approximates these 12 distributions of θ is the Uniform law between θ_up_ and θ_down_. Other conditions and values given to the data of the algorithmic procedure lead to a similar result, hence the formulation of hypothesis 5: the distribution of the angle θ follows an identical uniform law in all the hs of a muscle fiber stimulated in isometric conditions.

## Introduction

If the swinging lever arm hypothesis [1] is corroborated, it becomes legitimate to question the nature of the distribution governing the orientation θ of the levers of the WS myosin heads in any hs belonging to an isometrically tetanized fiber. Two models were proposed based on two observation methods: 1/ a Gaussian distribution according to the fluorescence polarization measurement after labelling the lever by introducing pairs of cysteines [2,3,4]; 2/ a Uniform density determined using the intensity of the the first order meridional diffraction named M3, obtained by X-ray interference technique [5,6,7,8,9].

We approach the problem from a theoretical point of view on the basis of postulate 4 of our model. In the work of the isometric tetanus plateau mentioned above, the dispersion of θ is given between ± 20° and ± 25°, i.e. an interval of 40° to 50°, figures less than 70°, the value used to evaluate the maximum angular interval during the WS (δθ_Max_). First, our reasoning will focus on δθ_Max_ and then we will apply it to an interval less than δθ_Max_.

Hypothesis 4 of the displacement of the lever in a fixed plane stated and demonstrated in the accompanying Paper 2 allows the calculation of the geometric coordinates of the 3 segments modelling a WS head in the inter-filament space and thus makes it possible to count the potentialities of WS in a hs. This result leads to hypothesis 5 where the distribution of the angular position (θ) of the levers belonging to the WS heads is uniform regardless of the fixed length of the hs between 1 and 1.1 μm. A first corollary of hypothesis 5 requires that uniformity applies to any angular interval between θ_up_ and θ_down_, in particular to the δθ_T_ interval observed during the isometric tetanus plateau. Perturbation by a length step varying from one to a few tens of nanometers starts with tetanization with isometry of an isolated muscle fiber. The uniformity postulate explains why, after this disturbance, the length of the fiber being fixed again, the tension always rises to a value equal to T0, the tension of the isometric tetanus plateau, because the uniform law on the interval δθ_T_ is common to all hs of the fiber before and at the very end of the disturb (see accompanying Paper 5). From hypothesis 5, it follows that some hs can have the same uniform distribution while having a different length, varying between 1 and 1.1 μm, thus allowing a non-uniformity of the length of hs described by different authors [10,11].

A third consequence is that the number of myosin heads potentially in WS decreases linearly with the length of the sarcomere when it is between 2.2 and 3.3 μm [12].

## Methods

All acronyms appearing in the text are explained in Table 1. The O_Afil_X straight line is the longitudinal axis of the Afil. The O_Afil_Y° straight line is the axis passing through the centers of the Afil and Mfil (O_Afil_ and O_Mfil_) connected by the cross-bridge (Fig 1b), an axis perpendicular to OAfilX. The lever S1b belonging to a WS myosin head moves in a fixed plane OA_Afil_XY specified by the angle β between O_Afil_Y° and O_Afil_Y (Fig 1b). The angle β also defines the orientation of the binding site (Asit) located on the surface of the actin molecule represented by point A. The knowledge of a second angle (α) located in the O_Mfil_Y°Z° transverse plane is necessary for the complete geometrization of the crossbridge. The α parameter characterizes the angle of the attachment point (D) of the rod S2 on the Mfil with respect to O_Mfil_Y° (Fig 1b). In the O_Mfil_Y°Z° transverse plane, an Afil and a Mfil are represented by a red and green solid circle, respectively (Fig 1c). Over the entire length of the Mfil except the bare zone, the myosin molecules (Mmol) are distributed in 9 rows (Mrow) parallel to the O_Mfil_X axis. In the O_Mfil_Y°Z° transverse plane, these 9 Mrow are represented by 9 red lines radiating around the Mfil. A Mfil is located in the center of a regular hexagon, each of the 6 vertices of which is occupied by the center of an adjacent Afil,. This hexagon is divided into 6 quadrilaterals in the shape of kites of equal surface area. One or two Mrow are present in each of the six kites (Fig 1c).

**Fig 1:**
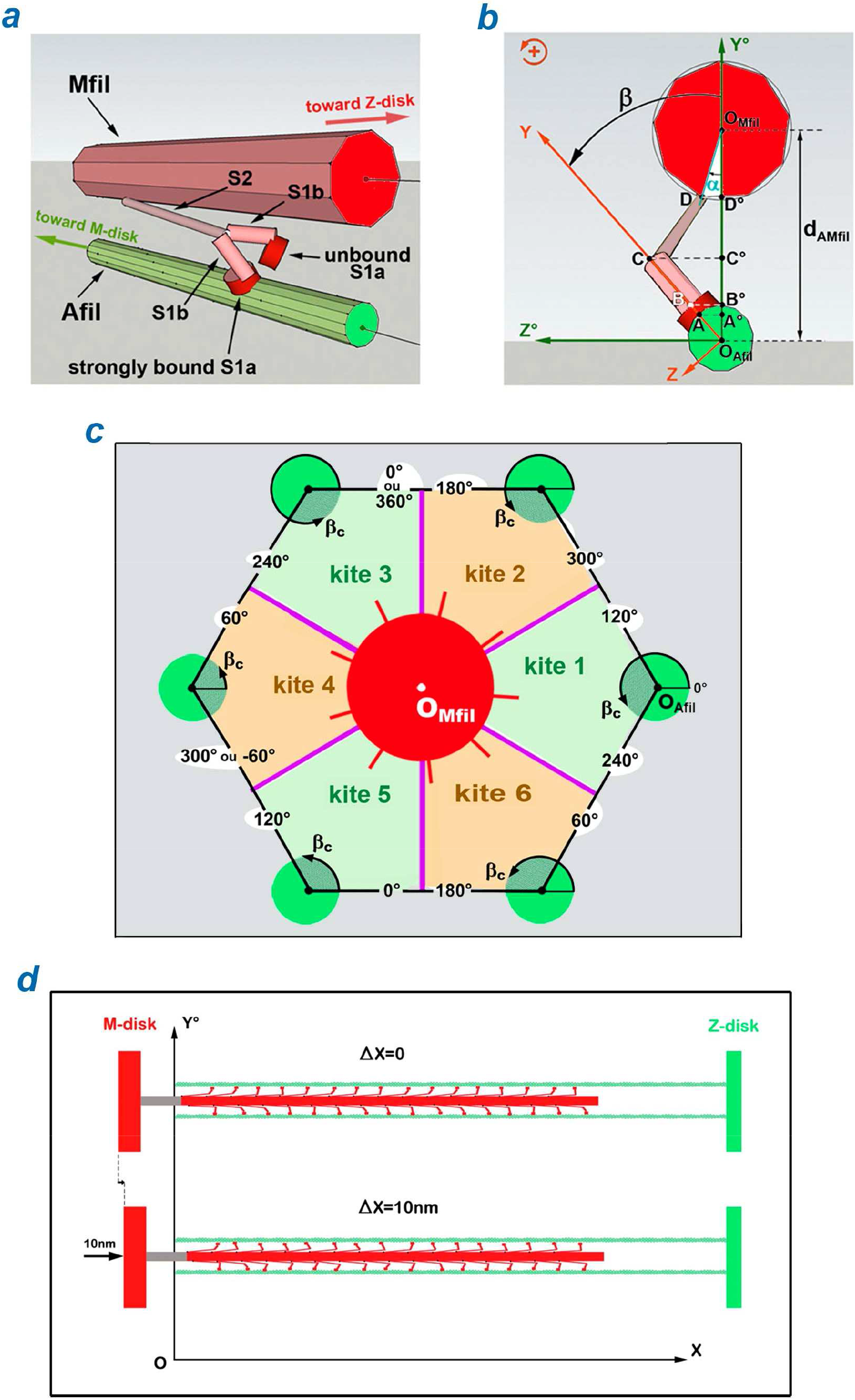
Cross-bridge geometrization. (a) Myosin molecule (Mmol) of which one of the 2 heads is in WS state. (b) Angles α and β. (c) A myosin filament (Mfil) surrounded by 6 homologous actin filaments (Afil) with characterization of the 6 kites. (d) Shortening by 10 nm of a half-sarcomere on the right (hsR) with fixed Z-disk.

**Table 1:**
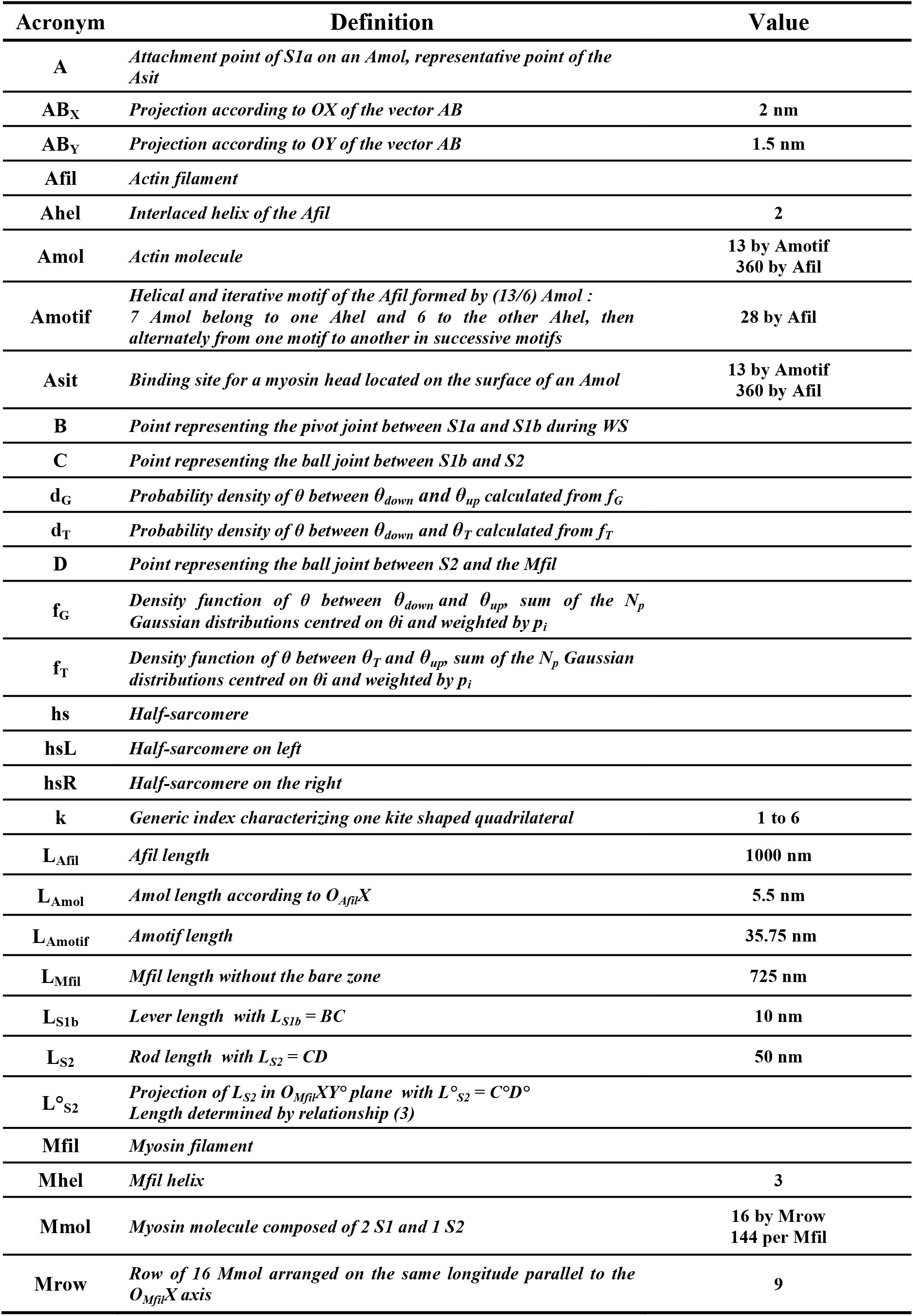
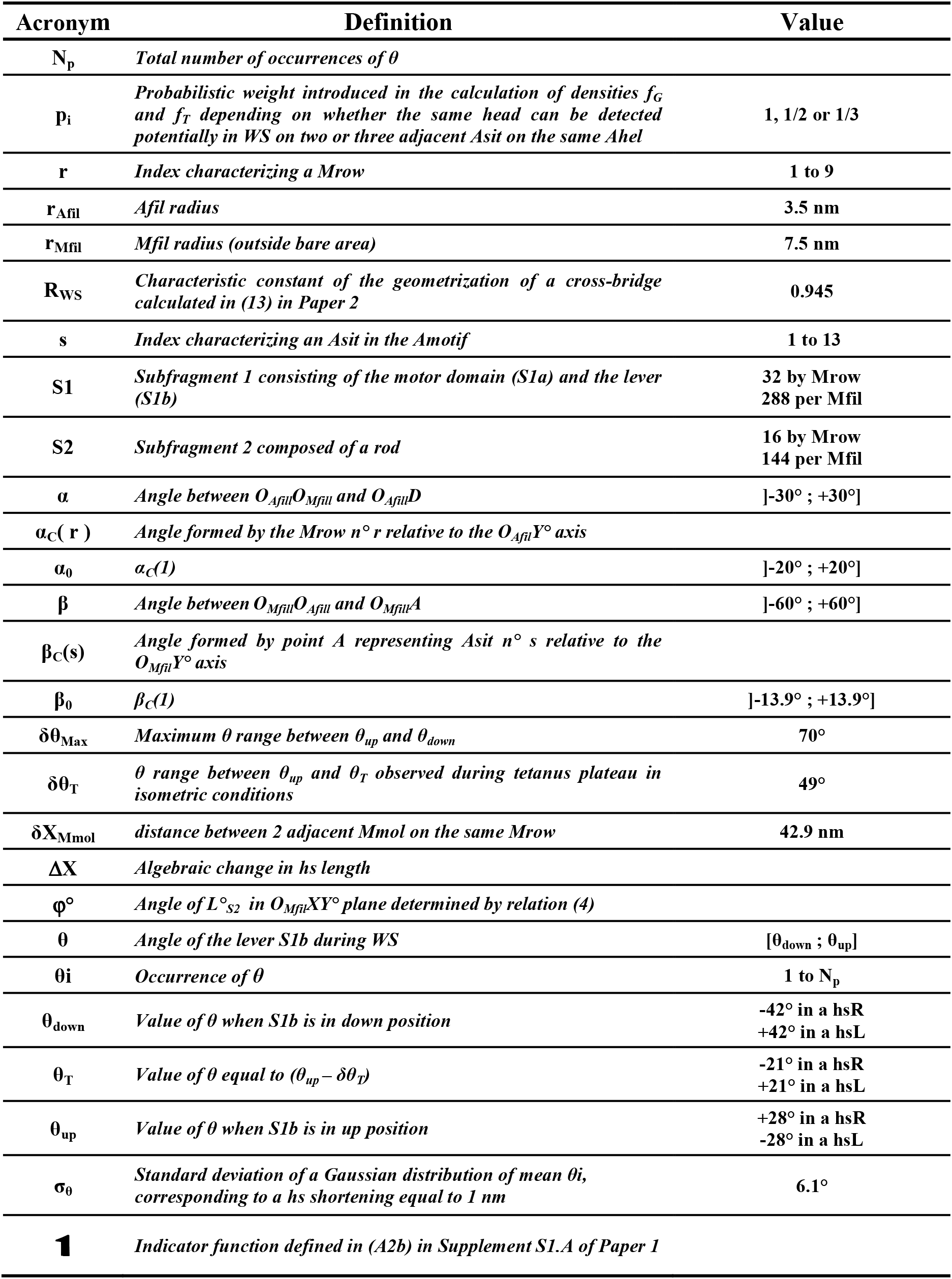
Geometric and numerical data characteristic of myosin filament (Mfil) and actin filament (Afil) in a half-sarcomere of a skeletal fiber; data from [13,14].

A hs is idealized by considering: 1/ all the Mfil of the hs are identical with the same orientation in the transverse plane of the hs; 2/ all the Afil of the hs are identical with the same orientation in the transverse plane of the hs.

### Determination of the angles α and β relative to a WS myosin head

The 9 Mrow of each Mfil are characterized by an r-index varying from 1 to 9. All the Mfil of the hs being oriented the same way, each of the 9 Mrow of each Mfil has a characteristic angle α_C_(r) identical in the OY°Z° plane. An Mfil is surrounded by 6 Afil (Fig 1c) and each Afil consists of the repetition of the same helical motif (Amotif) formed by (13/6) actin molecules (Amol), each Amol having a binding site (Asit). The 13 Asit of each of the 28 Amotif that follow one another along each Afil are characterized by an index s varying from 1 to 13; see Supplement S3.G. All the Afil of the hs being oriented the same way, each of the 13 Asit of each motif of each Afil has a characteristic angle βC(s) identical in the OY°Z° plane. We first assume that the points A and D of the poly-articulated chain that forms a myosin head potentially in WS (Figs 1b and 2A) are located in a single and same kite, i.e. that the Asit to which S1a is attached and the Mrow to which the rod S2 is attached belong to the same kite (Figs 2b and 2c). In a kite defined by adjacent Mfil and Afil, the α and β angles are defined with respect to the diagonal of the kite, the axis O_Mfil_O_Afil_, where O_Mfil_ and O_Afil_ are the respective centers of the Mfil and Afil. Knowing the respective values of α_C_ and β_C_, it is necessary to calculate the angles α and β according to the kite where the cross-bridge is located. The calculation method is developed in paragraph G.3 of Supplement S3.G.

**Fig 2.**
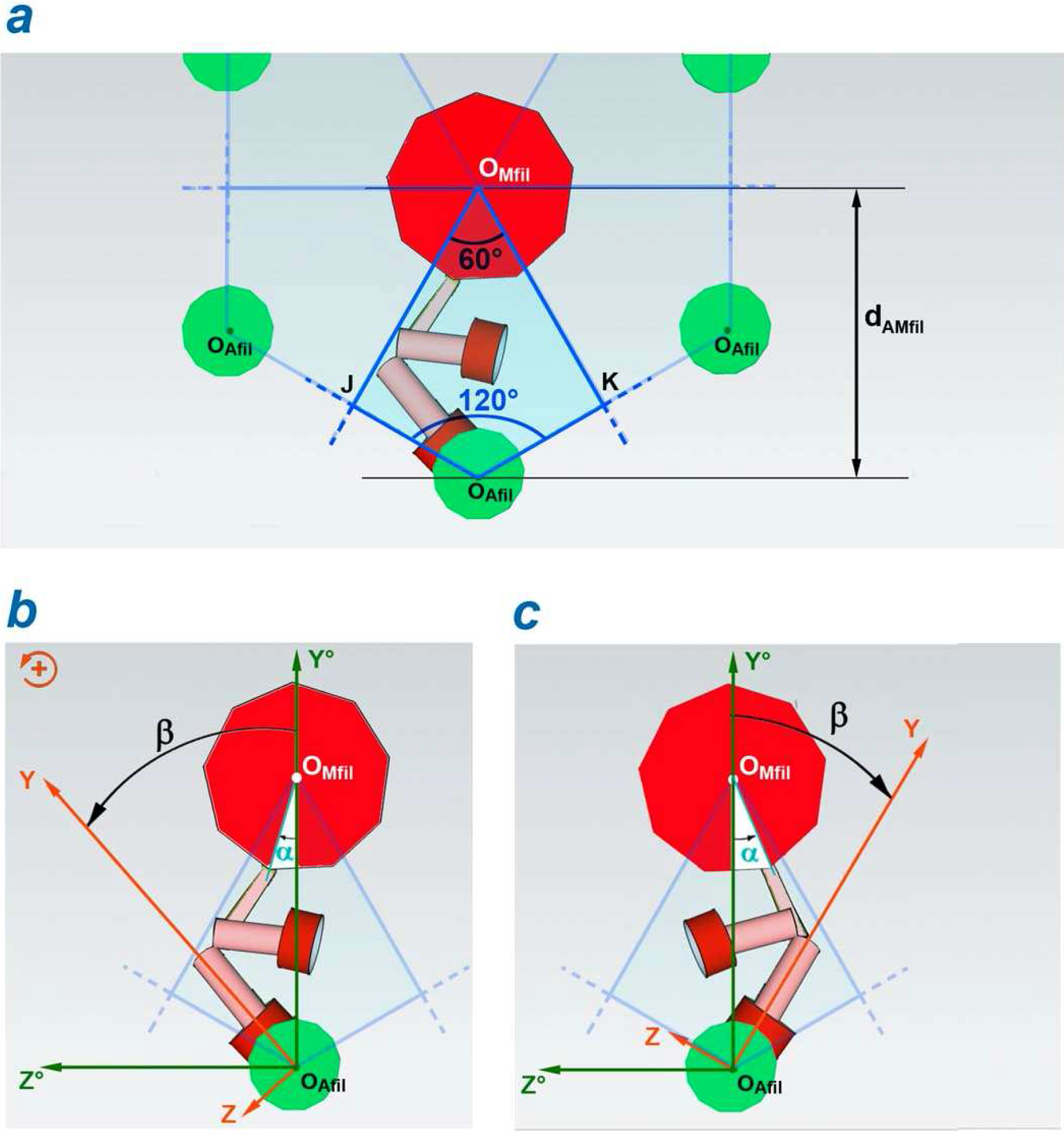
Angles α and β determined relatively to the O_Afil_, O_Mfil_ diagonal of the kite associated with the myosin molecule of which one of the 2 myosin heads is in WS. (a) Quadrilateral in the shape of a kite associated with a Mrow. (b) Case 1 where the WS head is located on the left relative to O_Afil_O_Mfil_ with −30° ≤ α < 0° and 0° ≤ β < +45°. (c) Case 2 where the WS head is located on the right relative to O_Afil_O_Mfil_ with 0° ≤ α < +30° and −45° ≤ β < 0°.

The 2 heads of a Mmol cannot be in WS state simultaneously because they are strongly attached to 2 different Asit, it is impossible for their respective S1b levers to move in 2 fixed planes characterized by 2 different angles β which must each remain constant during the longitudinal displacement of the Afil relative to the Mfil.

Geometrically, the 2 heads of a Mmol play an equivalent role with respect to point A. To define which head is most likely to transit in the WS state, we predict two cases:

**Case 1** (Fig 2b): α negative leads to β positive and only the left most head can bind strongly:

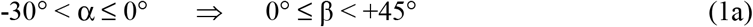

**Case 2** (Fig 2c): α positive leads to β negative and only the right most head can bind strongly:

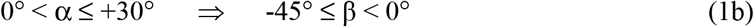

The 45° limit imposed on β in (1a) and (1b) comes from the condition imposed in (17) in accompanying Paper 2.

### Reminder of the geometric equations characterizing the cross-bridge of a WS head

In supplement S2.D of Paper 2, the abscissa of point A (X_A_) is calculated from the abscissa of point D (X_D_) on the basis of the 3 equations (D4), (D2) and (D3) reproduced below:

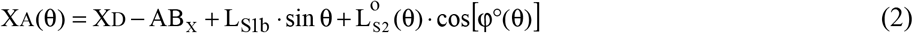

with

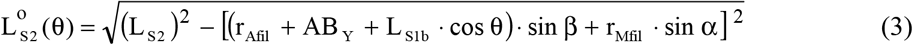

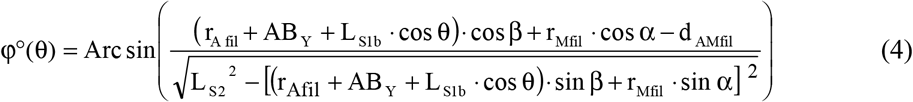

All the acronyms appearing in these 3 equations and the values assigned to them in the algorithmic calculations are listed in Table 1. When the angles α and β are fixed, the relationship (2) characterizes a bijection between the 2 variables X_A_ and θ. The knowledge of X_A_ determines the knowledge of θ and vice versa.

The linearization of expression (2) provides the relationship between hs displacement (ΔX) and lever rotation (Δθ) established in (19) in Paper 2 and duplicated below:

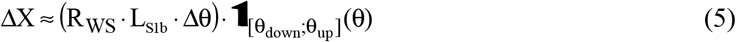

### Determination of the number of myosin heads potentially in WS

The length of a hs being fixed between 1 and 1.1 μm, the plane of the computer routine which makes it possible to calculate, on the one hand, the number of eventualities of WS heads belonging to a Mfil surrounded by six Afil and, on the other hand, the exact value of the angle θ associated with each of these eventualities, complies with the following algorithmic procedures. We precise that all acronyms in the form (Gx) where x is an integer refer to the equations in Supplement S3.G.

The six Afil surrounding a Mfil are immobilized (Fig 1d). The 360 Amol constituting an Afil are numbered from 1 to 360 in order of increasing abscissa on the O_Afil_X axis, the left edge of the first Amol serving as zero (Fig 1d). The point A representative of the Asit is located in the center of the associated Amol and the 360 abscissa of the 360 Asits are determined. A value of β_0_ is randomly selected between −13.85° and +13.85° according to (G9) and the angle β_C_ associated with each of the 360 Asit is calculated according to (G2). Then the angle β of the 360 Asit is evaluated relative to the diagonal of each of the six kites associated with the six Afil according to (G8). In the particular case where β_0_ = 0°, the results are delivered in column 4 of Table G2 for the first 13 Asit; the β values of the Asit numbered from 14 to 360 are deduced by iterative repetition. A value of α_0_ is randomly selected between −20° and +20° according to (G5). The angle α_C_ of each of the nine Mrow numbered from 1 to 9 by the index r is calculated using (G1). The equality (G3) gives the k-number of the kite where the Mrow n° r is located. The angle α associated with each of the nine Mrow is evaluated relative to the diagonal of the kite n° k according to (G4).

The abscissa of point D of the first Amol belonging to the Mrow n° 1 located in kite n° 1 (Fig 1c) is chosen arbitrarily over the length of the first Amotif (Fig 1d). The minimum and maximum abscissa of point A of the cross-bridge are determined by adding, respectively, the distances (L_S2_-L_S1b_+AB_X_) and (L_S2_+L_S1b_-AB_X_) to the abscissa of point D. Then we look for the numbers of Asit whose abscissa is between these two limits and whose value of β checks the inequalities (1a) or (1b) according to the sign of α. If the conditions are true, we calculate using equation (2) the two abscissa of the two points A of the cross-bridge corresponding to the two values, θ_down_ and θ_up_. It is recalled that the angles θ_down_ and θ_up_ are the boundaries of θ during the WS; their respective values are calculated in Paper 2 and displayed in Table 1. If the abscissa of the selected Asit is framed by these two new values, we count this possibility of WS and we calculate θ as a first approximation by linearly interpolating between θ_down_ and θ_up_ relative to the abscissa of this Asit. From this interpolated value, the exact value of θ is approached by iteration to an accuracy of 0.5° using equation (2). By loop the abscissa of the other points D of the Mrow n° 1 is calculated, each point D being spaced from the previous point D with a pitch equal to δX_Mmol_ (Table 1 and Fig 1d). The routine described for point D of the first Amol is applied successively to point D of the other 15 Amol of the Mrow n° 1. The r-index is incremented, r varying from 2 to 9 and the previous routine is replicated for each point D of the 16 Amol belonging to the Mrow n° r located in the associated kite n° k.

Then the Mfil is advanced by 1 nm compared to the six fixed Afil, a movement corresponding to a 1 nm shortening of the hs (Fig 1d). The algorithmic procedure described above is reproduced for each of 16 Amof of each of 9 Mrow of the Mfil. The hs shortening is iterated in steps of 1 nm up to a total distance of 11 nm, i.e. a value close to 11.5 nm, the stroke size of a myosin head (δX_Max_).

## Results

### Search for a probability density of the angle θ of the levers S1b belonging to the myosin heads potentially in WS

The geometric data entered into the computer routine are provided in the “Value” column of Table 1. In addition, the three following conditions are required:

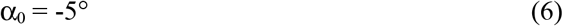

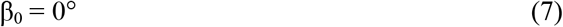

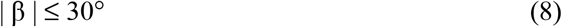

The position of point D of the first Mmol of the Mrow n° 1 of the Mfil is fixed at 34.5 nm relative to the first Amotif of the six Afil. Compared to this starting position, 11 shortenings of the hs are performed with a pitch of 1 nm, for a total length of 11 nm. To save space, only the even steps are presented in Figs 3 to 6. With the 3 conditions adopted in (6), (7) and (8), from one to four WS possibilities appear step by step according to the Mrow r-number, characterized by one to four occurrences of θ between θ_down_ and θ_up_. In the example in Fig 3, which refers to Mrow n° 5, there are two to four θ occurrences marked with a blue vertical line. The height of the blue line counts the number of Asit detected for the same value of θ; indexes of the Asit concerned is displayed above the blue vertical line. The difference between two successive indexes is a constant equal to 78, i.e. 6 times 13; see for example the series of three Asit (68, 146, 224) and (53, 131, 209) which appear in Figs 3b to 3f; the explanation comes from the fact that the conformation of the Mfil surrounded by the six Afil is identical after a longitudinal translation of six successive Amotif, a result that is interpreted with the data from Table 1:

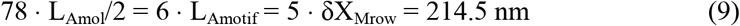

**Fig 3.**
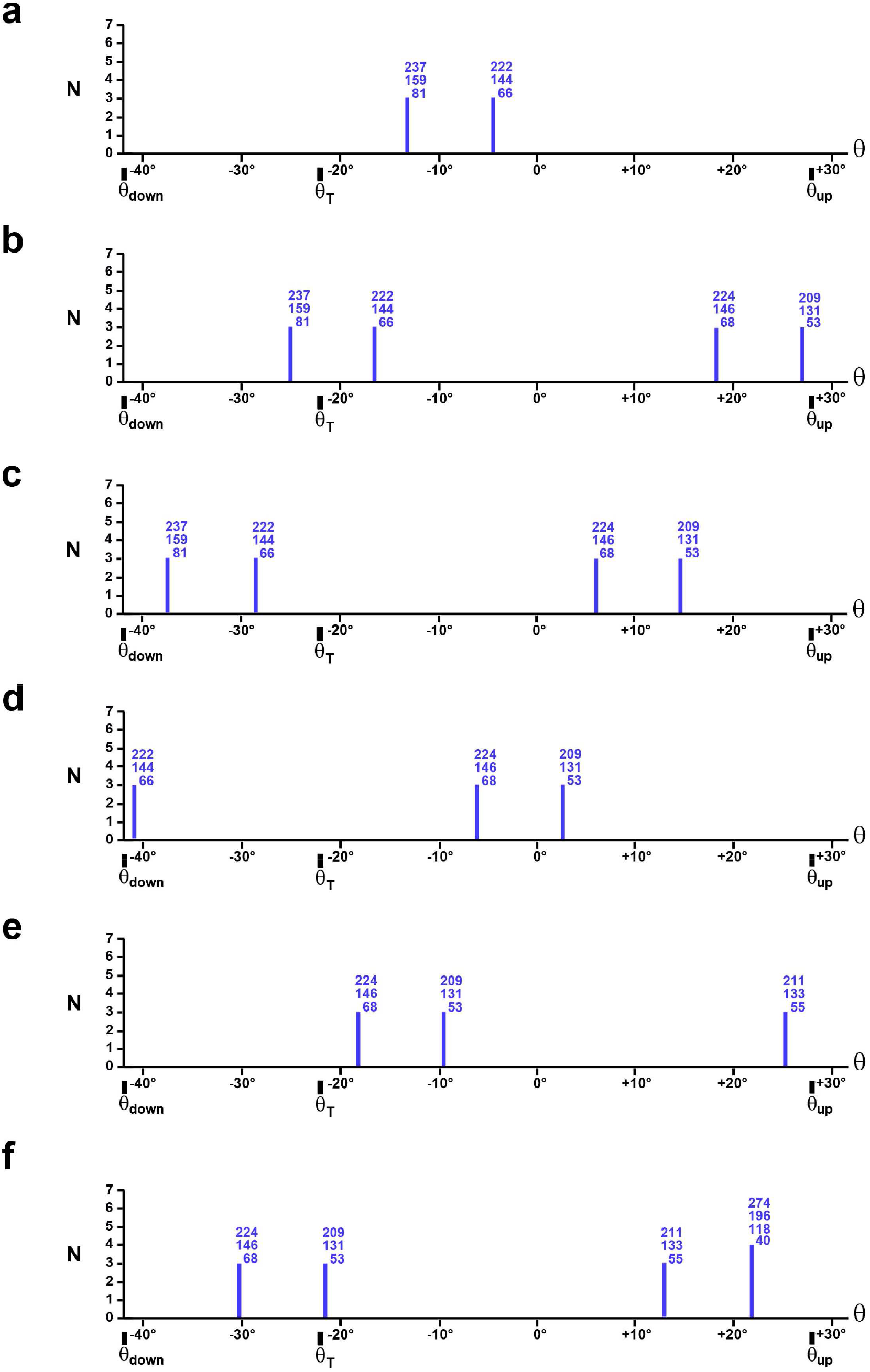
Numbers of WS possibilities for the 32 myosin heads of the n° 5 Mrow with the conditions: α_0_=−5°, β_0_=0° and |β|≤30°. a) The length of the hsR is fixed between 1 and 1.1 μm. (b), (c), (d), (e) and (f) After hsR shortening equal to 2, 4, 6, 8 and 10 nm, respectively.

The equalities present in (9) are reported on page 226 in [13].

The number of Asit for the same occurrence of θ is generally equal to 3 and rarely to 4 (Figs 3a to 3f) which is explained with equality (9) since:

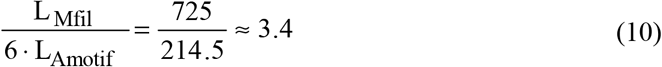

The previous findings are repeated for the other 8 Mrow of the Mfil. All the WS possibilities relating to the 288 heads of a Mfil with the conditions set out in (6), (7) and (8) are presented in each of the 6 graphs in Fig 4 associated with the even steps of the hs shortening. The angle θ is distributed by discrete values between the 2 terminals θ_down_ and θ_up_ relatively homogeneous way with the presence of one or two gaps of a width varying from 15° to 20°.

**Fig 4.**
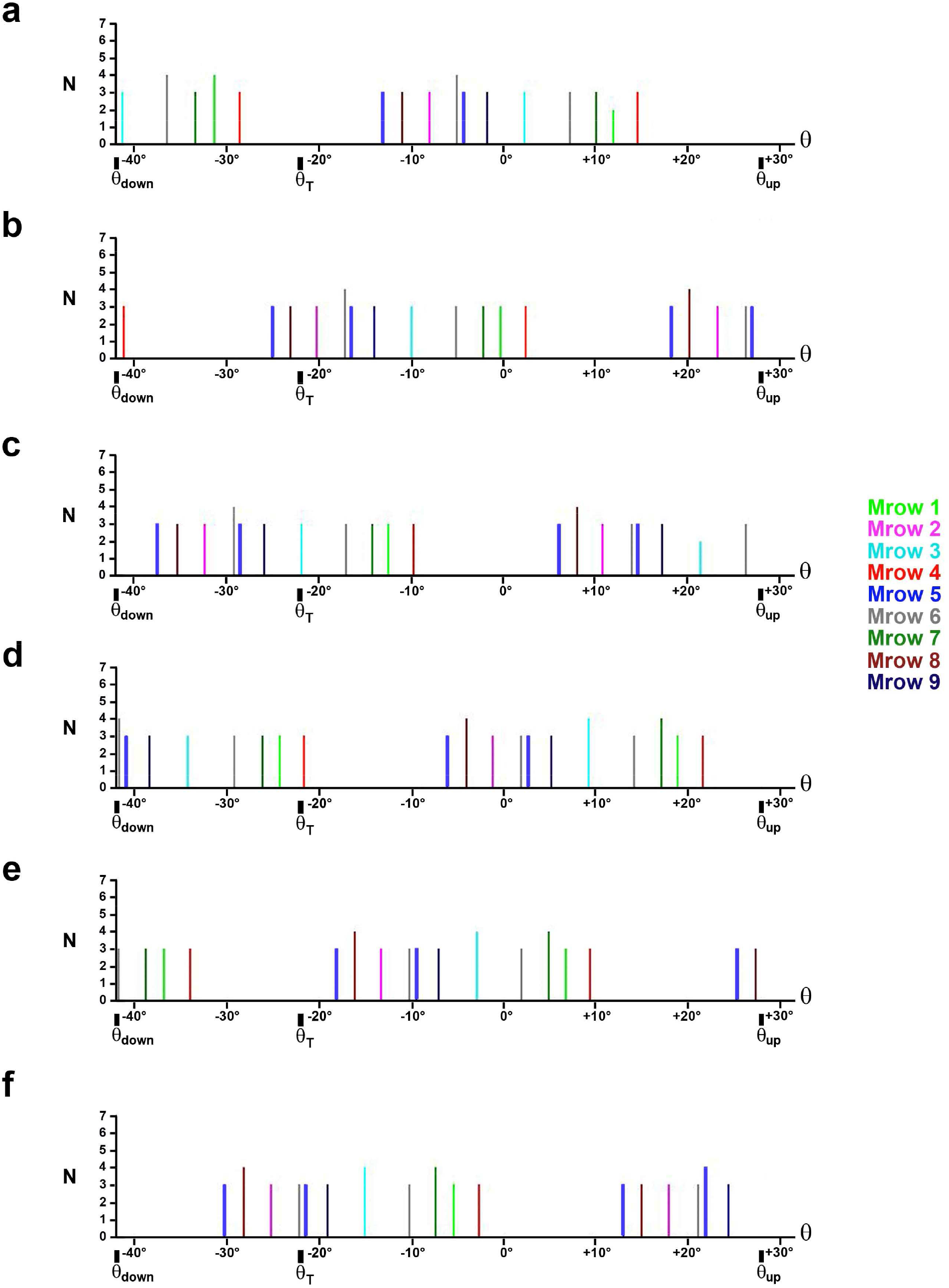
Numbers of WS possibilities for the 288 myosin heads belonging to the 9 Mrow of a Mfil with the conditions: α_0_=−5°, β_0_=0° and |β|≤30°. (a) The length of the hsR is fixed between 1 and 1.1 μm. (b), (c), (d), (e) and (f) After hsR shortening equal to 2, 4, 6, 8 and 10 nm, respectively. Each vertical line color refers to the color and therefore to the number of one of the 9 Mrow of the legend.

At this stage, the results relate to a Mfil surrounded by six Afil. A hs is composed of several hundred Mfil, from 400 Mfil to 2000 Mfil depending on the typology. In reality, the geometric values provided in Table 1 are not absolute constants but average data located in the middle of a confidence interval. Consequently, the θ values calculated on the basis of these data must *a minima* observe a variability. The integer number of occurrences of θ for the 9 Mrow is called as N_p_. Each of the N_p_ angles θ is indexed in ascending order between θ_down_ and θ_up_ using the discrete variable θi, the integer i varying from 1 to Np. In the examples in Fig 4, N_p_ is an integer number that varies between 15 and 20 depending on the step. The variability of θ is objectivized by assigning to each θi a normal probability density of mean (θi) and common standard deviation (σ_θ_) whose angular value corresponds in our model to a hs shortening (ΔX) equal to 1 nm. The relationship (5) gives:

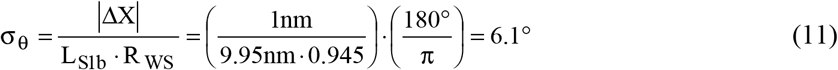

It remains to be determined the probabilistic weight to be given to each of the N_p_ values of θi. Indeed, the algorithmic procedure lists all cases of WS, and in particular those relating to the fixation of a myosin head on several neighbouring Asit on the same Ahel; see as an example, the duos of Asit numbered (66 and 68), (144 and 146), (222 and 224) on Figs 5b and 5c, or the duos numbered (53 and 55), (131 and 133), (209 and 211) on Figs 5e and 5f However, between the two eventualities mentioned, only one case is possible.

**Fig 5.**
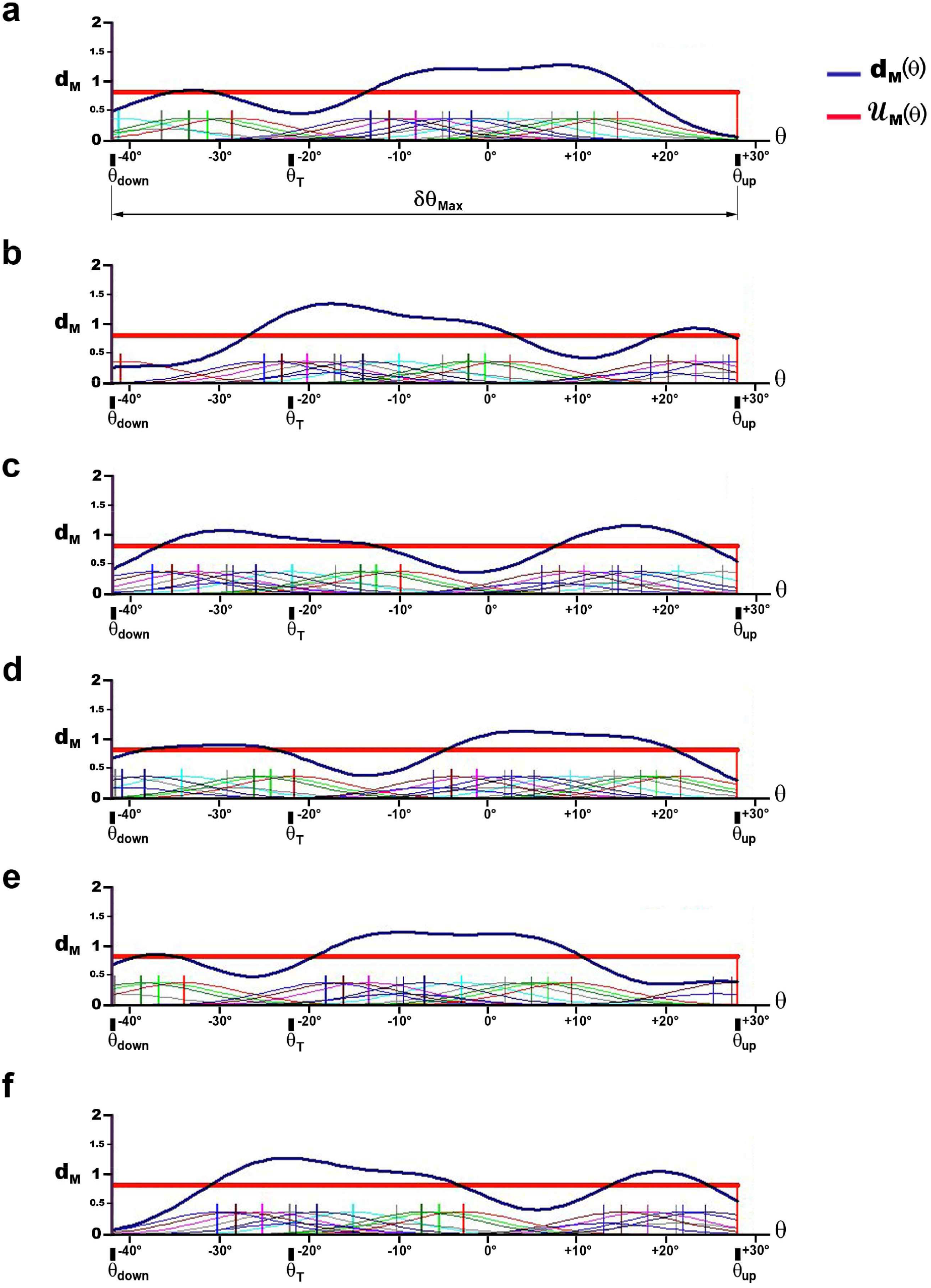
Probability densities d_G_ with conditions: α_0_=−5°, β_0_=0° and |β|≤30°. (a) The length of the hsR is fixed between 1 and 1.1 μm. (b), (c), (d), (e) and (f) After hsR shortening equal to 2, 4, 6, 8 and 10 nm, respectively.

For each hs shortening step, a function (f_M_) is defined as equal to the sum of the N_p_ Gaussian laws centred on one of the value of θi. Each Gaussian law is restricted to the interval [θ_down_; θ_up_] and is weighted by a probabilistic coefficient (p_i_). The function f_M_ is formulated as follows:

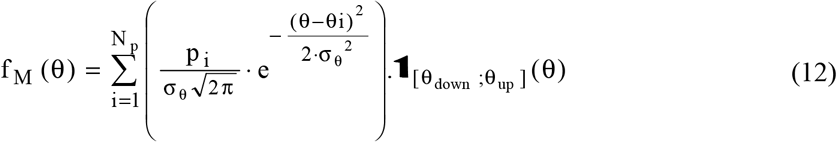

where N_p_ is the number of occurrences of θi corresponding to the shortening studied; pi is equal to 1, 1/2 or 1/3 depending on whether there are 1, 2 or 3 fixing possibilities for a WS head on 1, 2 or 3 adjacent Asit on the same Ahel; σ_θ_ is the common standard deviation equal to 6.1° according to (11); **1** is the indicator function presented in (A2b) in Supplement S1.A of Paper 1.

The probability density (d_M_) is introduced by normalizing f_M_:

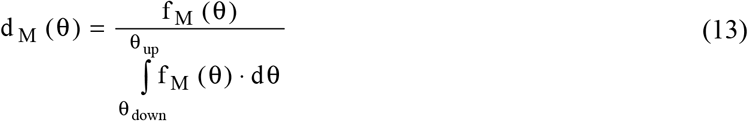

The calculations of the densities enounced in (12) and (13) are performed by computer programming for the reference position and then for the 11 hs shortenings. The plots of the N_p_ Gaussian laws appear in Figs 5a to 5f for even shortening steps where the height of the Gaussians is artificially increased by a factor of 6 for readability reasons. The probability density d_M_ is represented by a solid blue line on the 6 graphs in Fig 5. Among the classical statistical laws of a continuous random variable, the most convincing law for approaching the 12 densities d_G_ is the uniform law 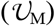 which is written on the interval [θ_down_; θ_up_]:

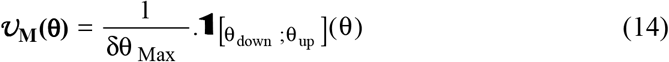

where 1/δθ_Max_=0.8185 with δθ_Max_=70° (Table 1), the calculation being carried out in radian.

The 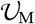 uniform law given in (I4) is represented by a red solid line (Figs 5a to 5f). The number of WS heads depends on many factors: calcium level, temperature, presence of an inhibitor, pH, etc. This means that the conditions tested in this paragraph are subject to change. To this end, we have examined in Supplement S3.H conditions optimizing the number of WS possibilities, and the conclusions issued are identical.

### Hypothesis 5 of θ uniformity

From hypothesis 4 relating to the geometrization of the WS state and with the contribution of the previous results, the following conjecture results: if a muscle fiber is isometrically stimulated and if the length of each hs is between 1 and 1.1 μm, then the distribution of the angular positions θ of the levers belonging to the myosin heads in WS follows the 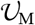 uniform law on [θ_down_; θ_up_] in each hs.

### Corollary of the homogeneity of the uniform law

The assumption of uniformity applies to any interval included in [θ_down_; θ_up_]. We can thus define a uniform law 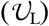 which is formulated on the interval [θ_2_; θ_1_] included in the interval [θ_down_; θ_up_]:

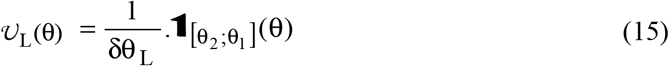

where θ_1_ et θ_2_ check θ_down_ ≤ θ_2_ < θ_1_ ≤ θ_up_ in a hsR and θ_up_ ≤ θ_1_ < θ_2_ ≤ θ_down_ in a hsL; δθ_L_ = |θ_2_ - θ_1_|.

### Application of the corollary

In the introduction, it was mentioned that during the isometric tetanus plateau, two models relating to the distribution of the angle θ, one uniform, the other Gaussian, were proposed in the literature over a range δθ_T_ between 40° and 50°. Our own model has recorded a value of δθ_T_ equal to 49°. In a hs on the right, we define the angle θ_T_ (T as Tetanus):

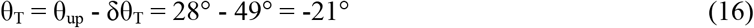

Among the N_p_ occurrences between θ_down_ and θ_up_, only the values of θi between θ_T_ and θ_up_ are used, the number of which is equal to N_T_ (Fig 4). The variability of θ is explained by assigning to the NT θi a normal probability density of mean (θi) and common standard deviation (σ_θ_). We define the function (f_T_) equal to the sum of the N_T_ Gaussian laws restricted to the interval [θ_T_; θ_up_], weighted by a probabilistic coefficient (p_i_), that is:

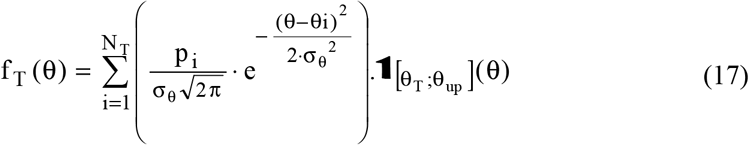

where σ_θ_ is a common standard deviation equal to 6.1° for the same reasons given in (12).

The probability density d_T_(θ) is calculated by normalizing f_T_(θ):

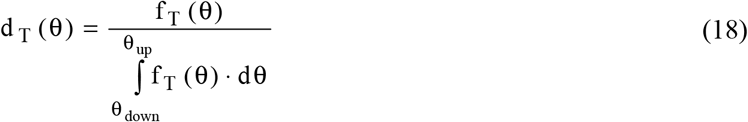

The calculations of (17) and (18) are performed by computer programming for the reference position and then for the 11 hs shortenings. The probability density d_T_ is represented by a solid blue line (Figs 6a à 6f). In accordance with experimental observations, we tested the uniform and Gaussian models.

**Fig 6.**
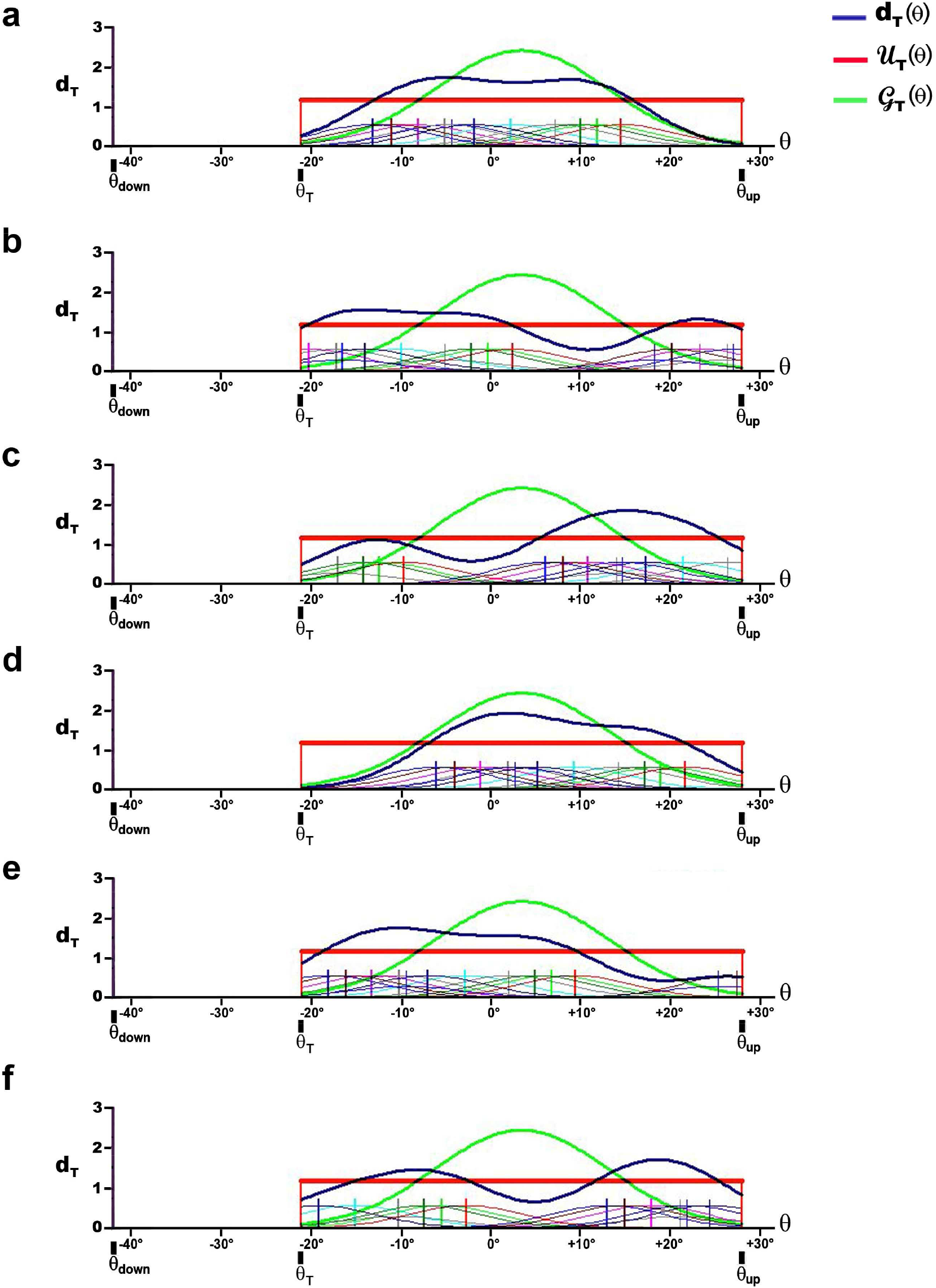
Probability densities d_T_ with conditions: α_0_=−5°, β_0_=0° and |β|≤30°. (a) The length of the hs on the right is fixed between 1 and 1.1 μm. (b), (c), (d), (e) and (f) After hsR shortening equal to 2, 4, 6, 8 and 10 nm, respectively.

The Uniform law 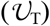 is formulated on the interval [θ_T_; θ_up_]:

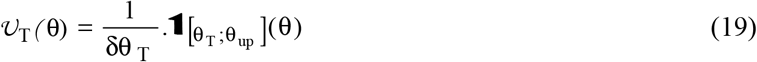

where 1/δθ_T_ = 1.1693 with δθ_T_ = 49°.

The Gaussian law 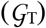 restricted to the interval [θ_T_; θ_up_] is formulated:

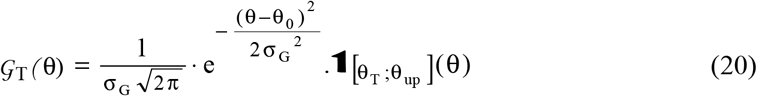

where θ_0_ is the mean on which the Gaussian is centred, i.e. the middle of the interval [θ_T_; θ_up_]:

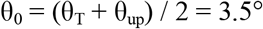

> σ_G_ is the standard deviation calculated so that the interval [θ_T_; θ_up_] contains at least 99% of the values so that 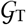 is representative of a probability density, that is:
>
> 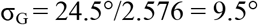

On the 6 graphs in Fig 6 corresponding to the reference position and the even shortenings, the uniform law is drawn with a thick red line and the Gaussian with a thick green line. By algorithmic procedure, the deviations 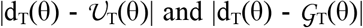 are summed for 100 θ values between θ_T_ and θ_up_ and incremented with the same pitch. The total of the deviations of the Gaussian model is twice as large as the total of the deviations of the uniform model. This result is found for the optimized conditions that are tested in supplement S3.H (Fig H5). Again the Uniform law prevails over other statistical laws and hypothesis 5 applies to any interval between θ_up_ and θ_down_.

## Discussion

### Data variability and model stability

A battery of tests was conducted using other values assigned to the model parameters provided in Table 1, variables used in algorithmic procedures such as angles α_0_ and β_0_, the starting position of the Mfil with respect to Afil, the inter-filament distance, the lengths of S1b and S2, the position of point B with respect to Asit, etc. Each of the trials concluded with results similar to those mentioned above.

The model is based on different assumptions: 1/ all Afil are identical, 2/ all Mfil are identical, 3/ the geometric parameters of a vertebrate hs remain constant during the experiments, i.e. the elongations, twists and other spatial deformations of the constituent elements of the hs are neglected, 4/ the inter-filament distance is fixed, 5/ the equality (9) which is at the origin of the discrete and repetitive nature of the angular occurrences θi is supposed to be verified at all times. To these objections, we answer that any modification that corrupts our modeling of the hs architecture *ipso facto* leads to an increase in the number N_p_ of angular positions θi, which results mathematically in a uniformization of the density of the discrete random variable θ between the 2 terminals θ_down_ and θ_up_. The differences between densities d_M_ and 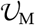 on the one hand, and d_T_ and 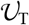 on the other hand, tend to decrease, thus supporting hypothesis 5.

### Uniformity of distribution during the isometric tetanus plateau

The corollary of hypothesis 5 specifies that uniformity applies to any angular interval between θ_up_ and θ_down_ and therefore to the interval δθ_T_ between θ_up_ and θ_T_ (Fig 5), an assumption in line with several experimental observations [9,12]. It should be noted that the Gaussian competing model is often used incorrectly because the standard deviation σG introduced in the law is equal to or greater than δθ_T_/2; the 99% confidence interval becomes in this case: [−2.576·δθ_T_/2; +2.576·δθ_T_/2], i.e. more than twice the extent of the announced interval.

The uniformity hypothesis has another consequence: if the angle θ is considered as a discrete random variable, then the number of heads is identical in all hs of the fiber maximally tetanized in isometric conditions. This outcome will be demonstrated by the laws of classical mechanics in accompanying Paper 4. This result is implicitly accepted by other researchers [3,10,15,16].

### The assumption of uniformity remains valid with hs of different lengths

The hypothesis is formulated from 11 shortenings with a total length of 11 nm from an initial position of the Mfil with respect to the Afil, position given equal to 34.5 nm. Other values within the range] 0; L_Amotif_] have been tested and the conclusion remains unchanged. We deduce that some hs can have the same uniform law while having a different length, varying between 1 and 1.1 μm. Hypothesis 5 provides an explanation for the non-uniformity of sarcomere length observed during the isometric tetanus plateau [1,2].

This result explains why, after a perturbation by a length step, the tension always rises to a value almost equal to that of the reference tetanus isometric tension (T0), because in the end we find the same uniform law on δθ_T_ which was present in all hs during the plateau preceding the disturbance (see accompanying Paper 5). It should be recalled that the absence of WS heads for θ between θ_down_ and θ_T_ during the isometric tetanus plateau is due to the slowly detachment, an event studied in Supplement S1.B of Paper 1.

### The number of WS heads varies according to the length of the hs

The probability densities d_G_ were determined as functions of the θi occurrences but without taking into account the repetition of θi according to equality (10); consequently the θ uniformity hypothesis extends to any Afil/Mfil overlap area between 0 and L_Mfil_=725 nm. It appears that the number of myosin heads in WS must decrease linearly with the length of the overlap area, i.e. with the length of the sarcomere when it is between 2.2 and 3.3 μm.

In accompanying Paper 4, the tension of the isometric tetanus plateau (T0) is determined. T0 is among other things a linear function of the number of myosin heads in WS per hs (Λ_0_). T0 must therefore decrease linearly with L0_s_ between 2.2 and 3.3 μm; this result is verified experimentally [3].

Conversely, since it is shown that during the isometric tetanus plateau, the number (Λ_0_) and distribution of the WS heads are identical in each hs and since experimentally T0 and L0_s_ (2.2μm≤L0_s_≤3.3μm), on the one hand, and T0 and Λ_0_, on the other hand, are linearly related, then Λ_0_ and L0_s_, are linearly related too, and the only statistical law to respond to this theoretical scheme is the Uniform law. This is another way to validate hypothesis 5 of uniformity.

### Number of WS per hs

The percentage of WS heads (pWS) during the isometric tetanus plateau is given as less than 20% for some authors [5,17,18], equal to or less than 43% for others [19]. Over the past ten years, there has been an apparent consensus around the value of 30% [10,16,20,21]. It is noteworthy that these figures refer to experiments carried out at low temperatures between 0°C and 6°C and that it is never clear whether the percentage is calculated on the total number of myosin molecules (144 per hs) or on the total number of myosin heads (288 per hs).

The tension measured during the tetanus plateau of a stimulated isometrically fiber doubles when the experimental temperature increases from 0°C to 30°C [22,23]. Since the tension of the stimulated fiber is assumed to depend on the number of WS heads [10,16], it is logical to assume that there are about twice as many heads at high temperature as at low temperature.

For these purposes we have calculated in Paragraph H.3 of Supplement S3.H several theoretical percentages during the isometric tetanus plateau over the interval δθ_T_ (pWS_T_) equal for one to 22% with Laplace equality and for the other to 25% (average at the bottom of the column “optimized conditions” of Table H2). We note that we obtain a percentage of 13%, which is half the average for the “normal conditions” column (Table H2).

## Conclusion

In our model, the WS is considered as a purely mechanical state (Fig B1 of Supplement S1.B to Paper 1) but the preparation of the WS has an entropic component. The first manifestation of the entropic nature of muscle contraction is found in the uniform and therefore random distribution of the orientation of levers S1b formulated with hypothesis 5.

## Supporting information

Data relating to computer programs used for Paper 3

Supplementary Chapter. Optimized conditions..

Computer programs for calculating and plotting densities of Teta angle

## Supporting information

### S3.G Supplementary Chapter. Calculation of the angles α and β

G. 1 Idealized geometric description of a vertebrate half-sarcomere with Figs G1 and G2, Table G1

G.2 Calculation of the index k, kite number where the Mrow n° r is located with Fig G3

G. 3 Determination of angles α and β with Fig G4 and Table G2

References of Supplement S3.G

### S3.H Supplementary Chapter. Optimized conditions

H.1 Probability densities of θ under optimized conditions on δθ_Max_ with Figs H1, H2 and H3

H.2 Probability densities of θ under optimized conditions on δθ_T_ with Figs H4 and H5

H.3 Calculation of the maximum percentage of myosin heads in WS with Tables H1 and H2

**CP3 Supplementary Material. Computer programs for calculating and plotting densities of θ angle.** Algorithms are written in Visual Basic 6.

**DA3 Supplementary Material. Data relating to computer programs used for Paper 3.** Access Tables are transferred to Excel sheets.

