## Supplemental Data 1 for "Mechanical model of muscle contraction. 3. The orientation of the levers belonging to the myosin heads in working stroke follows the same uniform law in all half-sarcomeres of an isometrically stimulated fiber"

### S3.G Supplementary Chapter of Paper 3

### Calculation of angles α and β

The architecture of a vertebrate half-sarcomere (hs) presented here conforms to the classical descriptions [1,2,3,4]. The length of the sarcomere is assumed to vary over the length range where the inter-filament overlap area of the myosin heads is maximum [5.6].As a majority of experiments are performed with frog fibers, we retain bounds 2 and 2.2 µm for the reference length range of an individual sarcomere [5].
The definitions of the acronyms appearing in the text are specified in Table G1.

#### G.1 Geometric description of an idealized half-sarcomere

In the cut plane of a hs represented by the transverse plane $O_{Mfil}Y°Z°$ (Fig G1), a myosin filament (Mfil) is patterned by a red circle. Each Mfil is located in the center of a regular hexagon, each of the six vertices of which is occupied by the center of an adjacent Mfil. The hexagon is formed by six equilateral triangles (Fig G1; purple lines), each orthocenter coinciding with the center of an actin filament (Afil) modelled by a green circle. The combination of these six orthocentres forms a new regular hexagon (Fig G1; dotted green) which consists of six kite shaped quadrilaterals numbered from 1 to 6 in the trigonometric direction. The longitudinal axes of a Mfil and an Afil are parallel to $O_{Mfil}X$, the longitudinal axis of the hs, and perpendicular to the $O_{Mfil}Y°Z°$ plane (Fig G2b). A Mfil is made up of three-stranded helix (Mhel) forming 9 rows (Mrow) parallel to the longitudinal axis; the 9 Mrows are represented by 9 red radiating lines around each Mfil on Figs G1 and G2a, and by 9 vertical and parallel lines on Fig G2b. A myosin molecule is called as Mmol and each Mrow has 16 Mmol spaced from $\delta X_{Mmol}$ for a total of 144 Mmol per Mfil (Fig G2b and Table G2). In our idealized hs model, all Mfils are identical with the same orientation (Fig G1), i.e. each of the 9 Mrow of each Mfil has an identical $\alpha_C$ angle with respect to the $O_{Mfil}Y°$ reference axis.

The angle $\alpha_C$ is calculated according to:

$$\alpha_C(r) = \alpha_0 + (r-1) \cdot \alpha_{Mrow} \qquad \text{(GI)}$$

where r is the Mrow index which varies from 1 to 9; $\alpha_0$ is the value of α relative to Mrow n°1 located in kite n° 1 with $\alpha_0 \in [-\alpha_{Mrow}/2 ; +\alpha_{Mrow}/2[$; $\alpha_{Mrow}$ is the constant angle between 2 adjacent Mrow in the transverse plane $O_{Mfil}Y°Z°$ such that $\alpha_{Mrow}$ = 360°/9 = 40°; the angular zero corresponds to the $O_{Mfil}Y°$ axis in Fig G1.

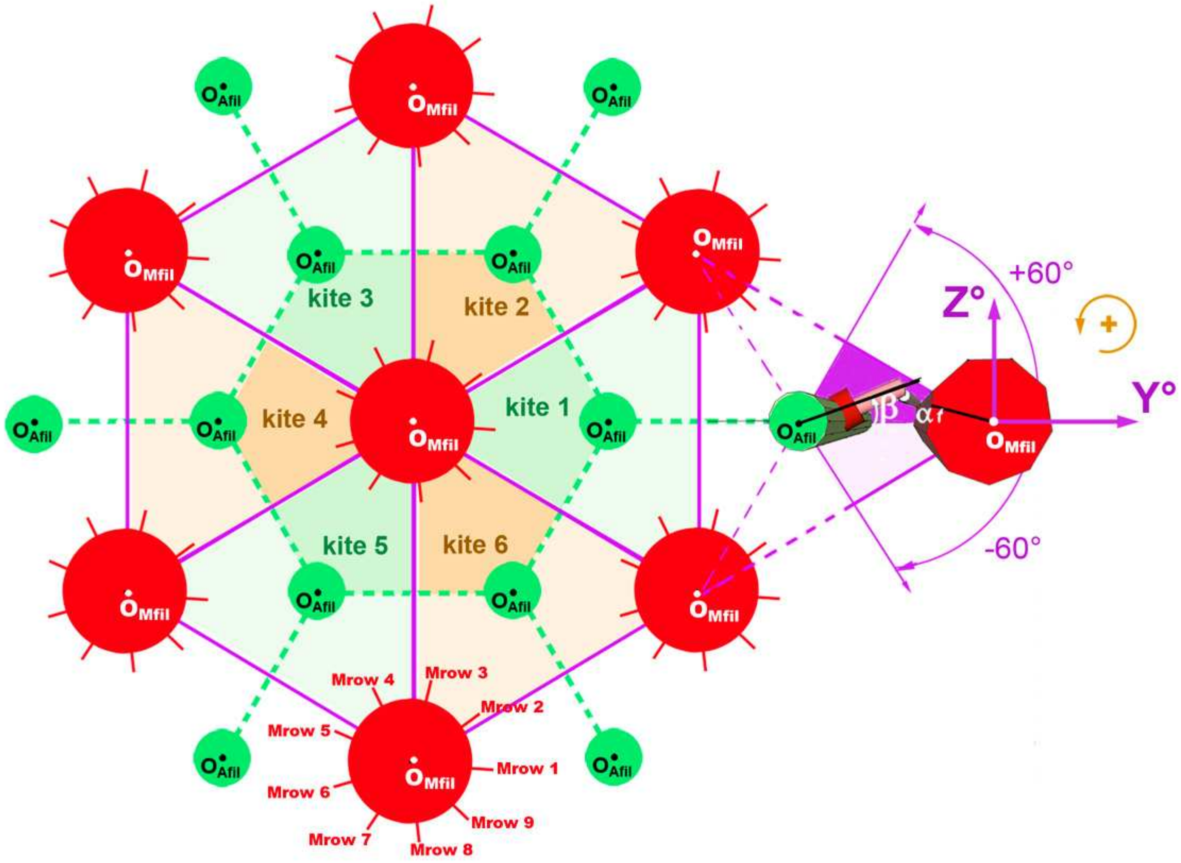


**Fig G1. Hexagonal lattices of actin and myosin filaments inter-digited in a vertebrate half-sarcomere.**

**Table G1**: **Geometric and numerical data characteristic of a myosin filament (Mfil) and an actin filament (Afil) in a half-sarcomere of skeletal muscle; values from [2].**

| | Acronym | Definition | Value | Formula |
|---|---|---|---|---|
| **Mfil** | $L_{Mfil}$ | *Mfil length without the bare area* | 725 nm | |
| | $r_{Mfil}$ | *Mfil radius (outside bare area)* | 7.5 nm | |
| | Mhel | *Mfil helix* | | |
| | $N_{Mhel}$ | *Number of coaxial Mhel of the Mfil* | 3 | |
| | $p_{Mhel}$ | *Mhel pitch* | 128.7 nm | $= p_{Mmol} \cdot N_{Mrow}$ |
| | $p_{Mmol}$ | *According to $O_{Mfil}X$, pitch between 2 successive Mmol belonging to the same Mhel* | 14.3 nm | $= p_{Mhel} / N_{Mrow}$ |
| | Mrow | *Row of 16 Mmol linked by S2 to a Mfil over the same longitude parallel to the $O_{Mfil}X$ axis* | | |
| | $N_{Mrow}$ | *Mmol number per turn of a Mhel*<br>*Mrow number per Mfil* | 9 | |
| | $\alpha_{Mrow}$ | *Angle between 2 adjacent Mrow in the transverse plane $O_{Mfil}X°Y°$* | 40° | $= 360° / N_{Mrow}$ |
| | $\delta X_{Mmol}$ | *According to $O_{Mfil}X$, distance between 2 adjacent Mmol in the same Mrow* | 42.9 nm | $= p_{Mhel} / N_{Mhel}$<br>$= p_{Mmol} \cdot N_{Mhel}$ |
| | $N_{Mmol,Mrow}$ | *Mmol number per Mrow* | 16 | $= L_{Mfil} / \delta X_{Mmol}$ |
| | $N_{Mmol}$ | *Mmol number per Mfil* | 144 | $= N_{Mmol,Mrow} \cdot N_{Mrow}$ |
| | $N_{S1}$ | *Number of myosin II heads per Mfil* | 288 | $= 2 \cdot N_{Mmol}$ |
| **Afil** | $L_{Afil}$ | *Afil length* | 1000 nm | |
| | $r_{Afil}$ | *Afil radius* | 3.5 nm | |
| | Ahel | *Interlaced helix of the Afil* | | |
| | $N_{Ahel}$ | *Ahel number* | 2 | |
| | Amotif | *Helical and iterative motif of the Afil formed by (13/6) Amol : 7 Amol belong to one of the Ahel and 6 to the other Ahel, then alternately from motif to motif in successive motifs* | | |
| | $N_{Amol,Amotif}$ | *Amol number per round of Ahel or per Amotif* | 13 | |
| | $L_{Amol}$ | *Amol length according to $O_{Afil}X$* | 5.5 nm | $L_{Amol}$ |
| | $L_{Amotif}$ | *Amotif length* | 35.75 nm | $= N_{Amol,Amotif} \cdot L_{Amol} /2$ |
| | $p_{Ahel}$ | *Ahel pitch* | 71. 5 nm | $= N_{Amol,Amotif} \cdot L_{Amol}$ |
| | $\beta_{Asit}$ | *Angle between 2 successive Asit located on 2 Amol belonging to the same Ahel in the transverse plane $O_{Afil}Y°Z°$* | 27.7° | $= 360° / N_{Amol,Amotif}$ |
| | $N_{Amotif}$ | *Amotif number per Afil* | ~28 | $= L_{Afil} / L_{Amotif}$ |
| | $N_{Amol,Ahel}$ | *Amol number per Ahel* | ~180 | $= L_{Afil} / L_{Amol}$ |
| | $N_{Amol}$<br>$N_{Asit}$ | *Amol or Asit number per Afil* | ~360 | $= N_{Amol,Ahel} \cdot N_{Ahel}$<br>$= N_{Amol,Amotif} \cdot N_{Amotif}$ |

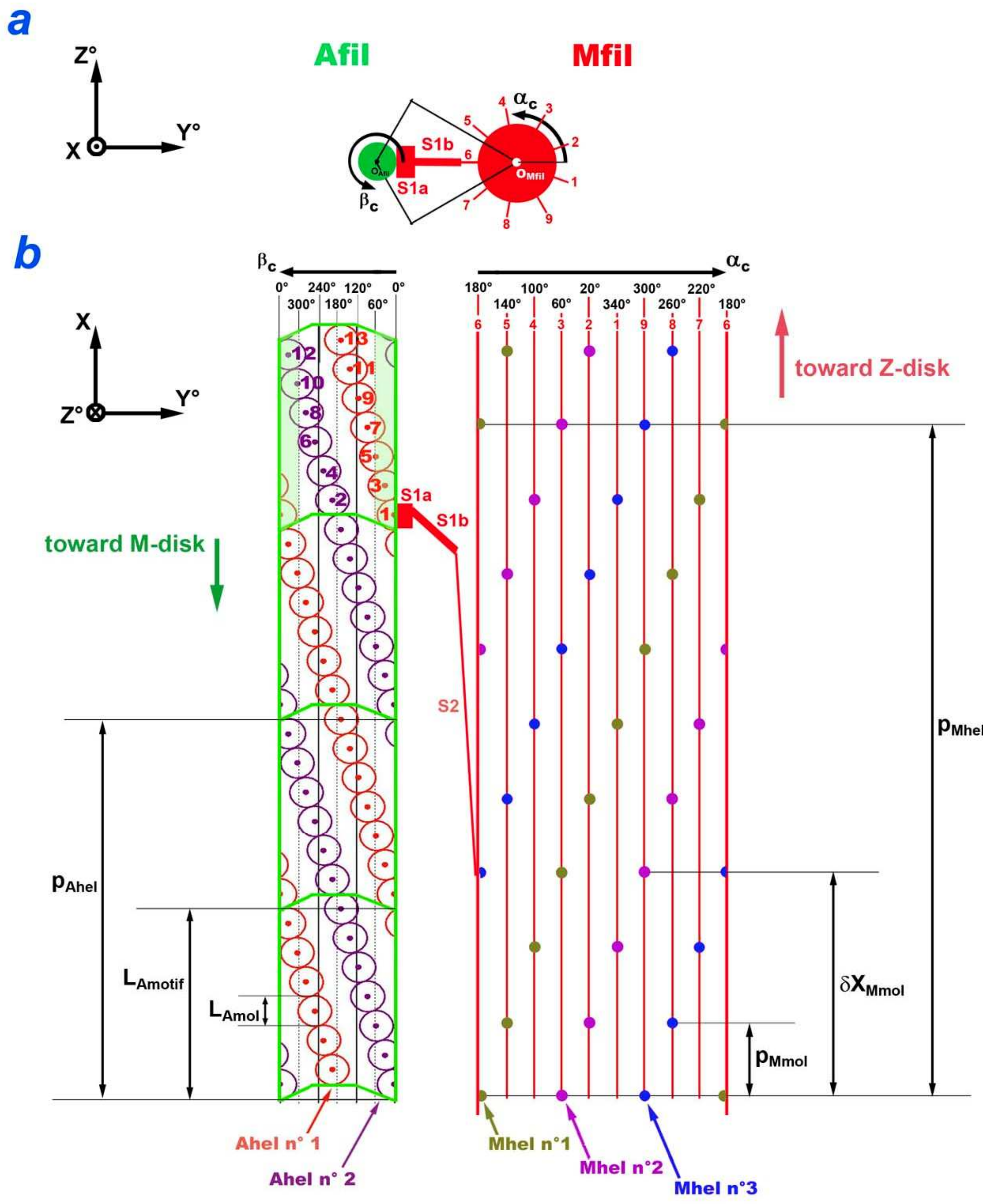


**Fig G2. A myosin filament (Mfil) and an actin filament (Afil) connected by a WS myosin head with $\alpha_0$ = -20° and $\beta_0$ = 0°.**

(a) Cutting plane in OY°Z° of the Mfil and Afil located in kite n° 4; the WS myosin head belongs to Mrow n° 6 and is strongly linked to Asit n° 1. (b) Mfil and Afil unwound in the OXY° plane. Each red or purple circle of the Afil represents an Amol. Each khaki, purple or blue dots on the Mfil correspond to the ball joints of the S2 rods of the Mmol.

An actin molecule is called as Amol. An Afil is made up of 2 intertwined helix (Fig G2b; Ahel 1 and Ahel 2). On an Afil, we observe the repetition (28 times) of the same motif delimited by a thick green line on Fig G2b, named as Amotif, consisting of (13/6) Amol [2,7]. In total, there are about 360 Amol per Afil (Table G1). On the surface of each Amol is a binding site for the myosin head, this site is called as Asi. For each Amotif there are 13 Asit numbered from 1 to 13 (Fig G2b).

In our idealized model, all Afils are identical with the same orientation (Fig G1), i.e. the 13 Asit of an Amotif have 13 constant $\beta_C$ angles relative to $O_{Afil}Y°$, formulated according to:

$$\beta_C(s) = \beta_0 + \left\lfloor \frac{(s-1) + 13 \cdot \mathbb{1}_{h=2}}{2} \right\rfloor \cdot \beta_{Asit} \quad \text{(G2)}$$

where s is the Asit index varying from 1 to 13 in order of increasing abscissa on the $O_{Afil}X$ axis (columns 1 and 3 of Table G2); $\beta_0 \in [\, -\beta_{Asit}/2 \ ; +\beta_{Asit}/2 \,[$ ; $\beta_{Asit}$ is the constant angle between 2 successive Asit on the same Ahel in the $O_{Mfil}Y°Z°$ tansverse plane such that $\beta_{Asit}$ = 360°/13 = 27.7°; h is the Ahel number equal to 1 or 2 (Table G2 ; column 2); $\mathbb{1}$ is the indicator function defined in (A2b) in Supplement S1.A of accompanying Paper 1.

The 13 values from $\beta_C$ appear in column 4 of Table G2 for the special case $\beta_0$=0°.

#### G.2 Calculation of the k-index of the kite where Mrow n° r is located

Each myosin head in working stroke (WS) occupies a kite whose 4 vertices are $O_{Mfil}$, $O_{Afil}$, J and K (Fig G3a). The two points J and K are the midpoints of the segments joining $O_{Afil}$ at the centers of the two adjacent Afil on the right and left, respectively. Regardless of the distance between $O_{Mfil}$ and $O_{Afil}$, the angle formed by the 2 segments $O_{Mfil}J$ and $O_{Mfil}K$ is 60° and the angle formed by the 2 segments $O_{Afil}J$ and $O_{Afil}K$ is 120° (Fig G3a).

The index of the kite in which the Mrow n° r is located is formulated:

$$k(r) = \text{int}\left(\frac{\alpha_C(r) + 30°}{60°}\right) + 1 \quad \text{(G3)}$$

where r is the Mrow index varying from 1 to 9; int is the integer part; $\alpha_C$( r) is the angle defined in (G1).

In the particular case where r=9 and $\alpha_C(9) \geq 330°$, the calculation in (G3) provides an index k equal to 7 which in fact corresponds to kite n° 1.

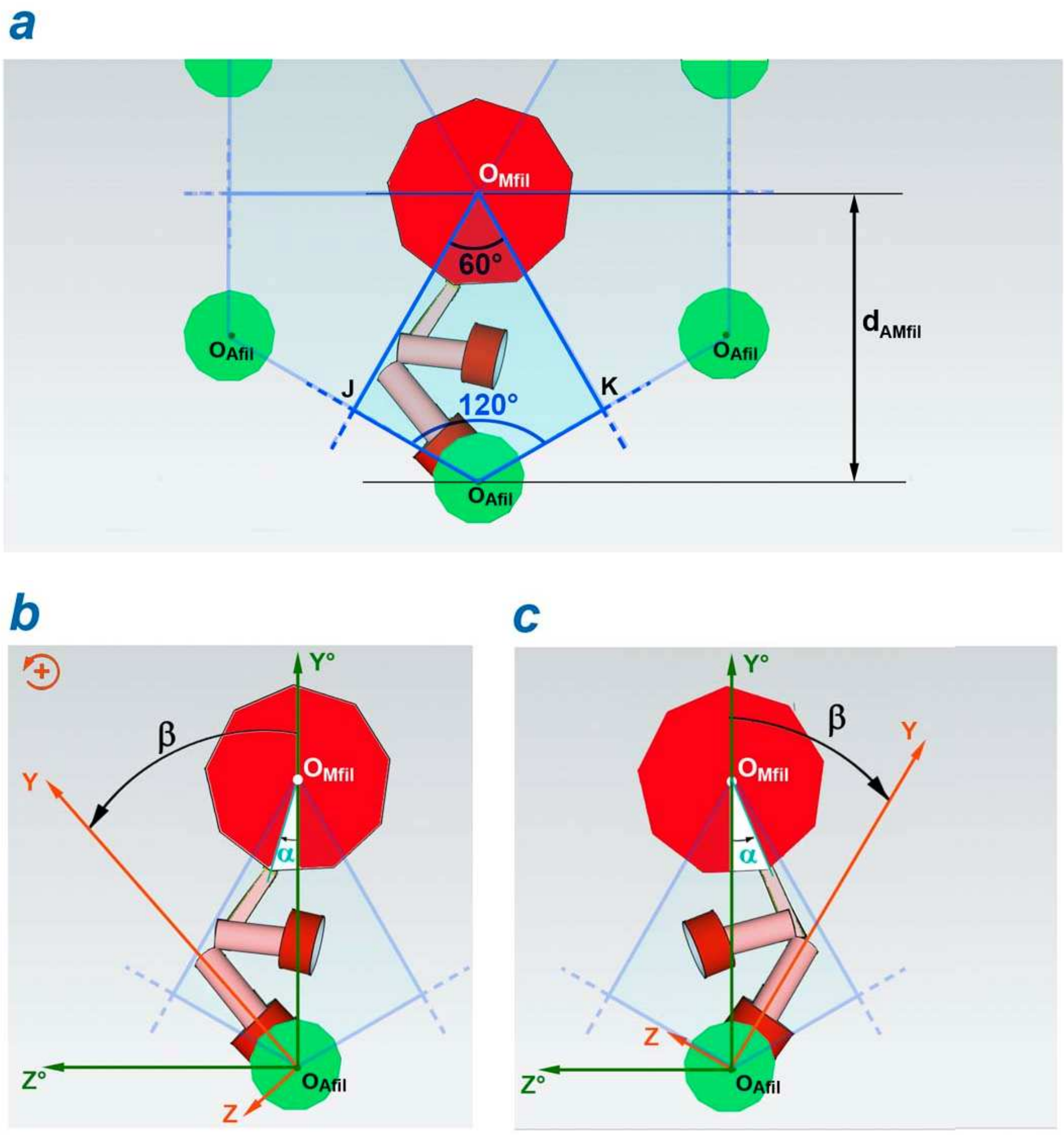


**Fig G3. Angles α and β expressed relative to the $O_{Afil}O_{Mfil}$ diagonal of the kite associated with the myosin molecule of which one of the 2 myosin heads is in WS.**

(a) Kite shaped quadrilateral associated with a Mrow. (b) Case 1 where the WS head is located on the left relative to $O_{Afil}O_{Mfil}$ with $-30° \leq \alpha < 0°$ and $0° \leq \beta < +60°$. (c) Case 2 where the WS head is located to the right relative to $O_{Afil}O_{Mfil}$ with $0° \leq \alpha < +30°$ and $-60° \leq \beta < 0°$.

#### G.3 Determination of angles α and β

The $O_{Mfil}O_{Afil}$ line is an axis of symmetry of the kite, which serves as a reference axis, both for the α angle of the Mrow to which the WS head belongs, and for the β angle of the Asit to which the motor domain S1a is strongly bound (Figs G1, G3b and G3c). The angles α and β are calculated with respect to the diagonal of the kite n° k where the Mrow n° r and the Asit n° s are positioned.

##### *G.3.1 Calculation of α*

The angle α corresponding to the diagonal of kite n° 1 is 0° and the diagonals of 2 adjacent kites are spaced 60° apart (Figs G1 and G3a). Thus the angle α of a WS myosin head belonging to Mrow n° r is calculated with respect to the diagonal of the kite n° k according to (G1):

$$\alpha(r) = \alpha_0 + (r-1)\cdot\alpha_{Mrow} - [k(r)-1]\cdot 60° \qquad \text{(G4)}$$

where k(r) is defined in (G3).

It is noted:

$$\forall\alpha_0 \in [-20;+20°[, \qquad \alpha \in [-30;+30°[ \qquad \text{(G5)}$$

##### *G.4.1 Calculation of β*

The WS head belonging to the Mrow n° r is located in one of the 6 kites of index k. Each kite has for common summit $O_{Mfil}$, the center of the Mfil, and for opposite summit one of the 6 centers of the 6 Afil surrounding the Mfil (Figs G1 and G4). The $\beta_C$ angle of the Asit to which the head is attached belongs to the green-grey zone specific to each of the 6 Afil (Fig G4). This area is between two terminals shown in Fig G4. The integer (m) is defined such that:

$$m = (k + 2) \cdot \text{mod } 6 \qquad \text{(G6)}$$

where k is the kite index calculated in (G3).

In support of Fig G4, the $\beta_C$ angle of the Asit n° s located in the kite n° k must check the following conditions:

$$[(m - 1) \cdot 60°] \leq \beta_C(s) < [(m + 1) \cdot 60°] \qquad \text{(G7)}$$

Since the angle of the diagonal of each kite is in the middle of the interval defined by the 2 terminals given in (G7), the $\beta_C$ angle corresponding to this diagonal is equal to (m·60°). We deduce that the angle β of a WS head belonging to the Mrow n° r is calculated with respect to the diagonal of the kite n° k according to (G2) with (G6) and (G7):

$$\beta(s,k) = \beta_0 + \left\lfloor \frac{(s-1) + 13\cdot\mathbf{1}_{h=2}}{2} \right\rfloor \cdot 27.7° - [(k+2)\text{ mod } 6]\cdot 60° \qquad \text{(G8)}$$

It is noted:

$$\forall\ \beta_0 \in [-13.85;+13.85°[, \qquad \beta(s,k) \in [-60;+60°[ \qquad \text{(G9)}$$

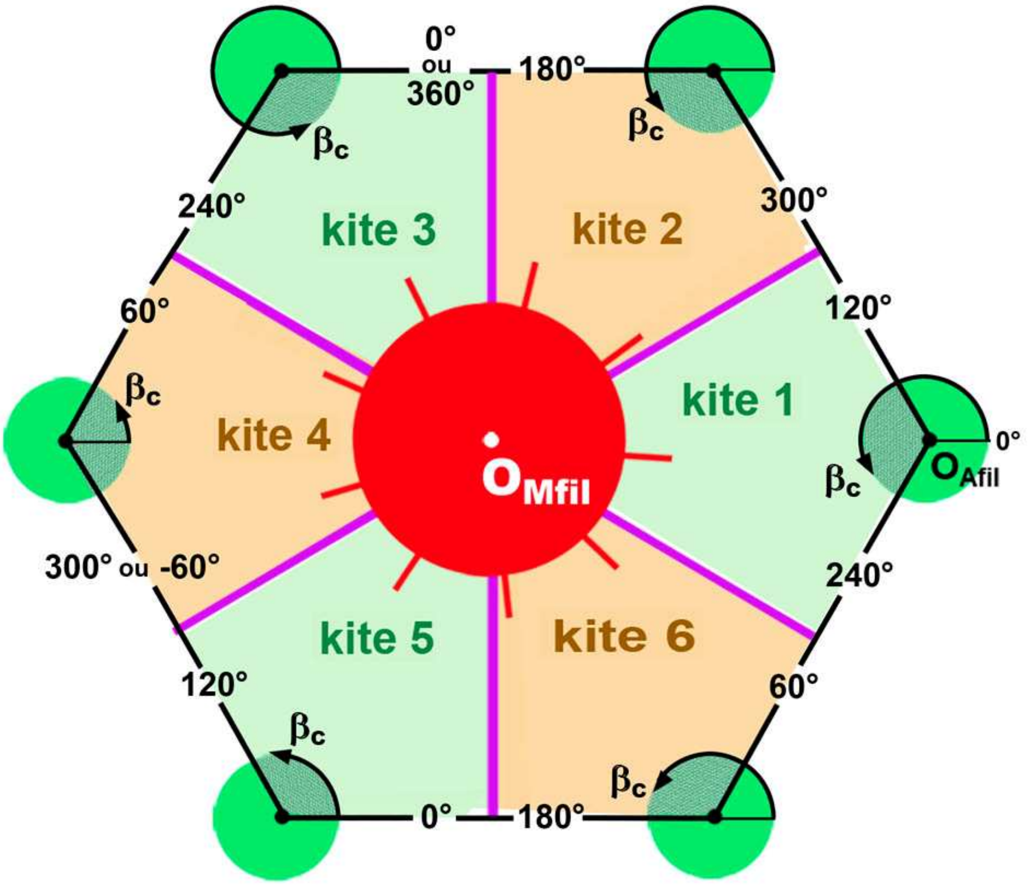


**Fig G4. Minimum and maximum terminals of $\beta_C$ in the 6 kites surrounding a Mfil.**

**Table G2**: **Geometric characteristics of the 13 Asit of an Amotif with $\beta_0$=0°**

| Asit | Ahel | $X_{Asit}$ | $\beta_C(s)$ | $\beta(s,k)$ | | | | | |
|---|---|---|---|---|---|---|---|---|---|
| | | (nm) | (°) | (°) | | | | | |
| s | h | | | kite 1 | kite 2 | kite 3 | kite 4 | kite 5 | kite 6 |
| 1 | 1 | I.75 | 0 | | | | 0 | -60 | |
| 2 | 2 | 5.5 | 193.8 | +13.9 | -46.2 | | | | |
| 3 | 1 | 8.25 | 27.7 | | | | +27.7 | -3I.3 | |
| 4 | 2 | 11 | 221.5 | +41.5 | -18.5 | | | | |
| 5 | 1 | 13.75 | 55.4 | | | | +55.4 | -4.6 | |
| 6 | 2 | 16.5 | 249.2 | | +9.2 | -50.8 | | | |
| 7 | 1 | 19.25 | 83.1 | | | | | +23.1 | -36.9 |
| 8 | 2 | 22 | 276.9 | | +36.9 | -23.1 | | | |
| 9 | 1 | 24.75 | 110.8 | | | | | +50.8 | -9.2 |
| 10 | 2 | 27.5 | 304.6 | | | +4.6 | -55.4 | | |
| 11 | 1 | 30.25 | 138.5 | -41.5 | | | | | +18.5 |
| 12 | 2 | 33 | 33I.3 | | | +3I.3 | -27.7 | | |
| 13 | 1 | 35.75 | 166.2 | -13.9 | | | | | +46.2 |

#### Special case with $\beta_0$=0°

In the example in Fig G2a, the WS head belongs to Mrow n° 6 and is located in kite n° 4. Only the Asit whose angle $\beta_C$ is between -60° and +60°, i.e. greater than 300° and less than 60°, are located in kite 4 (Fig G4), either according to column 4 of Table G2, the Asit n° 1, 3, 5, 10 and 12; see the two areas tinted pale green of the highest Amotif on the Afil in Fig G2b.

We applied the formula (G8) with $\beta_0$ nil. The 26 calculations of β related to the 6 kites are shown in the last 6 columns of Table G2.

In our model, if we apply the condition |β| ≤ 45° set out in accompanying Paper 2, then a WS myosin head has the possibility to attach itself to only 3 to 4 Asit at most among the 13 Asit of an Amotif.
