## Supplementary Chapter. Optimized conditions.. for "Mechanical model of muscle contraction. 3. The orientation of the levers belonging to the myosin heads in working stroke follows the same uniform law in all half-sarcomeres of an isometrically stimulated fiber"

### S3.H Supplementary Chapter of Paper 3

#### Optimized conditions

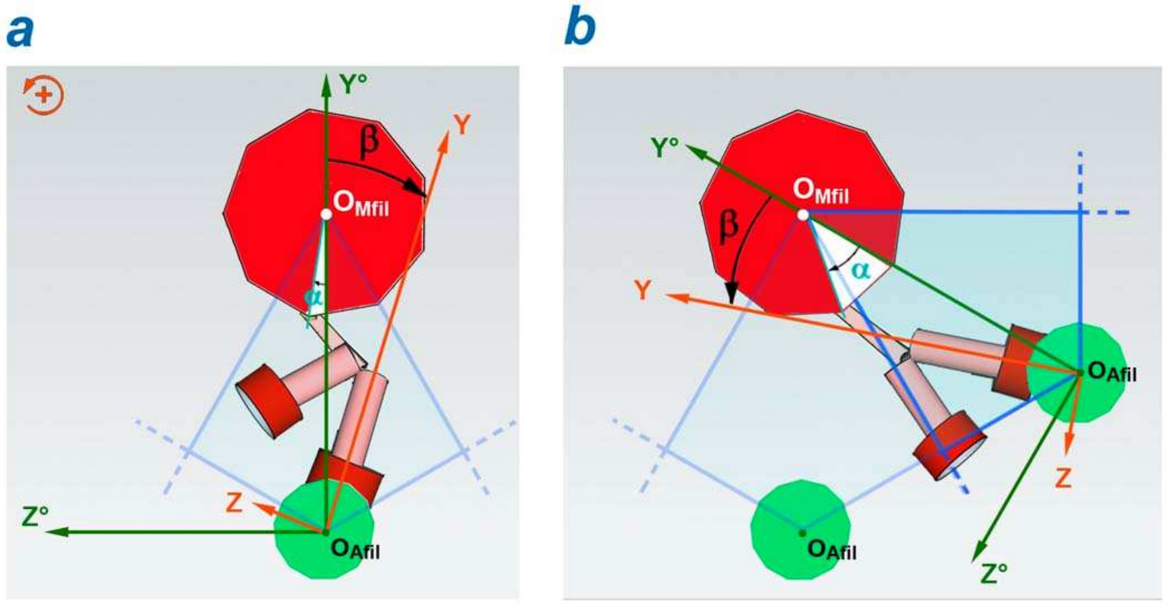


**Fig H1. Angles α and β expressed relative to the $O_{Afil}O_{Mfil}$ diagonal of the kite associated with the myosin molecule of which one of the 2 myosin heads is in WS.**

(a) Special case 1 where the WS head is located on the right relative to $O_{Afil}O_{Mfil}$ with 0 ≤ α ≤ +5° and 0° ≤ β < +45°. (c) Special case 2 where the WS head is located on the right relative to à $O_{Afil}O_{Mfil}$ with +25° ≤ α < +30° and 0° ≤ β < +45°, with β calculated in the adjacent kite on the right.

**H.1 Probability densities of θ under optimized conditions over the interval $\delta\theta_{Max}$**

Rules (1a) and (1b) set out in the Methods section of Paper 3 are amended in the following three special cases:

**1/** If α is close to 0°, then β can be positive or negative (Fig H1a):

$$-5° \le \alpha \le +5° \quad \Rightarrow \begin{cases} -45° \le \beta \le 0° \\ \text{or} \\ 0° \le \beta \le +45° \end{cases} \tag{H1}$$

**2/** Possibility for a WS head to be situated in the kites n° k or n° (k-1)

If α is close to the border between 2 kites, then a WS myosin head can be located in the adjacent kite:

$$-30° \le \alpha \le -25° \quad \Rightarrow \begin{cases} 0° \le \beta \le +45° \\ \text{or} \\ \text{changeover} \;\Rightarrow\; +30° \le \alpha \le +35° \;\Rightarrow\; -45° \le \beta \le 0° \end{cases} \tag{H2a}$$

**3/** Possibility for a WS head to be situated in the kites n° k or n° (k+1)

Second case of location in the adjacent kite illustrated in Fig H1b:

$$+25° \le \alpha \le +30° \quad \Rightarrow \begin{cases} -45° \le \beta \le 0° \\ \text{or} \\ \text{changeover} \;\Rightarrow\; -35° \le \alpha \le -30° \;\Rightarrow\; 0° \le \beta \le +45° \end{cases} \tag{H2b}$$

The 4 values ±5° and ±25° are chosen arbitrarily. Except for these 3 special cases, rules (1a) and (1b) of Paper 3 remain applicable. Conditions (6) and (7) are retained but condition (8) is modified by increasing the range of β, in accordance with the limits indicated in (16) in accompanying Paper 2, that is:

$$|\,\beta\,| \le 45° \tag{H3}$$

When the α angle verifies one of the conditions set out in (H1), (H2a) or (H2b), a computer routine tests the proposed alternatives while following the rules (6) and (7) of Paper 3 supplemented by (H3) and the case where a WS occurs is retained. With these optimized conditions, WS potentialities are counted and ordered according to the θ angle. As an example, the occurrences θi for Mrow n° 5 appear on the 6 graphs in Fig H2 which refer to the even steps. Compared to Fig 3 of Paper 3, there is an increase in the number of occurrences from 4 to 5 values. Among the 288 heads belonging to a Mfil, all the WS possibilities under optimized conditions are displayed in the 6 graphs in Fig H3. The discrete distributions of θi between $\theta_{down}$ and $\theta_{up}$ appear more homogeneous compared to those in Fig 4 of Paper 3.

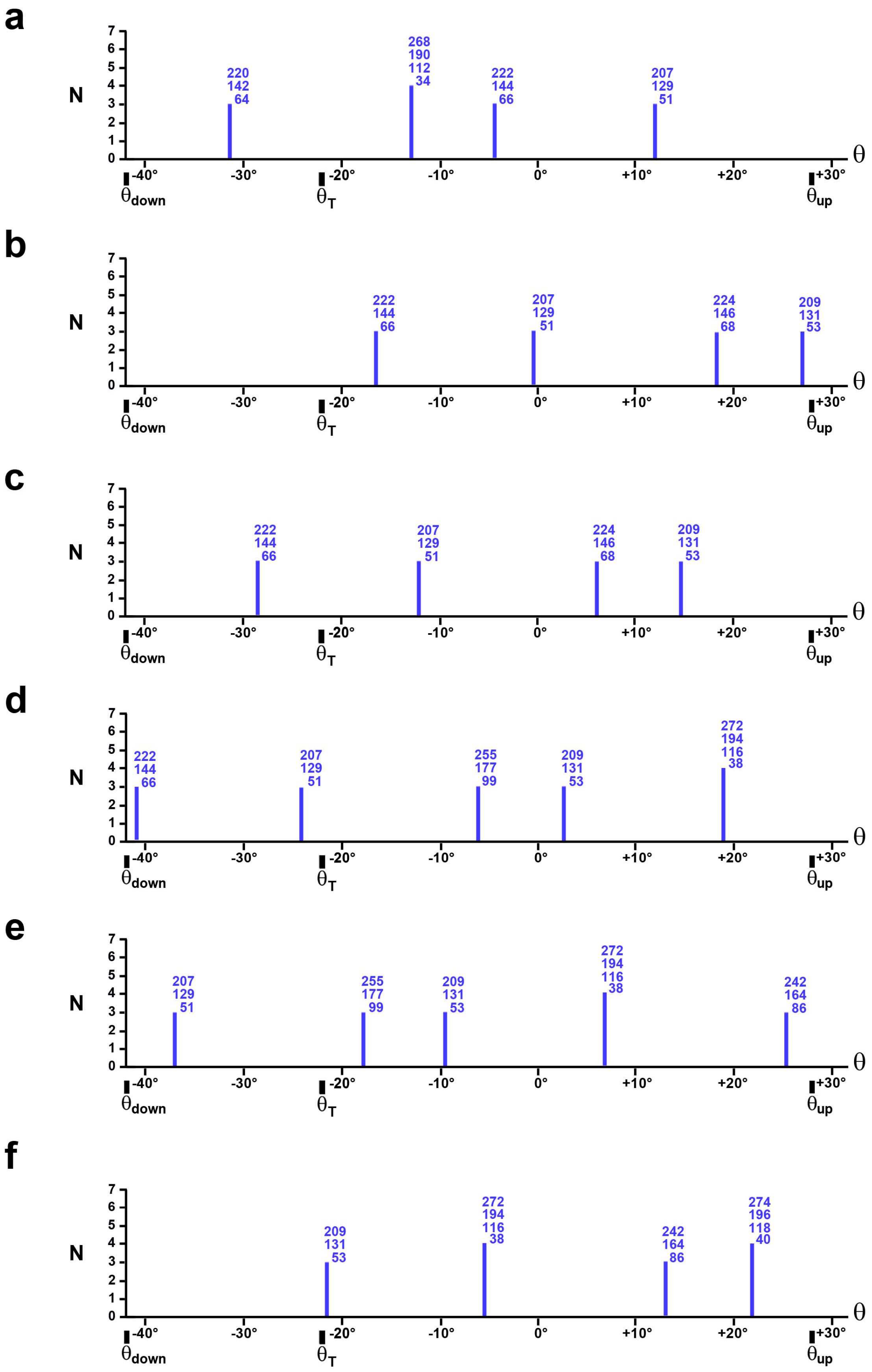


**Fig H2. Maximum number of WS possibilities for the 32 myosin heads of the Mrow n° 5 under optimized conditions with $\alpha_0 = -5°$, $\beta_0 = 0°$ and $|\beta| \leq 45°$.**

(a) Reference position of the hsR whose fixed length is between 1 and 1.1 µm. (b), (c), (d), (e) and (f) After hsR shortening equal to 2 nm, 4 nm, 6 nm, 8 nm and 10 nm, respectively.

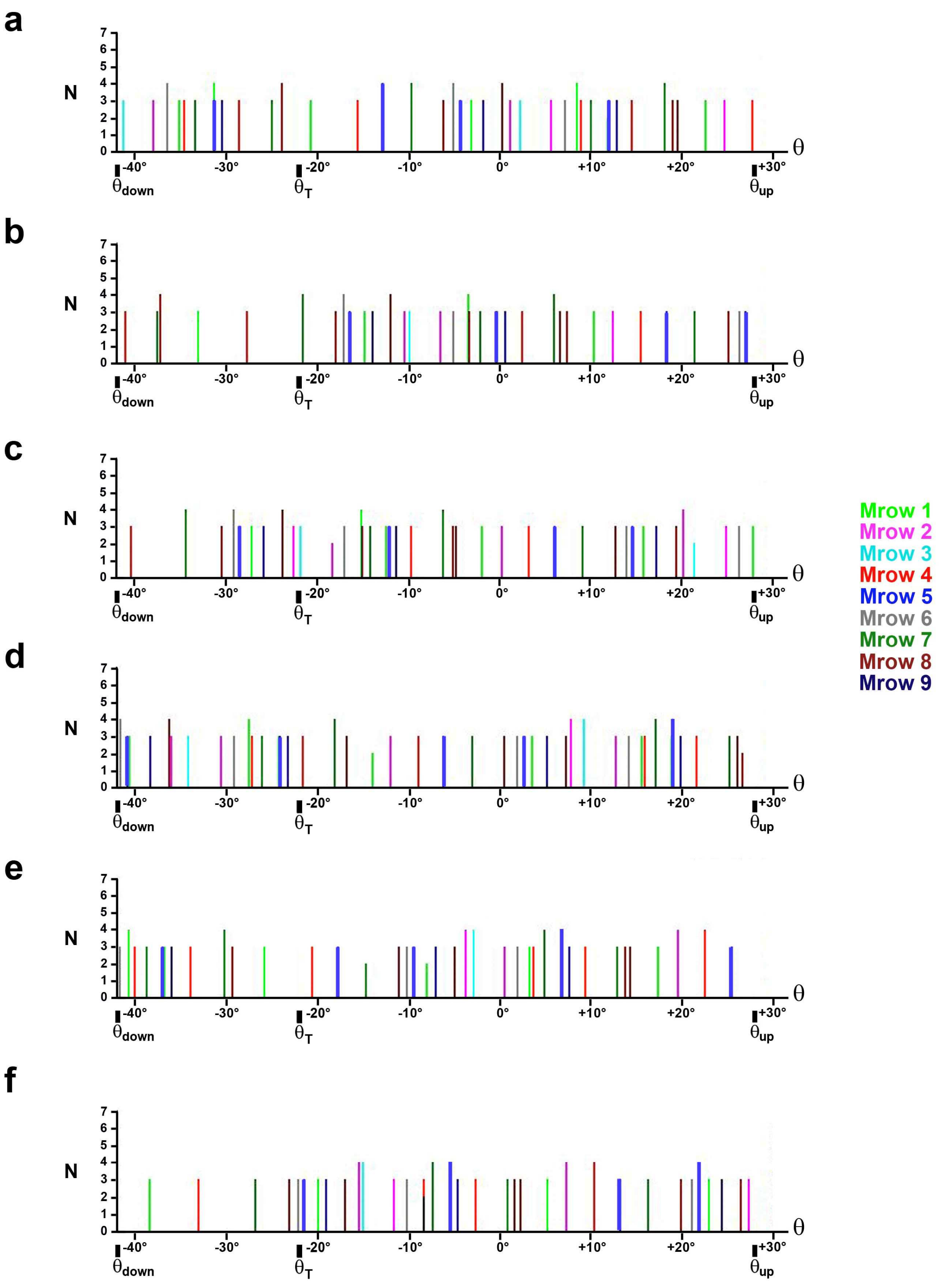


**Fig H3. Maximum number of WS possibilities for the 288 myosin heads of the 9 Mrow of a Mfil under optimized conditions with $\alpha_0 = -5°$, $\beta_0 = 0°$ and $|\beta| \leq 45°$.**

(a) Reference position of the hsR whose fixed length is between 1 and 1.1 µm. (b), (c), (d), (e) and (f) After hsR shortening equal to 2, 4, 6, 8 and 10 nm, respectively.

Each vertical line color refers to the color and therefore to the number of one of the 9 Mrow of the legend.

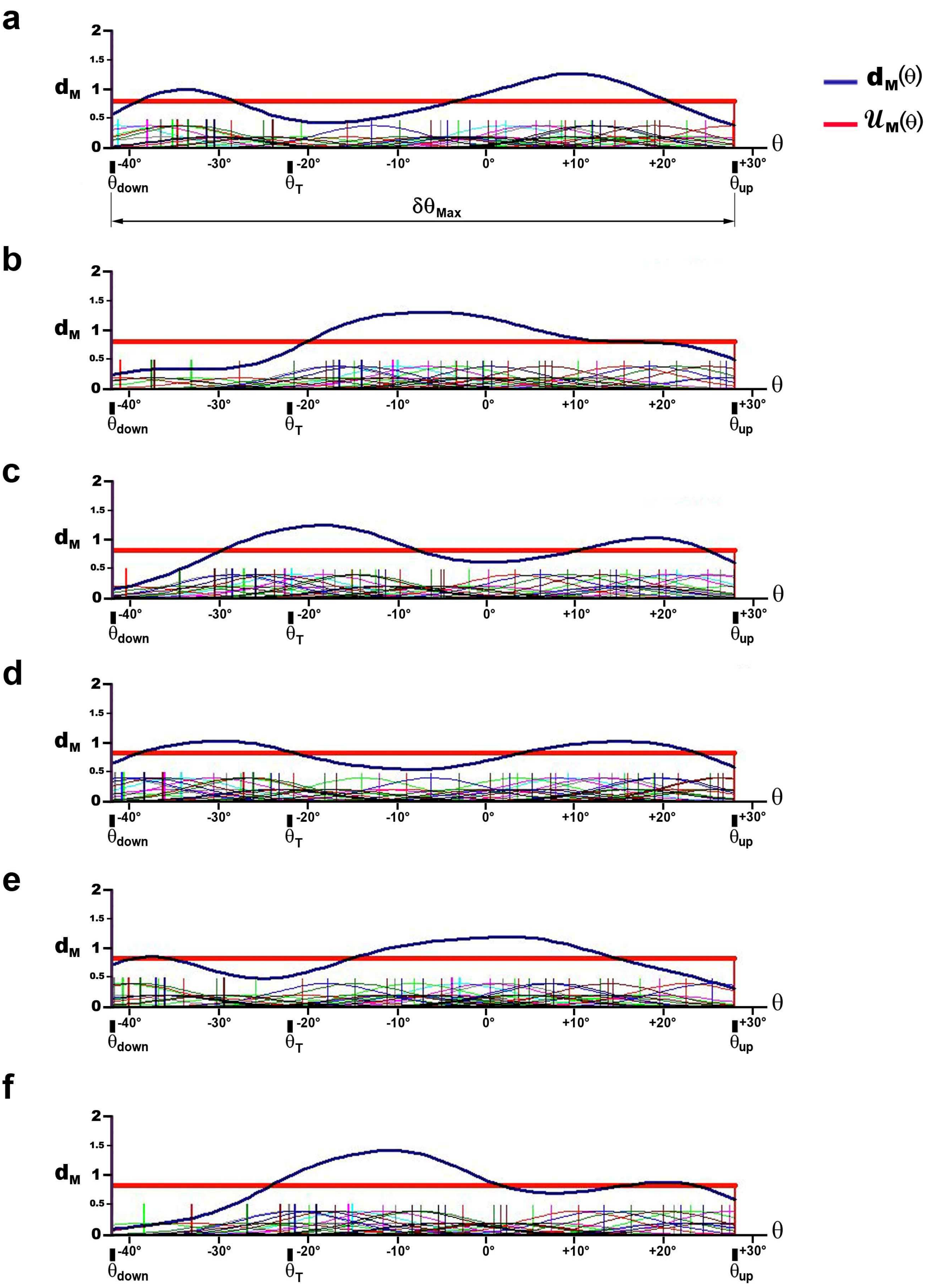


**Fig H4. Probability density $d_G$ calculated under optimized conditions with $\alpha_0$ = -5°, $\beta_0$ = 0° and |β| ≤ 45°.**

(a) The length of the hsR is fixed between 1 and 1.1 µm.

(b), (c), (d), (e) and (f) After hsR shortening equal to 2, 4, 6, 8 and 10 nm, respectively.

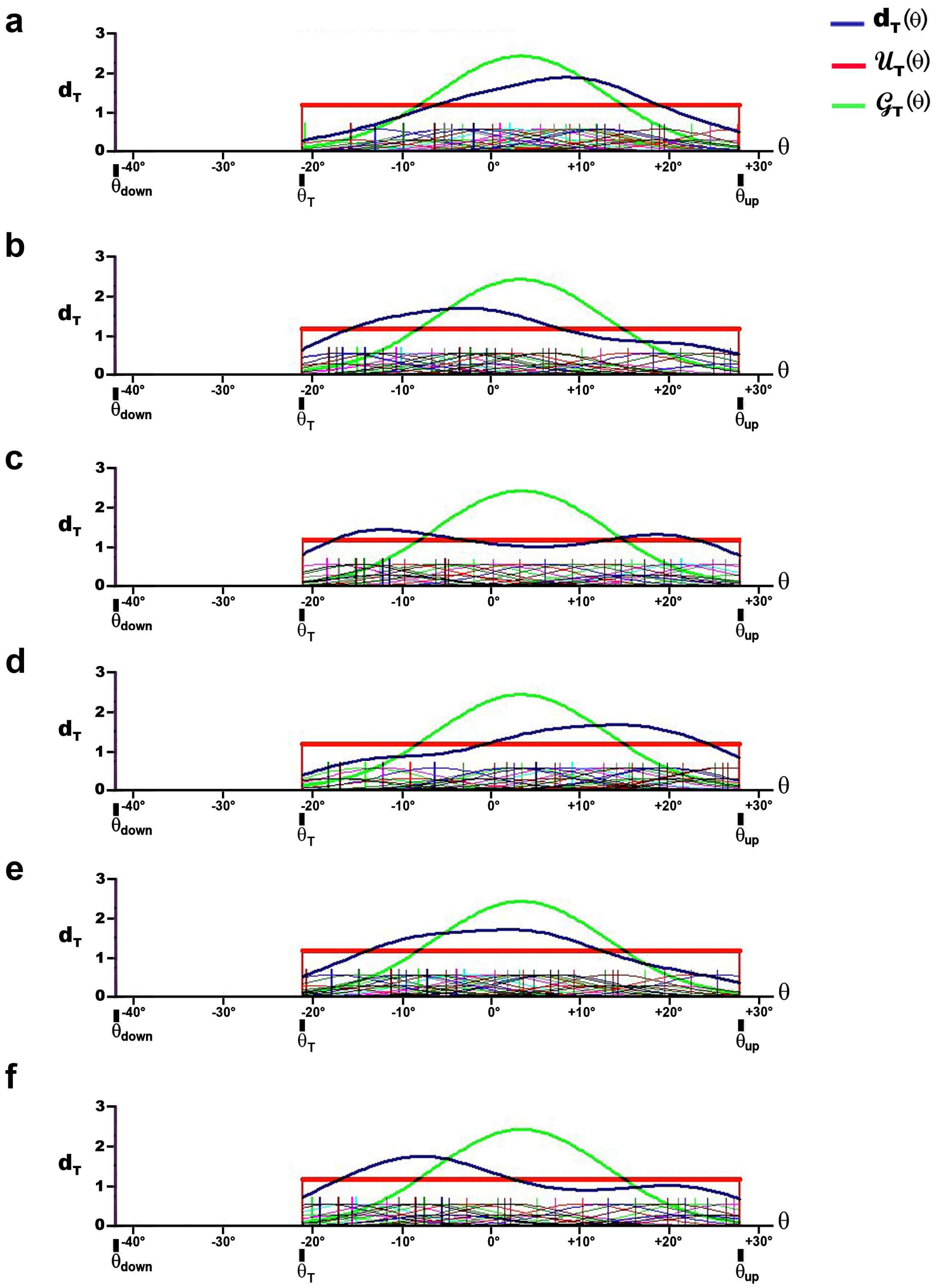


**Fig H5. Probability density $d_T$ calculated under optimized conditions with $\alpha_0$ = -5°, $\beta_0$ = 0° and $|\beta| \leq 45°$.**

(a) The length of the hsR is fixed between 1 and 1.1 µm.

(b), (c), (d), (e) and (f) After hsR shortening equal to 2, 4, 6, 8 and 10 nm, respectively.

Again, the variability of $\theta$ is explained by assigning to each $\theta i$ a normal probability density with a mean ($\theta i$) and a common standard deviation ($\sigma_\theta$) equal to 6.1°. For the reference position and for each of the 11 hs shortenings, a function ($f_M$) and the associated probability density ($d_M$) are determined according to equations (12) and (13), respectively. The density $d_M$ is represented by a solid blue line on the 6 graphs in Fig H4, corresponding to even shortenings. Among the classical statistical laws, the most convincing law for approaching the 12 densities $d_M$ is again the Uniform law expressed in (14) in Paper 3, a law characterized by a red horizontal line on the 6 graphs in Fig H4.

##### H.2 Probability densities of $\theta$ under optimized conditions over the interval $\delta\theta_T$

The angle $\theta_T$ is characterized as (16). The discrete values $\theta i$ between $\theta_T$ and $\theta_{up}$ are selected (Fig H3). For the reference position and for each of the 11 hs shortenings, a function ($f_T$) and the associated probability density ($d_T$) are determined according to equations (17) and (18), respectively. The density $d_T$ is plotted with a thick blue on the 6 graphs in Fig H5. Each probability density $d_T$ is compared with the uniform law ($\mathcal{U}_T$ ) and the Gaussian law ($\mathcal{G}_T$) defined on the interval $\delta\theta_T$ according to (19) and (20). The $\mathcal{U}_T$ law is represented by a red horizontal line and the $\mathcal{G}_T$ law by a green line (Fig H5). By algorithmic procedure, the deviations $|d_T(\theta) - \mathcal{U}_T(\theta)|$ and $|d_T(\theta) - \mathcal{G}_T(\theta)|$ are summed for 100 $\theta$ values between $\theta_T$ and $\theta_{up}$ and incremented with the same pitch. The Gaussian model gives a total of the deviations 2.2 times greater than that of the uniform model.

##### H.3 Calculation of the maximum theoretical percentage of myosin heads in WS

WS heads are counted taking into account the probabilistic weight ($p_i$) introduced in the expression (12), i.e. when several neighbouring sites are eligible for the WS of a head, only one option is selected. The resulting figure is divided by the total number of heads per hs, 288 according to Table 1, which provides the maximum percentage of WS heads per hs.

The percentage is calculated over the range $\delta\theta_{Max}$ ($pWS_M$).

On average over the 12 shortening steps (Table H1; columns 3 and 5):

$pWS_M$ is 18% under standard conditions as defined by (1a), (1b), (6), (7) and (8).

$pWS_M$ is 29% under optimized conditions with the additional cases specified in (H1), (H2a), (H2b) and (H3).

The percentage is calculated on the range $\delta\theta_T$ ($pWS_T$).

On average over the 12 shortening steps (Table H2; columns 3 and 5):

$pWS_T$ is 13% under normal conditions.

$pWS_T$ is equal to 25% under optimized conditions.

**Table H1**: **Percentage of WS on $\delta\theta_{Max}$ ($pWS_M$) for hsR shortenings from 0 to 11 nm according to 2 condition types**

| | | Normal conditions with $\|\beta\| \leq 30°$ | | Optimized conditions With $\|\beta\| \leq 45°$ |
|---|---|---|---|---|
| **ΔX** | **Fig**[(1)] | **$pWS_M$** | **Fig** [(2)] | **$pWS_M$** |
| 0 | 5a | 16% | H4a | 29% |
| -1 nm | | 17% | | 29% |
| -2 nm | 5b | 16% | H4b | 25% |
| -3 nm | | 18% | | 29% |
| -4 nm | 5c | 19% | H4c | 31% |
| -5 nm | | 21% | | 34% |
| -6 nm | 5d | 19% | H4d | 34% |
| -7 nm | | 17% | | 28% |
| - 8 nm | 5e | 17% | H5e | 29% |
| -9 nm | | 17% | | 26% |
| - 10 nm | 5f | 17% | H4e | 28% |
| -11 nm | | 19% | | 31% |
| **Average** | | 17.8% | | 29.4% |
| **Standard deviation** | | 1.5% | | 2.7% |

[(1)] The figures belong to Paper 3. [(2)] The figures belong to Supplement S3.H.

**Table H2**: **Percentage of WS on $\delta\theta_T$ ($pWS_T$) for hsR shortenings from 0 to 11 nm according to 2 condition types**

| | | Normal conditions with $\|\beta\| \leq 30°$ | | Optimized conditions with $\|\beta\| \leq 45°$ |
|---|---|---|---|---|
| **ΔX** | **Fig** | **$pWS_T$** | **Fig** | **$pWS_T$** |
| 0 | 6a | 12% | H5a | 24% |
| -1 nm | | 13% | | 26% |
| -2 nm | 6b | 13% | H5b | 25% |
| -3 nm | | 13% | | 27% |
| -4 nm | 6c | 12% | H5c | 25% |
| -5 nm | | 16% | | 26% |
| -6 nm | 6d | 13% | H5d | 24% |
| -7 nm | | 13% | | 22% |
| - 8 nm | 6e | 14% | H5e | 25% |
| -9 nm | | 14% | | 26% |
| - 10 nm | 6f | 13% | H5e | 26% |
| -11 nm | | 14% | | 27% |
| **Average** | | 13.3% | | 25.3% |
| **Standard deviation** | | 1.1% | | 1.4% |

The stroke size ($\delta X_{Max}$) is equal to 11.5 nm according to (5) with $\delta\theta_{Max}$=70°. We check (Table 1):

$$2 \cdot L_{Amol} = 11 \text{ nm} \qquad \text{(H4)}$$

Let be the distance separating 3 adjacent Asit on the same Ahel, length close to $\delta X_{Max}$. According to Laplace's definition, the maximum theoretical probability of achieving WS on $\delta\theta_{Max}$ ($pWS_{Max,th}$) is equal to the ratio of the number of favourable cases to the number of possible cases. Geometrically, $pWS_{M,theor}$ is calculated as the ratio of the distance between 3 Asit to the Amotif length, either from (H4) and the data in Table 1 :

$$pWS_{Max,theor} \approx \frac{2 \cdot L_{Amol}}{L_{Amotif}} \approx 30\% \qquad \text{(H5)}$$

This figure is close to 29%, the average of $pWS_M$ collected at the bottom of the "optimized conditions" column in Table H1.

The theoretical maximum probability of WS realization on $\delta\theta_T$ ($pWS_{T,theor}$) is calculated according to a rule of three:

$$pWS_{T,théor} = pWS_{M,theor} \cdot \frac{\delta\theta_T}{\delta\theta_{Max}} = 30\% \cdot \frac{49°}{70°} \approx 22\% \qquad \text{(H6)}$$

This figure is slightly less than 25%, the average of $pWS_T$ collected at the bottom of the "optimized conditions" column in Table H2.
