## Supplementary material for "Mechanical model of muscle contraction. 3. The orientation of the levers belonging to the myosin heads in working stroke follows the same uniform law in all half-sarcomeres of an isometrically stimulated fiber": Computer programs for calculating and plotting densities of Teta angle

### CP3 Computer programs for calculating and plotting densities of θ angle

Calculations and plots always start by running “Sub AAMM_BUFFON()” routine.

1/ The first step consists in creating Access tables with “Sub AAMM_BUFFON_0_CREA_Tables()” routine.

2/ Tables are filled with the necessary data for starting.

3/ Once the conditions are selected, clicking on OK button (Fig CP3.1) starts the “Sub_ AAMM_BUFFON_angle _Teta_stocke” : all calculations are performed as described in the paragraph “Determination of the number of myosin heads potentially in WS” of the Methods section

4/ The graphs are plotted with Sub_ AAMM_BUFFON_angle_Teta_trace_Angle (fig CP3.2 et CP3..3).

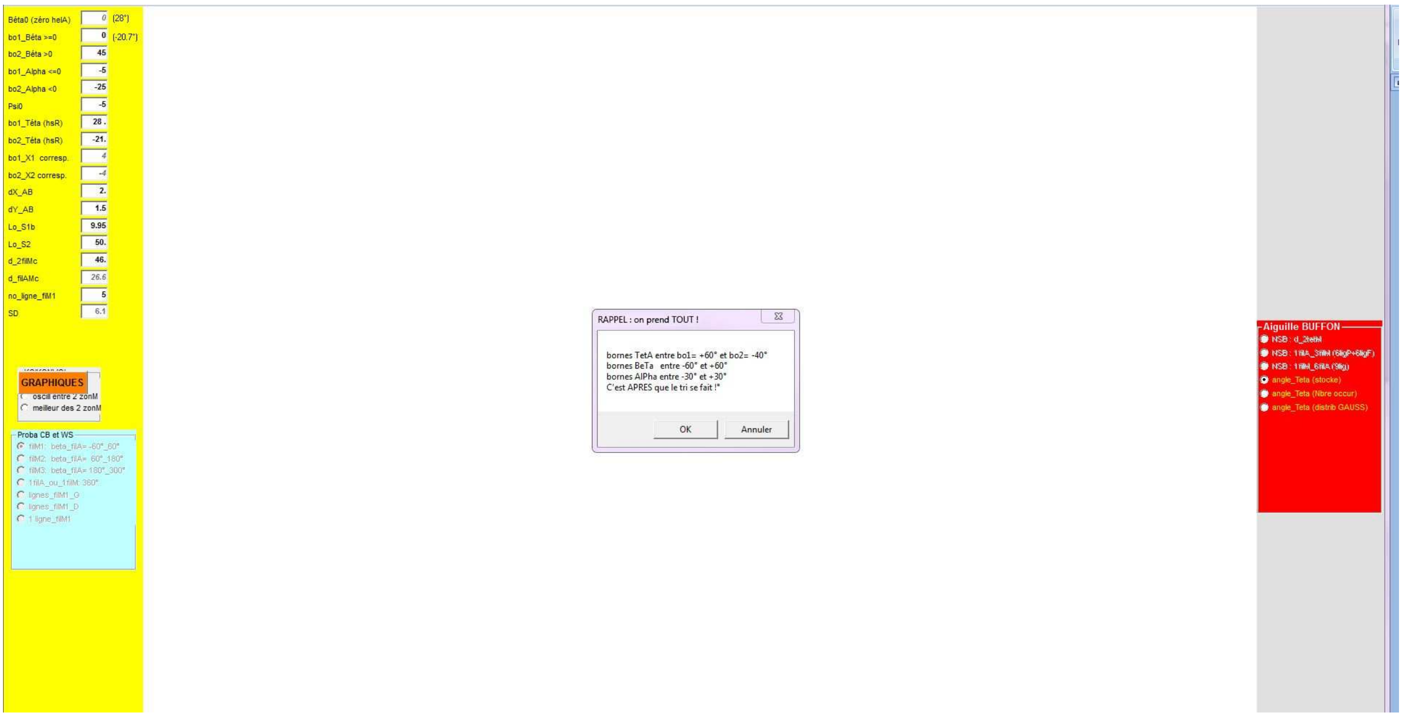


**Fig CP3.1. Screenshot after starting the routine "AAMM_BUFFON()" to start the calculations.**

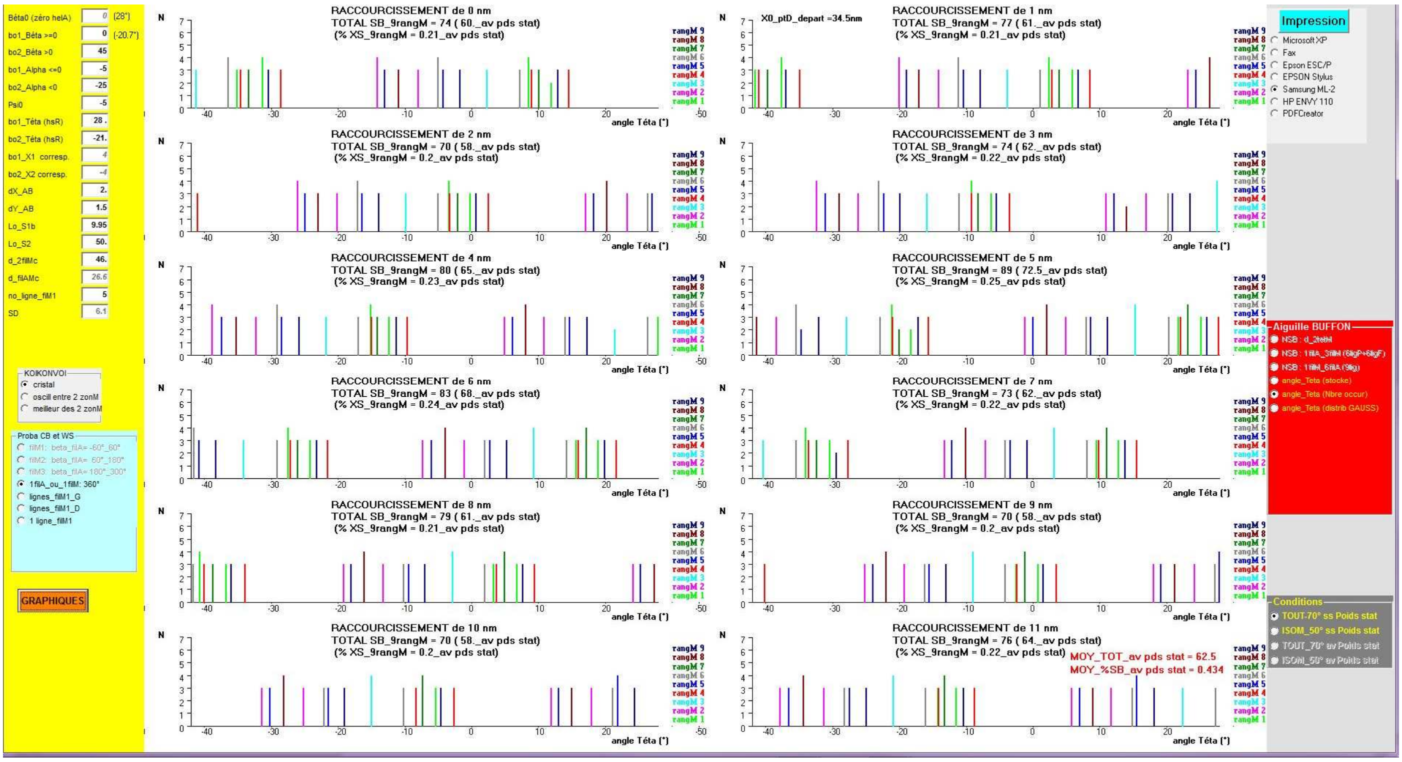


Fig CP3.2. Screenshot after starting the routine "AAMM_BUFFON()" to start the plot of the angular positions θi.

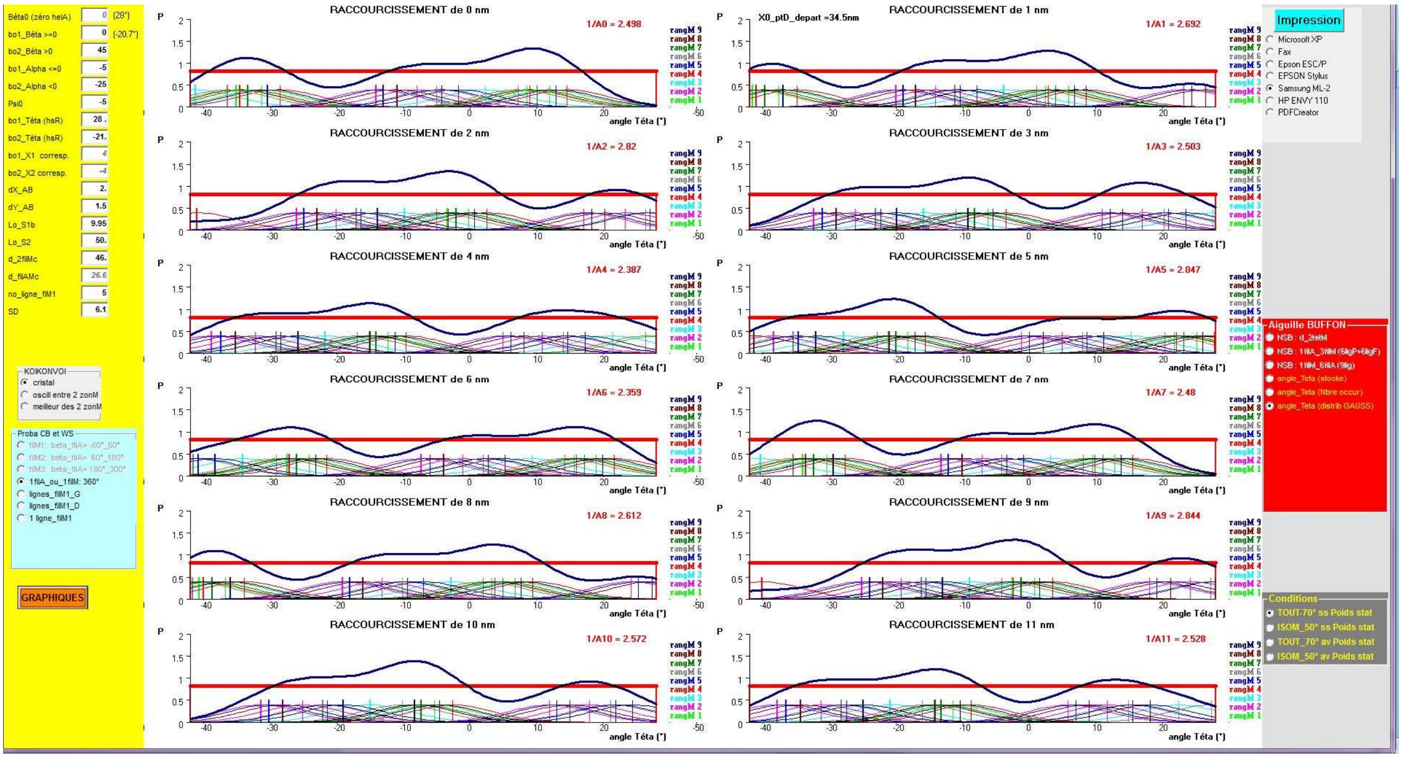


Fig CP3.3. Screenshot after starting the " AAMM_BUFFON()"to start the plot of the θi distributions.

#### Sub AAMM_BUFFON()

```
Dim tot(1 To 13) As Integer, BeTa_k As Single

'If mmm = 1 Then
O.Cls
X.Cls
' X0 = -5 '-10IIf(mmm = 1, -15, -25)  'decalge vers la droite pour misen page impression
' Y0 = -20
' larg = 110 '115IIf(mmm = 1, 140, 130) ' 140 'IIf(CYCIMP!Chi(3).Value = 0, 140, 170)
' haut = 140
' hentete = 7: hpage = 20
X0 = -10
larg = 120
Y0 = -30

'bon on y est
'***etape praparation table filCB_16tetM
tx1 = "BUFFON_X" 'X =tous les sitA sur 715 nm d'un filA (zonz correspondant au recouvrement complet filM/filA + raccourcissement de 15 nm
tx2 = "BUFFON_N" 'nbre de CB possibles
tx3 = "BUFFON_A" 'angulation teta raprtiotn gaussienne vers repartition uniforme
tx4 = "BUFFON_O" 'ordre des angles pour tracés teta raprtiotn gaussienne vers repartition uniforme
txV = "BUFFON_Verif" ' en fait en doublon avec loiBA mais laisser
zYb = "BUFFON_G"
'**ICI***
drap = 2 '0= cree table+remplit  1:remplit champs sitA   2=routine
'si besoin de tout recréer il faut enlever cette ligne
'If FrBUFF.Tag > 0 Then drap = 2 'on pass en routine

'---Pas ICI---c'est AU DESSUS
If drap = 0 Then AAMM_BUFFON_0_CREA_Tables 'on créée les 5 tables + copie à la main de BUFFON_Double

'-------ouverture des 4 tables---------
Set loiBX = dbM.OpenRecordset("BUFFON_X", dbOpenTable)
loiBX.Index = "X"
Set loiBN = dbM.OpenRecordset("BUFFON_N", dbOpenTable)
loiBN.Index = "PK"
Set loiBA = dbM.OpenRecordset("BUFFON_A", dbOpenTable)
loiBA.Index = "PK"
Set loiBO = dbM.OpenRecordset("BUFFON_O", dbOpenTable)
loiBO.Index = "PK"
Set loiBR = dbM.OpenRecordset("BUFFON_RWS", dbOpenTable)
loiBR.Index = "PK"
Set loiBV = dbM.OpenRecordset(txV, dbOpenTable)
loiBV.Index = "PK"
Set loiBG = dbM.OpenRecordset("BUFFON_G", dbOpenTable)
loiBG.Index = "PK"

'------------- mise à jour de la table R_WS
r_filA = 3.5 ''nm par défaut
r_filM = 7.5 'nm par défaut
'XS1a = Val(TAM(10).Text) '<0 versions 123_2016
XS1a = -Val(TAM(10).Text) '<0 versions >=4_2017
YS1a = Val(TAM(11).Text) '
'LS1b = 11 '9 'nm par défaut
BeTArd = 0 ' 10 * pi / 180 ' mes calculs ont ete fait fait pour 10°(compromis entre 0 et 20°)
ALPHA_S2rd = 0
Xd = 0 'on s'en fout cest =juste pour verifer la formeule

vd = "on passe"
'vd = "MAJ"
```

```
If vd = "MAJ" Then 'rappel les angles ont ete mesure ds hsL puis changé de signes mais AAMM_BUFFON_calcul_X_ptA_ds_hsR
  loiBR.MoveFirst
  Do Until loiBR.EOF()
    LS1b = loiBR("L_S1b")
    aX = loiBR("delta_start_X")
    LS2 = loiBR("L_S2")
    d_2filMc = loiBR("d_2filMc")
    d_filAMc = d_2filMc * Sqr(3) / 3
    TetaF = loiBR("tetaF_hsR") 'en deg
    Teta1 = loiBR("teta1_hsR") 'en deg
    Teta3 = loiBR("teta3_hsR") 'en deg
    R_WS = loiBR("R_WS")
    loiBR.Edit
      loiBR("d_filAMc") = d_filAMc

     'Teta et delta_Teta
      loiBR("delta_pre_teta") = TetaF - Teta1
      loiBR("delta_MAX_teta") = Teta1 - Teta3
      loiBR("delta_TOUT_teta") = TetaF - Teta3

     'on repasse en degré
      Teta1 = loiBR("teta1_hsR")
      Teta2 = Teta1 - aX * 180 / (R_WS * LS1b * pi)
      loiBR("teta2_hsR") = Int(Teta2 * 10 + 0.5) / 10
      Teta0 = (Teta1 + Teta2) / 2
      loiBR("teta0_ISO") = Int(Teta0 * 10 + 0.5) / 10
      Teta0 = (TetaF + Teta2) / 2
      loiBR("teta0_PRE_ISO") = Int(Teta0 * 10 + 0.5) / 10
      loiBR("delta_ISO_teta") = Int((TetaF - Teta2) * 10 + 0.5) / 10
      loiBR("delta_start_teta") = Int((Teta1 - Teta2) * 10 + 0.5) / 10

     'L_WS ds hsR (d'où l'explication du signe négatif)
      Teta1 = Teta1 * pi / 180: X1 = AAMM_BUFFON_calcul_X_ptA_ds_hsR(Teta1)
      Teta3 = Teta3 * pi / 180: X3 = AAMM_BUFFON_calcul_X_ptA_ds_hsR(Teta3)

      loiBR("delta_MAX_X") = Int((X1 - X3) * 10 + 0.5) / 10
      loiBR("delta_MAX_X_exact") = X1 - X3 '=10 nm


      TetaF = TetaF * pi / 180: X2 = AAMM_BUFFON_calcul_X_ptA_ds_hsR(TetaF)
      loiBR("delta_TOUT_X") = Int((X2 - X3) * 10 + 0.5) / 10
      loiBR("delta_TOUT_X_exact") = X2 - X3

      Teta2 = (TetaF - Teta1) / 2
       X1 = AAMM_BUFFON_calcul_X_ptA_ds_hsR(Teta2)
       X2 = AAMM_BUFFON_calcul_X_ptA_ds_hsR(-Teta2)
      loiBR("delta_pre_X_exact") = X1 - X2
      loiBR("delta_pre_X") = Int((X1 - X2) * 10 + 0.5) / 10 ' X1 - X2 '=4 à 5.5 nm

     '------------
    loiBR.Update
    loiBR.MoveNext
  Loop
End If
If vd = "MAJ" Then Stop 'on vérifie

'***etap 2: on remplit
'N_HelA = 2
'N_sitA_par_motif = 13
'Dim LS1a As Single, LS1b  As Single, LS2 As Single
```

```
'Dim BeTA As Single, d_filAMc As Single, r_filA As Single, r_filM As Single, dY_AM As Single, XS1a As Single, YS1a As Single, LS2_p As Single

'variables
bo1_dg = Val(TAM(6).Text) 'bo1=borne_TetaF_preWS_hsR
bo2_dg = Val(TAM(7).Text) 'bo2=borne_Teta3_finWS_hsR
LS1b = Val(TAM(12).Text)
LS2 = Val(TAM(13).Text)
d_2filMc = Val(TAM(14).Text): If d_2filMc < 40 Then Stop 'pas normal
d_filAMc = d_2filMc * Sqr(3) / 3
TAM(15).Text = Format(d_filAMc, "##.#")

loiBR.Seek "=", LS1b, LS2, d_2filMc:
If loiBR.NoMatch Then
   vd = MsgBox("données non incorporées dans la table Buffon_RWS", 0, "d_f2filM + Ls1b + LS2")
   Exit Sub
End If
'attention les mesures d'angles ont été faites en hsL et maintenet on passe en hsR
TetaF = loiBR("tetaF_hsR")
Teta1 = loiBR("teta1_hsR") '+ 30 '°
Teta2 = loiBR("teta2_hsR") '- 16.2° dteta_startWS=4 7.5°
Teta3 = loiBR("teta3_hsR") ' - 32.5 '° dteta_MAX=67 .5°
R_WS = loiBR("R_WS") '0.945
aX = loiBR("delta_start_X") ' 6.5 à 7.5 nm

' on le marque
X.CurrentX = 7.8: X.CurrentY = 0.8: X.Print "(" + Format(Teta1) + "°)"
X.CurrentX = 7.8: X.CurrentY = 3.2: X.Print "(" + Format(Teta2) + "°)"

'constantes

d_2tetM = 42.9 'nm = 3 * 14.3nm par défaut
p_HelA = 2.75 'nm par défaut
Ard_HelA = 2 * pi / 13 'Atn(1) * 8 / N_sitA_par_motif = 27.692°= 0.4833 rd

'calculs divers ds hsR
TetaF = TetaF * pi / 180 'rd
Teta1 = Teta1 * pi / 180 'rd
Teta2 = Teta2 * pi / 180 'rd
Teta3 = Teta3 * pi / 180 'rd
Teta0 = (Teta1 + Teta2) / 2 'rd

bo1_rd = bo1_dg * pi / 180 'rd   b=borne_Teta_preWS
If bo1_rd < 0 Then Stop 'on est ds hsR MERDE
bo2_rd = bo2_dg * pi / 180 'rd
If bo2_rd > 0 Then Stop 'on est ds hsR MERDE

bo1_nm = LS1b * R_WS * (bo1_rd - Teta0)
bo2_nm = LS1b * R_WS * (bo2_rd - Teta0)

bo1_nm = Int(bo1_nm * 10 + 0.5) / 10 'b=borne_X_preWS
bo2_nm = Int(bo2_nm * 10 + 0.5) / 10


k = 8: AAMM_label_ds_table_filD k
filD.Edit: filD("vv_d") = bo1_nm: filD.Update
TAM(8).Text = Format(bo1_nm) 'Format(bo1_nm, "#0.#")
k = 9: AAMM_label_ds_table_filD k
filD.Edit: filD("vv_d") = bo2_nm: filD.Update
TAM(9).Text = Format(bo2_nm) 'Format(bo2_nm, "#0.#")
'rappel calcul inverse : b_rd = Teta0 - b_nm /(LS1b * R_WS)
```

```
If drap < 2 Then
   If drap = 1 Then dbM.Execute "DELETE * FROM " + tx1

    ' Renvoie une valeur aléatoire comprise entre 0 et 60
    Beta0 = Val(Right(Format(Time), 2))
    If Beta0 = 60 Then Beta0 = 0

    Beta0 = 0

     'enreg de la valeur
    k = 0: AAMM_label_ds_table_filD k
    filD.Edit: filD("vv_d") = Beta0: filD.Update
    'affichage de la val beta0 calculée
    TAM(0).Text = Format(Beta0)
    'on passe en rd
    Beta0 = Beta0 * Atn(1) / 45

    For z = 1 To 360 ' 300*2.75nm= 825 nm pour couvrir L_filM=70nm + 125 nm de raccourcissement (zone 1.1µm à 1µm par hs)
       motif = Int(z / 13 - 0.01) + 1
       i = z Mod 13 ''N_sitA_par_motif = 13
       If i = 0 Then i = 13

       loiBX.AddNew
        loiBX("no") = z
        loiBX("mot13_2hel") = i
        loiBX("X") = p_HelA * z
        If z / 2 > Int(z / 2) Then 'helice 1 : s impair = 1,3,5,...
          loiBX("helA") = 1
          If motif / 2 > Int(motif / 2) Then 'N° impairs
             k = (i + 1) / 2
          Else 'no pairs
             k = 7 + i / 2
          End If
          loiBX("mot13_1hel") = k
          loiBX("sitA") = "M" + Format(motif) + "_h1_A" + Format(i)

        Else 'hélice 2 : s pair = 2,4,...
          loiBX("helA") = 2
          If motif / 2 > Int(motif / 2) Then 'N° impairs
             k = 7 + i / 2
          Else 'no pairs
             k = (i + 1) / 2
          End If
          loiBX("sitA") = "M" + Format(motif) + "_h2_A" + Format(i)
       End If

       'loiBX("A_X") = r_HelA * Cos(i * Ard_HelA)
       'loiBX("A_Y") = r_HelA * Sin(i * Ard_HelA)
       loiBX("mot13_1hel") = k
       BeTA = Beta0 + Ard_HelA * (k - 1)
       If BeTA > 2 * pi Then BeTA = BeTA - 2 * pi
       loiBX("Beta_rd") = BeTA
       loiBX("Beta_deg") = Int(BeTA * 1800 / pi + 0.5) / 10

       BeTA = BeTA * 180 / pi

      '***************
       For k = 1 To 6
          BeTa_k = BeTA - ((k + 2) Mod 6) * 60
          If BeTa_k >= -60 And BeTa_k <= 60 Then
             loiBX("Beta_" + Format(k)) = BeTa_k
             'If z = 2 Then Stop
```

```
            End If
            If z Mod 13 = 1 Then loiBX("Beta_3") = 60
            If z Mod 13 = 10 Then loiBX("Beta_4") = -2 * 360 / 13
            If z Mod 13 = 12 Then loiBX("Beta_4") = -360 / 13
        Next
        loiBX.Update
    Next
 If drap = 0 Then Stop 'vaut mieux tout relancer apres avoir mis : drap=2

Else 'drap=2
  For t = 1 To 20
    vx = "tetM_" + Format(t)
    dbM.Execute "UPDATE " + tx1 + " SET " + vx + " = Null"
  Next
End If

'***************
Screen.MousePointer = 11
If FrBUFF.Tag < 4 Then dbM.Execute "DELETE * FROM BUFFON_N"
If loiBX.RecordCount < 50 Then Stop 'table pas remplit
loiBX.MoveFirst
li = 0

sX = 1 ' nm par défaut (le filM avance sur le filA ce qui correspond bien à un racourcisst
       '= aussi pas/pitch de deplacment du filM/filA =ordre de grandeur de la variabilité

d_2tetM = 42.9 ' nm par defaut cste ds monde animal_chordés
N_tetM_Long = 16 ' Int(725 / d_2tetM) par defaut cste ds monde animal_chordés

bo1_Beta = Val(TAM(1).Text) 'bo1_Beta >=0
bo2_Beta = Val(TAM(2).Text) 'bo2_Beta <60°

bo1_Alpha = Val(TAM(3).Text) 'bo1_Alpha =0
bo2_Alpha = Val(TAM(4).Text) 'bo2_Alpha > - 30°
Psi0 = Val(TAM(5).Text) 'angle de départ de la 1ere ligne_16tetM du filM1

'FIXE
X0_ptD_depart = 34.5 '21.45 'nm 34.5 'dot etre inf à L_Amotif = 35.75 nm

'sX = 1 nm /pas(pitch) de déplacement du filM/filA
Npts = Int(p_HelA * 13 / sX - 0.1) ' = 36 =LmotifA = p_HelA/2 =71.5nm/2
                                   'avec p_HelA * 13 = 35.75 nm
'par précaution
For t = 1 To 9: no_Ligne(t) = 0: Next

Select Case FrBUFF.Tag
     Case 2: AAMM_BUFFON_6filA_autour_1filM
     Case 3: AAMM_BUFFON_angle_Teta_stocke
     Case 4, 5: AAMM_BUFFON_angle_Teta_trace_Angle
    'Case 5: AAMM_BUFFON_angle_Teta_trace_GAUSS
    Case Else: Stop 'a faire
    End Select

Screen.MousePointer = 0
End Sub
```

#### Sub AAMM_BUFFON_0_CREA_Tables()

```
 'Dim ntb As TableDef, ind As Index, fd As Field

'1 loi_BUFFON_X (X pour abscisse sur axe OX)
  For t = 0 To dbM.TableDefs.Count - 1
       If dbM.TableDefs(t).Name = tx1 Then dbM.TableDefs.Delete tx1: Exit For
   Next t
    Set ntb = New TableDef
    With ntb
     .Name = tx1
     .Fields.Append .CreateField("no", dbInteger) '0/
     .Fields.Append .CreateField("helA", dbInteger) '2/
     .Fields.Append .CreateField("mot13_2hel", dbInteger) '3/
     .Fields.Append .CreateField("mot13_1hel", dbInteger) '3/
     .Fields.Append .CreateField("sitA", dbText, 20) '1/
     .Fields.Append .CreateField("X", dbSingle) '4/
     .Fields.Append .CreateField("Beta_rd", dbSingle) '5
     .Fields.Append .CreateField("Beta_deg", dbSingle) '5
      For i = 1 To 6 '16*43nm=690nm (+10nm=700nm)
         '.Fields.Append .CreateField("Kyte_" + Format(i), dbInteger)
         .Fields.Append .CreateField("BeTa_" + Format(i), dbSingle)
         .Fields.Append .CreateField("BeTa_RA_" + Format(i), dbSingle) 'RA= rotation aléatoire
     Next 'i

     For i = 1 To 20 '16*43nm=690nm (+10nm=700nm)
         .Fields.Append .CreateField("tetM_" + Format(i), dbSingle)
     Next 'i

     Set ind = New Index
     ind.Name = "PK": ind.Fields = "+no": ind.Primary = True: ind.Unique = True
     .Indexes.Append ind
     Set ind = New Index
     ind.Name = "X": ind.Fields = "+X": ind.Primary = False: ind.Unique = True
     .Indexes.Append ind
    End With
    dbM.TableDefs.Append ntb

'2 loi_BUFFON_N (N pour Nombre de SB)
    For t = 0 To dbM.TableDefs.Count - 1
       If dbM.TableDefs(t).Name = tx2 Then dbM.TableDefs.Delete tx2: Exit For
   Next t
    Set ntb = New TableDef
    With ntb
     .Name = tx2
     .Fields.Append .CreateField("comm", dbText, 20) '1/
     .Fields.Append .CreateField("d_2tetM", dbSingle) '0/
     .Fields.Append .CreateField("no_courbe", dbInteger) '0/
     .Fields.Append .CreateField("rangM", dbInteger) '0/
     .Fields.Append .CreateField("pasX", dbSingle) '2/
     .Fields.Append .CreateField("N_SB", dbInteger)
     .Fields.Append .CreateField("p_SB", dbSingle)

     Set ind = New Index
     ind.Name = "PK": ind.Fields = "+no_courbe;+pasX": ind.Primary = True: ind.Unique = True
     .Indexes.Append ind

    End With
    dbM.TableDefs.Append ntb

'3 loi_Buffon_A (A pour Angle)
  For t = 0 To dbM.TableDefs.Count - 1
       If dbM.TableDefs(t).Name = tx3 Then dbM.TableDefs.Delete tx3: Exit For
```

```
Next t

Set ntb = New TableDef
 With ntb
  .Name = tx3
  .Fields.Append .CreateField("no_courbe", dbInteger) '0/
  .Fields.Append .CreateField("pasX", dbSingle) '2/
  .Fields.Append .CreateField("rangM", dbInteger) '0/
  .Fields.Append .CreateField("zonM60", dbInteger) '0/
  .Fields.Append .CreateField("entre_2_zonM", dbInteger) '0/
  .Fields.Append .CreateField("Psi", dbSingle) '0/
  .Fields.Append .CreateField("Alpha_S2", dbSingle) '0/
  .Fields.Append .CreateField("no_A", dbInteger)
  .Fields.Append .CreateField("tetM16", dbInteger)   'no de la tetM sur les 16 qui accroche le sitA
  .Fields.Append .CreateField("tetM16_no_pos", dbInteger) 'no de la positon 1=meilleur 2 3 ?
  .Fields.Append .CreateField("poids_stat", dbSingle) '0/
  .Fields.Append .CreateField("Alin", dbSingle)
  .Fields.Append .CreateField("A", dbSingle)
  .Fields.Append .CreateField("A_N_interp", dbInteger)
  .Fields.Append .CreateField("Ac_en_deg", dbSingle)
  .Fields.Append .CreateField("Ac_en_deg_N_interp", dbInteger)
  .Fields.Append .CreateField("Ac_en_deg_mult_6point3", dbSingle)
  .Fields.Append .CreateField("sitA", dbInteger)
  .Fields.Append .CreateField("Beta_360_sitA", dbSingle)
  .Fields.Append .CreateField("Beta_PouM_60_sitA", dbSingle)  'ds zone 120°=+ ou - 60 (PlusouMoins)
  .Fields.Append .CreateField("X_sitA", dbSingle)
  .Fields.Append .CreateField("X_ptA", dbSingle)
  .Fields.Append .CreateField("X_ptAlin", dbSingle)
  .Fields.Append .CreateField("LS2_p", dbSingle)
  .Fields.Append .CreateField("angle_FI0", dbSingle)

  Set ind = New Index
  ind.Name = "PK": ind.Fields = "+no_courbe;+rangM;+no_A": ind.Primary = True: ind.Unique = True
  .Indexes.Append ind
   Set ind = New Index
  ind.Name = "Teta": ind.Fields = "+no_courbe;+rangM;+A": ind.Primary = False: ind.Unique = False: ntb.Indexes.Append ind

 End With
 dbM.TableDefs.Append ntb

'4 loi_Buffon_O (O pour Ordre/ordonnancement)
      For t = 0 To dbM.TableDefs.Count - 1
          If dbM.TableDefs(t).Name = tx4 Then dbM.TableDefs.Delete tx4: Exit For
      Next t
       Set ntb = New TableDef
       With ntb
        .Name = tx4
        '.Fields.Append .CreateField("o", dbInteger) 'o=0 tout 1=cas 1 2=cas 3 etc aléatroire Tote
        .Fields.Append .CreateField("cond", dbInteger)
        .Fields.Append .CreateField("pasX", dbSingle) '2/
        .Fields.Append .CreateField("rangM", dbInteger) '0/
        .Fields.Append .CreateField("zonM60", dbInteger) '0/
        .Fields.Append .CreateField("BeTA", dbSingle) '0/
        .Fields.Append .CreateField("Alpha_S2", dbSingle) '0/
        .Fields.Append .CreateField("A_val", dbSingle) '0/
        .Fields.Append .CreateField("Alin_val", dbSingle) '0/
        .Fields.Append .CreateField("N_sitA", dbInteger)
        .Fields.Append .CreateField("poids_stat", dbSingle) '0/

        'en verite je vous le dis 4 doit suffire
        For i = 1 To 8 '16*43nm=690nm (+10nm=700nm)
            vx = "sitA" + Format(i)
```

```
            .Fields.Append .CreateField(vx, dbInteger)

        Next 'i

        Set ind = New Index
        ind.Name = "PK": ind.Fields = "+cond;+pasX;+rangM;+A_val": ind.Primary = True: ind.Unique = True
        .Indexes.Append ind
    End With
      dbM.TableDefs.Append ntb


'5 loi_Buffon_Verif (V pour verif en fait doublon de Buffon_Angle
  For t = 0 To dbM.TableDefs.Count - 1
     If dbM.TableDefs(t).Name = txV Then dbM.TableDefs.Delete txV: Exit For
  Next t

  Set ntb = New TableDef
   With ntb
    .Name = txV
   .Fields.Append .CreateField("no_courbe", dbInteger) '0/
    .Fields.Append .CreateField("pasX", dbSingle) '2/
    .Fields.Append .CreateField("rangM", dbInteger) '0/
    .Fields.Append .CreateField("zonM60", dbInteger) '0/
    .Fields.Append .CreateField("entre_2_zonM", dbInteger) '0/
    .Fields.Append .CreateField("Psi", dbSingle) '0/
    .Fields.Append .CreateField("Alpha_S2", dbSingle) '0/
    .Fields.Append .CreateField("no_A", dbInteger)
    .Fields.Append .CreateField("i_yt", dbInteger)
    .Fields.Append .CreateField("tetM16", dbInteger)   'no de la tetM sur les 16 qui accroche le sitA
    .Fields.Append .CreateField("poids_stat", dbSingle) '0/
    .Fields.Append .CreateField("Alin", dbSingle)
    .Fields.Append .CreateField("A", dbSingle)
    .Fields.Append .CreateField("A_N_interp", dbInteger)
    .Fields.Append .CreateField("Ac_en_deg", dbSingle)
    .Fields.Append .CreateField("Ac_en_deg_N_interp", dbInteger)
    .Fields.Append .CreateField("Ac_en_deg_mult_6point3", dbSingle)
    .Fields.Append .CreateField("sitA", dbInteger)
    .Fields.Append .CreateField("Beta_360_sitA", dbSingle)
    .Fields.Append .CreateField("Beta_PouM_60_sitA", dbSingle)  'ds zone 120°=+ ou - 60 (PlusouMoins)
    .Fields.Append .CreateField("X_sitA", dbSingle)
    .Fields.Append .CreateField("X_ptA", dbSingle)
    .Fields.Append .CreateField("X_ptAlin", dbSingle)
    .Fields.Append .CreateField("X_ptAc_en_deg", dbSingle)
    .Fields.Append .CreateField("diffX_ptA", dbSingle)
    .Fields.Append .CreateField("diffX_ptAc_en_deg", dbSingle)
    .Fields.Append .CreateField("LS2_p", dbSingle)
    .Fields.Append .CreateField("angle_FI0", dbSingle)

    '------------

    Set ind = New Index
    ind.Name = "PK": ind.Fields = "+no_courbe;+rangM;+no_A;+i_yt": ind.Primary = True: ind.Unique = True
    .Indexes.Append ind

   End With
   dbM.TableDefs.Append ntb


'6 loi_BUFFON_G (G=distribution GAUSS)
   For t = 0 To dbM.TableDefs.Count - 1
     If dbM.TableDefs(t).Name = zYb Then dbM.TableDefs.Delete zYb: Exit For
  Next t
```

```
    Set ntb = New TableDef
    With ntb
     .Name = zYb
     .Fields.Append .CreateField("pt", dbInteger) '1/
     .Fields.Append .CreateField("Aire", dbSingle) '0/
     .Fields.Append .CreateField("1/Aire", dbSingle) 'N=nomalisé/
     .Fields.Append .CreateField("Moy_Aire", dbSingle) '0/
     .Fields.Append .CreateField("Moy_Aire_Norm", dbSingle) 'N=nomalisé/
     .Fields.Append .CreateField("Moy_arith", dbSingle) '0/
     .Fields.Append .CreateField("Moy_arith_Norm", dbSingle) 'N=nomalisé/
     For i = 0 To 11 '
         vx = "sG_" + Format(i)
         .Fields.Append .CreateField(vx, dbSingle)
     Next 'i
     .Fields.Append .CreateField("Moy_Ect_LU", dbSingle) 'Moyenne des ecarts/LU
     .Fields.Append .CreateField("Moy_Ect_LG", dbSingle) 'Moyenne des ecarts/LG
     .Fields.Append .CreateField("LG", dbSingle) 'LG= loilaplace gauss pour comparer loi unifomre et gaussN=nomalisé/
     .Fields.Append .CreateField("Ecart_LU", dbSingle) 'ecart entre valeur calculée et val de la loi uniforme
     .Fields.Append .CreateField("Ecart_LG", dbSingle) 'ecart entre valeur calculée et par rapport à la loi uniforme


     Set ind = New Index
     ind.Name = "PK": ind.Fields = "+pt": ind.Primary = True: ind.Unique = True
     .Indexes.Append ind

    End With
    dbM.TableDefs.Append ntb

End Sub
```

#### Sub AAMM_BUFFON_angle_Teta_stocke()

```
Dim NsitA_succ As Integer, sitA_succ(1 To 4) As Integer, A_succ(1 To 4) As Single

'RAPPELS
'd_2tetM = 42.9 'nm
'N_tetM_Long = Int(700 / d_2tetM) = 16 S par rangM
If Fresc.Tag = 0 Then bo1_Alpha = 0 ': bo2_Alpha = -30'on tient pas compte des

vd = MsgBox("bornes TetA entre bo1= +60° et bo2= -40° " + vbCr + "bornes BeTa   entre -60° et +60°" + vbCr  + "bornes AlPha entre -30° et +30°" + vbCr + "C'est APRES que le tri se  fait !°", 1, "RAPPEL : on prend TOUT !")
If vd = 2 Then Exit Sub

dbM.Execute "DELETE * FROM " + tx3 'loiBA

'AU HASARD
  'Randomize 50
  'X0_ptD_depart = Int(Rnd * 50) / 10 'nm
'FIXE
 ' X0_ptD_depart = 34.5 '21.45 'nm 34.5

  'on prend des bornes excessives +1° pour variabil ité intrinsèque
  bo1_dg = 28 ' 30 '29 '30 '°
  bo2_dg = -42 ' -40 '-43 '-45 '°

  bo1_rd = bo1_dg * pi / 180 'rd   b=borne_Teta_preWS
  bo2_rd = bo2_dg * pi / 180 'rd

  'si on passe par sub
  'LS2_p = Sqr(LS2 ^ 2 - ((r_filA + YS1a + LS1b * Cos(angle_Teta_rd)) * Sin(BeTArd) + r_filM * Sin(ALPHA_S2rd)) ^ 2)
  'yx = d_filAMc - (r_filA + YS1a + LS1b * Cos(angle_Teta_rd)) * Cos(BeTArd) - r_filM * Cos(ALPHA_S2rd) '>0
  'I1 = ASINr(yx / LS2_p) 'en rd


   Xd = 0
   BeTArd = 10 * pi / 180 '10°
   ALPHA_S2rd = 0
   xmi = AAMM_BUFFON_calcul_X_ptA_ds_hsR(Teta0) 'teata en rd
   xx = AAMM_BUFFON_calcul_X_ptA_ds_hsR(bo1_rd)
   xn = AAMM_BUFFON_calcul_X_ptA_ds_hsR(bo2_rd)
   bo1_nm = xx - xmi
   bo2_nm = xn - xmi

  bo1_nm = Int(bo1_nm * 10 + 0.5) / 10 'b=borne_X_preWS
  bo2_nm = Int(bo2_nm * 10 + 0.5) / 10

  S1 = 0: S2 = 0

' i obligatoire car incremment du déplacment de pas sX = 1 nm
For i = 0 To 11 ' on a que 12 courbes disponibles pour 12 déplacement et de 0 à 11 nm et donc 12 repartionns gaussienes qui doivent (hyp) conduire à la m^me distribution
     'Psi0 = 20
    For rangM = 1 To 9  ' 1 à 9 rang M réprties entre filM1_G  et filM1_D

       'etape 1 calcul de l'angle alpha propre à chaque rang>M
         xmi = Psi0 + 40 * (rangM - 1) 'xmi = angle Psi = alpha_360(r) eq(1.8) varie de -20°=-20°+40°x(1 -1)=0°-20° à 340°=+20+40°x(9-1)=320°+20°
         If xmi >= 330 Then xmi = xmi - 360 'on repase dans kite_1

        'l'indice 1 car il faudra 1 moment choisir entre 2 zones_cerf-volant_kite variant de 1 à 6
         zonM60(1) = Int((xmi + 30) / 60) 'de 0 à 5  = z_60  eq(1.9)
```

```
        ALPHA_S2 = xmi - 60 * zonM60(1) ' = alpha(r) eq(1.10)


        '   Fresc.Tag = 0 >> cristal pur pas de chgt zone possibles
        '   Fresc.Tag = 1 >> chgt zone possibles/ a priroi j'ai laissé tomber ce truc
        '   fresc.tag=  2 >> choix augmenté

        If Fresc.Tag = 2 And Abs(ALPHA_S2) >= Abs(bo2_Alpha) Then 'bo2_alpha entre -30° et -20°
          entre_2_zonM = IIf(ALPHA_S2 < 0, -30, 30)  '= sur_le_fil entre 2 zonM60 -1 fuadra prendre la zonz d'avant +1 la zone d'apres
        Else
          entre_2_zonM = 0
        End If

        zonM60(1) = zonM60(1) + 1 ' zonM60= kite de 1 à 6
        If zonM60(1) = 7 Then Stop

        X0_ptD_alea = X0_ptD_depart + 14.3 * ((rangM - 1) Mod 3) '27
```

**AAMM_BUFFON_calculs_NSB_1lig_16tetM_autour_1filA**

```
    Next 'rangM

     '2 on fait un décompte ds loiBN totes rangM confondu
      sYY = 0
     loiBA.Seek ">=", i + 1, 1, 1: If loiBA.NoMatch Then Stop 'cela veut dire qu'aucune tetM de cette rangM n'a pu s'accrocher ? à verifier
     Do While loiBA("no_courbe") = i + 1
         sYY = sYY + 1
         loiBA.MoveNext
         If loiBA.EOF Then Exit Do
     Loop

     loiBN.AddNew
           loiBN("d_2tetM") = 1
           loiBN("no_courbe") = i + 1
           loiBN("rangM") = 0 'on s'en fout on fait la somme
           loiBN("pasX") = i * sX
           loiBN("N_SB") = sYY
     loiBN.Update
Next 'i=courbes obliger de fermer ici

'Stop 'on vérifie

'et maintenant on ordonne tout ça dans table tx4=loiBO
'ce qui suppose que ligne et coube sont associe àVERFIER
dbM.Execute "DELETE * FROM " + tx4
'je mets de l'ordre dans la angles avec une nouvelle base
loiBA.Index = "teta"

'bo1_Beta = Val(TAM(1).Text) 'bo1_Beta >=0
        'bo2_Beta = Val(TAM(2).Text) 'bo2_Beta <60°

        'bo1_Alpha = Val(TAM(3).Text) 'bo1_Alpha =0
        'bo2_Alpha = Val(TAM(4).Text) 'bo2_Alpha > - 30°
 bo1_dg = Val(TAM(6).Text) 'bo1=borne_Teta1_preWS_hsR
 bo2_dg = Val(TAM(7).Text) 'bo2=borne_Teta2_preWS_hsR
        'BeTA = Abs(loiBA("BetA_PouM_60_sitA"))
       'ALPHA_S2 = Abs(loiBA("ALPHA_S2"))

For k = 0 To 1 'k=cond 0=tout 1 =cond
For i = 0 To 11 'les 12 deplacemnt
For rangM = 1 To 9  ' 1 à 9 rang M réprties entre filM1_G  et filM1_D
```

```
   loiBA.Seek ">=", i + 1, rangM, -90: If loiBA.NoMatch Then Stop
   'on prend tout
   'en fait ici mettre condition spllment 'selon les 3 angles Teta/BeTa/Alpha
   'on crre la 1ere valeur

   Do While loiBA("no_courbe") = i + 1 And loiBA("rangM") = rangM

     'Conditions : 1 je laisse tomber alpha
     '            2 Je laisse tomber BeTA >= bo1_Beta And
     '                                                    30°              -17°
     'If k = 0 Or (k = 1 And Abs(loiBA("BetA_PouM_60_sitA")) <= bo2_Beta And loiBA("A") <= bo1_dg And loiBA("A") >= bo2_dg)
Then
     If Abs(loiBA("BetA_PouM_60_sitA")) <= bo2_Beta And (k = 0 Or (k = 1 And loiBA("A") <= bo1_dg And loiBA("A") >= bo2_dg))
Then
         loiBO.Seek "=", k, i, rangM, loiBA("A")
         If loiBO.NoMatch Then 'pas trouvé donc on crrée
           loiBO.AddNew
             loiBO("cond") = k ' 0 = TOUT 1=avec conditions
             loiBO("pasX") = i
             loiBO("rangM") = loiBA("rangM")
             loiBO("zonM60") = loiBA("zonM60")
             loiBO("BeTA") = loiBA("BetA_PouM_60_sitA")
             loiBO("Alpha_S2") = loiBA("Alpha_S2")
             loiBO("poids_stat") = loiBA("poids_stat")
             loiBO("A_val") = loiBA("A")
             loiBO("Alin_val") = loiBA("Alin")
             loiBO("N_sitA") = 1
             loiBO("sitA1") = loiBA("sitA")
           loiBO.Update

         Else 'existe deja donc je mets à jour
           'petit verif
           'If loiBO("rangM") <> loiBA("rangM") Then Stop
           If loiBO("zonM60") <> loiBA("zonM60") Then Stop
           If Fresc.Tag = 0 And loiBO("BeTA") <> loiBA("BetA_PouM_60_sitA") Then Stop
           'If loiBO("Alpha_S2") <> loiBA("Alpha_S2") Then Stop
           'If loiBO("poids_stat") <> loiBA("poids_stat") Then Stop

           loiBO.Edit
             sN = loiBO("N_sitA") + 1
             loiBO("N_sitA") = sN
             loiBO("sitA" + Format(sN)) = loiBA("sitA")
           loiBO.Update

         End If
     End If

     loiBA.MoveNext
     If loiBA.EOF Then Exit Do
   Loop
Next 'rangm
Next 'i =deplacments
Next 'k = conditions

'--------------
'on recalcul le vrai poids car je me suis rendu compte que selon les conditions toutes les vlaeur de teta n'étaient pas forcement
retenues
'donc obliger de le refaire et de le stocker dans sitA8
For k = 0 To 1 'k=cond 0=tout 1 =cond
For i = 0 To 11 'les 12 deplacemnt
For rangM = 1 To 9  ' 1 à 9 rang M réprties entre filM1_G  et filM1_D
```

```
    loiBO.Seek ">=", k, i, rangM, -90: If loiBO.NoMatch Then Stop
    Do While loiBO("cond") = k And loiBO("pasX") = i And loiBO("rangM") = rangM
      If IsNull(loiBO("sitA8")) Then
          NsitA_succ = 1
          sitA_succ(NsitA_succ) = loiBO("sitA1")
          A_succ(NsitA_succ) = loiBO("A_val")
           'Debug.Print "****** REPER"
          For z = 1 To 12
             loiBO.MoveNext
             If loiBO.EOF Then Exit For
             If loiBO("rangM") <> rangM Then Exit For
             If loiBO("sitA1") - sitA_succ(NsitA_succ) = 2 Then
               NsitA_succ = NsitA_succ + 1
               sitA_succ(NsitA_succ) = loiBO("sitA1")
               A_succ(NsitA_succ) = loiBO("A_val")
             End If
          Next 'z
           For z = 1 To NsitA_succ
             loiBO.Seek "=", k, i, rangM, A_succ(z): If loiBO.NoMatch Then Stop
             loiBO.Edit: loiBO("sitA8") = NsitA_succ: loiBO.Update

          Next 'z

         loiBO.Seek ">=", k, i, rangM, A_succ(1)

      End If
      loiBO.MoveNext
      If loiBO.EOF Then Exit Do
    Loop
Next 'rangm
Next 'i =deplacments
Next 'k = conditions


'*******************************

vd = MsgBox("tu peux tracer", 0, "OK")


End Sub
```

```
Sub AAMM_BUFFON_6filA_autour_1filM()
'     *  *
'   *  O  *
'     *  *
'd_2tetM = 42.9 ' nm ici on ne bouge plus
'N_tetM_Long = 16 ' Int(725 / d_2tetM)
sXY = 0 'total des CB pour les lignes_16tetM appartenat aux 3 filM dont les tetM peuvent etre en liaison avec le fiLA

For sN = 1 To 9 '9 lignes  car 9 x 40°= 360°
    'on simplifie
    xmi = Psi0 + 40 * (sN - 1) + 30
    k = Int(xmi / 60)
    If k < 0 Then Stop 'k=0
    xmi = Psi0 + 40 * (sN - 1) - k * 60

     vd = "lignePAStrouvée" 'on retrouve nos petits

                '          0°                  -2 0°
                If xmi <= bo1_Alpha And xmi >= bo2_Alpha Then ' alhpa_S2 negatif donc beta postif
                   'beta positf
                    Beta1 = bo1_Beta 'borne min
                    Beta2 = bo2_Beta 'borne maxi
                     vd = "lignetrouvée" 'c'est ok

                '              20°                    0°
                ElseIf xmi <= -bo2_Alpha And xmi >= -bo1_Alpha Then ' alhpa_S2 positif donc Beta négatif
                  'beta negatif
                   Beta1 = -bo2_Beta 'borne min
                   Beta2 = -bo1_Beta 'borne maxi
                    vd = "lignetrouvée"   'c'est ok
                End If

     If vd = "lignetrouvée" Then
        ALPHA_S2 = xmi
        no_filA = k + 1
       ' X0_ptD_alea
       q = 10
        If Fresc.Tag = 0 Then 'cristal
            k = sN Mod 3: If k = 0 Then k = 3
            X0_ptD_alea = q + 14.3 * k
        Else
           Randomize ' 15
            If Fresc.Tag = 1 Then ' aleatoire +/- 10°: on juste sur 1 nm
               X0_ptD_alea = q + Int(30 * Rnd + 0.5) / 10
            Else 'totalement aléatoire
               X0_ptD_alea = q + Int(360 * Rnd + 0.5) / 10
            End If
       End If
     End If

If vd = "lignetrouvée" Then
     sYY = 0 'somme des N_SB
     For i = 0 To Npts '=36 on se deplace sur l'axe OX logitudinal du pas sX=1nm   sur 35 nm =longueur du motif_13/6_molA
          AAMM_BUFFON_calculs_NSB_1lig_16tetM_autour_1filA 'à X fixé
     Next 'i
     sXY = sXY + sYY
    If sYY = 0 Then Stop

    'calcul p_SB/WS
    For i = 0 To Npts
        loiBN.Seek "=", sN, i * sX: If loiBN.NoMatch Then Stop
        loiBN.Edit
```

```
        If sYY > 0 Then
           loiBN("p_SB") = loiBN("N_SB") / sYY
        Else
         loiBN("p_SB") = 0
        End If
        If i = 0 Then loiBN("comm") = "N_SB = " + Format(sYY)
      loiBN.Update
   Next
 End If

 'et maintenant on trace avec sN = no de la courbe de 1 à 6

   vx = "deltaX (nm)"
   xn = 0
   xx = (Npts + 1) * sX ' p_HelA
   xt = sX 'p_HelA '1

   vy = "N": yn = 0: yx = 10: yt = 1
   'vy = "p": yn = 0: yx = 1: yt = 0.05

   Select Case sN
     '1ere col
     Case 1: impscale X0, Y0 + haut + hentete, X0 + 2 * larg, Y0 - 5 * haut - hpage
     Case 2: impscale X0, Y0 + 2 * haut + hentete, X0 + 2 * larg, Y0 - 4 * haut - hpage
     Case 3: impscale X0, Y0 + 3 * haut + hentete, X0 + 2 * larg, Y0 - 3 * haut - hpage
     Case 4: impscale X0, Y0 + 4 * haut + hentete, X0 + 2 * larg, Y0 - 2 * haut - hpage
     Case 5: impscale X0, Y0 + 5 * haut + hentete, X0 + 2 * larg, Y0 - haut - hpage
     Case 6: impscale X0, Y0 + 6 * haut + hentete, X0 + 2 * larg, Y0 - hpage
     Case 7: impscale X0 - larg, Y0 + haut + hentete, X0 + larg, Y0 - 5 * haut - hpage
     Case 8: impscale X0 - larg, Y0 + 2 * haut + hentete, X0 + larg, Y0 - 4 * haut - hpage
     Case 9: impscale X0 - larg, Y0 + 3 * haut + hentete, X0 + larg, Y0 - 3 * haut - hpage
   End Select

   'titre
   O.FontSize = 11: O.FontBold = -1
   impenvi

   loiBN.Seek "=", sN, 0
   If loiBN.NoMatch = 0 Then
     O.FontBold = -1
      O.CurrentX = 5: O.CurrentY = 110
      O.CurrentX = 5: O.CurrentY = 98:  O.Print "X_départ " + IIf(Fresc.Tag = 0, "fixé", "aléatoire") + " = " + Format(X0_ptD_alea) + " nm"
      O.CurrentX = 5: O.CurrentY = 85:  O.Print "Alpha_S2 = " + Format(ALPHA_S2) + "°"

      O.CurrentX = 5: O.CurrentY = 110
      O.Print "ligne n° " + Format(sN) + " de filM1  _" + Format(N_tetM_Long) + " tetM / " + loiBN("comm") + "      avec filA" + Format(no_filA)
      O.CurrentX = 40: O.CurrentY = 98
      z = (no_filA - 1) * 60
      O.Print "Zone_filM1 =[" + Format(z - 30) + "°;" + Format(z + 30) + "°] centrée sur " + Format(z)  + "°"

    O.FontBold = 0

    'barres en histogramme
      xav = 0
      For t = 0 To Npts
         loiBN.Seek "=", sN, t * sX ': If loiBN.NoMatch Then Stop
         If t > 0 Then O.Line (xap, 0)-(xap, yap)
         xap = ((t + 1) * sX - xn) * 100 / (xx - xn)
         yav = loiBN(vy + "_SB")
         yav = (yav - yn) * 100 / (yx - yn)
```

```
            O.Line (xav, yav)-(xap, yav)
            O.Line (xav, 0)-(xav, yav)
            yap = yav
            xav = xap ' xav = (t - xn) * 100 / (xx - xn)
        Next
        'on termine
        O.Line (xav, 0)-(xav, yap)
   Else
      O.CurrentX = 50: O.CurrentY = 102
      O.Print "Pas de ligne dans cette zone"
   End If

 Next 'sN pour calcul d_2tetM=35.75+2.75*sN

 impscale X0, Y0 + 6 * haut + hentete, X0 + 2 * larg, Y0 - hpage
  'rque en bas à droite
        'total CB des lignes
   'Case 1: O.Print "ordre " + Oesc(Fresc.Tag).Caption
   'Case 2: O.Print "d_filAM = " + TAM(15).Text + " nm"
   'Case 3: O.Print "ds hsR: bo1_Teta = " + TAM(6).Text + "° (Teta1 = " + Format(Int(Teta1 * 1800 / pi  + 0.5) / 10) + "°)"
   'Case 4: O.Print "ds hsR: bo2_Teta = " + TAM(7).Text + "° (Teta2 = " + Format(Int(Teta2 * 1800 / pi  + 0.5) / 10) + "°)"
   'Case 5: O.Print "d_bo = " + Format(Val(TAM(6).Text) - Val(TAM(7).Text)) + "° (dTeta1  " + Format(Int ((Teta1 - Teta2) * 1800 / pi
+ 0.5) / 10) + "°)"

     With O
        .FontSize = 14: .FontBold = -1
        'ordre
         O.CurrentX = 120: O.CurrentY = 140: O.Print "ordre " + Oesc(Fresc.Tag).Caption
        'd_filAM
         .CurrentX = 120: .CurrentY = 120
         O.Print "d_filAM = " + TAM(15).Text + " nm"
        'bornes teta
         .CurrentX = 120: .CurrentY = 100
         O.Print "ds hsR: bo1_Teta = " + TAM(6).Text + "° (Teta1 = " + Format(Int(Teta1 * 1800 / pi +  0.5) / 10) + "°)"
         .CurrentX = 120: .CurrentY = 80
         O.Print "ds hsR: bo2_Teta = " + TAM(7).Text + "° (Teta2 = " + Format(Int(Teta2 * 1800 / pi +  0.5) / 10) + "°)"

         .CurrentX = 120: .CurrentY = 60
         O.Print "d_bo = " + Format(Val(TAM(6).Text) - Val(TAM(7).Text)) + "°  (dTeta =  " + Format(Int(( Teta1 - Teta2) * 1800 / pi +
0.5) / 10) + "°)"

        'total CB
         .CurrentX = 120: .CurrentY = 35
         O.Print "TOTAL CB pour les 6 lignes 16_tetM = " + Format(sXY)
        .FontBold = 0
     End With

End Sub
```

#### Sub AAMM_BUFFON_calculs_NSB_1lig_16tetM_autour_1filA() 'à X fixé

```
'table filA remplie dans sub_AAMM_dist_calculs_sitA

Dim B1 As Integer, B2 As Integer, A_compteur As Integer, A_best As Single, okZ As Integer, Btxt As String
'PRNCIPE :
'on fait defiler les 16_tetM (ou + pour d_2tetM_varie) d'une mê rangée pour savoir combien sont susceptibles d'accrocher un sitA
'pour cela on doit au préalble savoir quel estle filA est entouré de 3 filM soit 120°(=360°/3) pour ch acun
'ou le film est entouré de 6 filA

'on doit donc savoir sur quelle zobe de filA (3 zones de 120°) et quelle rangee en face entre filM1 fil M2 et filM3 selon ordre ou aléatoire
'cette zonz de 120° est divisée en 2 pour repartiti ton des sitA (angle beta selon OX)

'**** SURTOUT pas de i , ni de sN
 sY = 0 'compteur de GP

 Xa = X0_ptD_alea + i * sX ' abscisse départ point D de la 1ere tetM de la rangée_16tetM sur axe OX
                          ' avec X0_ptD_alea zero X alétoire comris entre 0 et 35 nm calculé ds chaque subroutine appelante
                          ' et sX = pas de dépalcement
                          ' For i = 0 To Npts '0 à 35 nm pour histogramess variation d_2tetM
                          ' For i = 0 To 11 'nm  pour calculs_angle_Teta

    'on prépare table 'tx1 = "loi_BUFFON_N" en vidant les champs tetM
    '16*9 = 144 = 150 molM = 150tetM (1 ou G)
    '16*43nm=690nm (+10nm=700nm)
    If FrBUFF.Tag < 3 Then
       For t = 1 To 20 'laisser 20 faut tout effacer
          vx = "tetM_" + Format(t)
          dbM.Execute "UPDATE " + tx1 + " SET " + vx + " = Null"
       Next 't
    End If

     ALPHA_S2rd = ALPHA_S2 * pi / 180

    'on teste 1 à 1 les 16 S1 d'1 rangM (celles qui sont du bon cote doné par alpha qui done beta)
     t_test = 25 '25

     A_compteur = 0 'car plusiers possibilites de position angulaires teta par tetM

     For t = 1 To N_tetM_Long ' = L_film_hs/d_2tetM= 700/42.9 = 16,32 = 16 en general

        'principe on strocke totu ce qui concerne la tetM ds loiBV puis a la fin on recopie dans loiBA ce qui est le bon choix en fonction de frESC.tag
         dbM.Execute "DELETE * FROM " + txV 'loiBV erif

         Xd = Xa + (t - 1) * d_2tetM 'abscisse point D pour chacune des 16 tetM alignées sle long d'1 même rangée sur axe OX tous les d_2tetM

        'ATTENTION  raccourcissement ds hsR : dX_T=[Xmini ; X_maxi] avec X_mini correspond à bo2_Teta_rd(finWS)  et X_maxi à bo1_teta(debWS)

         xn = Xd + XS1a + LS2 - LS1b  'nm -10 à -15 corr à bo2_nm
         xx = Xd + XS1a + LS2 + LS1b  'nm  corr à bo1_nm

         If xx <= xn Then Stop 'verif OCAZOU

        'donc on parcourt de xx à xn (on fait comme pour 1 raccsst) pour trouver un sitA susceptible de coller
        '*** ATTENTION  on teste toutes les possibilités donc possibilite de 1 à 3 sitA possibles par tetM
        'on va en arriere on chercher la possibilité la plus longue dans le cadre d'1 raccsst
         F = 0 'compteur de possibilites de GP_CB dans loiBV
```

```
loiBX.Seek ">=", xx:  If loiBX.NoMatch Then Stop
Do While loiBX("X") >= xn 'calculs avec conditions Beta

  ''BeTA = loiBX("beta_deg") 'du filA ' = beta_360 (col 4 table 3)
  ''yav = BeTA 'pour le memoriser

  If Abs(entre_2_zonM) = 30 Then   'on tire au sort entre zonM60 et zonM60-1 ou zonM60+1
    'Fresc.Tag = 0 >> cristal pur pas de chgt zone possibles
    'Fresc.Tag = 1 >> chgt zone possibles
    If Fresc.Tag <> 2 Then Stop

       'on fait les 2(zonM60 et okZ pour trouver le meilleur
         'Stop 'a priro il faut recalcul S2_alpha dans le nouveau cerf_violant (il devient superie à -30 ou +30
       If entre_2_zonM = -30 Then 'zonM precedente
         ALPHA_S2 = 60 - Abs(ALPHA_S2)
          zonM60(2) = zonM60(1) - 1
          If zonM60(2) = 0 Then zonM60(2) = 6 'xmi=330°entre zonM60=6 et zonM60=1
      Else 'zonM suivante
          ALPHA_S2 = -60 + Abs(ALPHA_S2)
          zonM60(2) = zonM60(1) + 1
          If zonM60(2) = 7 Then zonM60(2) = 1
      End If
      'on teste les 2 options
     B1 = 1: B2 = 2

  Else 'normal 180 300 60
     B1 = 1: B2 = 1
  End If

  For B = B1 To B2

      'BeTA = yav - Choose(zonM60(B), 180, 240, 300, 0, 60, 120) '
      Btxt = "beta_" + Format(zonM60(B))
      If IsNull(loiBX(Btxt)) Then
        BeTA = 500 '*
      Else
        BeTA = loiBX("beta_" + Format(zonM60(B)))
      End If

      'cgt de zonzq
      If Fresc.Tag = 2 And Abs(ALPHA_S2) <= Abs(bo1_Alpha) Then ' à nouveau osicllation au hasard (mvt brownien)

          'on commence par celui_ci
            Beta1 = -60 '-bo2_Beta 'borne min
            Beta2 = 0 '-bo1_Beta 'borne maxi
            entre_2_zonM = -5

      'RAPPEL si cristal fresc.tag=0 ->> bo1_Alpha=0
         ' ne faudrait-il pas prendre 35°+30°+5° NON! car deja fait avant idiot
       '              -30°                -5° à 0]
      ElseIf ALPHA_S2 >= -40 And ALPHA_S2 <= bo1_Alpha Then  ' alhpa_S2 negatif donc beta postif
          'beta positf
            Beta1 = 0 'bo1_Beta 'borne min
            Beta2 = 60 'bo2_Beta 'borne maxi
               '         ]0 à +5°                +30°
      ElseIf ALPHA_S2 > -bo1_Alpha And ALPHA_S2 < 40 Then  ' alhpa_S2 positif donc Beta négatif
          'beta negatif
            Beta1 = -60 '-bo2_Beta 'borne min
            Beta2 = 0 '-bo1_Beta 'borne maxi
      Else
         Stop 'pas normal
      End If
```

```
              If i = 20 And rangM = 1 And t = t_test Then ' pas raccste variant de 0 à 11nm
                Debug.Print "rangM = " + Format(rangM), "tetM = " + Format(t), "entre_2_zonM " + Format(entre_2_zonM)
                 Debug.Print "no_sitA = " + Format(loiBX("no")), "X_sitA = " + Format(loiBX("X")) + " nm", "Beta_360° = " +
Format(loiBX("Beta_deg")) + "°"
                 Debug.Print "beta= " + Format(BeTA), "beta1= " + Format(Beta1), "beta2= " + Format(Beta2)
               End If

              '   0<beta1<beta<beta2        ou       beta1<beta<beta2<0
               If BeTA >= Beta1 And BeTA <= Beta2 Then
                    AAMM_BUFFON_calculs_NSB_1lig_16tetM_autour_1filA_OK

               If Fresc.Tag = 2 And entre_2_zonM = -5 Then 'on teste l'autre posiotn
                  Beta1 = 0 'bo1_Beta 'borne min
                  Beta2 = 60 ''bo2_Beta 'borne ma xi
                  entre_2_zonM = 5
                  If BeTA >= Beta1 And BeTA <= Beta2 Then
                     AAMM_BUFFON_calculs_NSB_1lig_16tetM_autour_1filA_OK
               End If

            Next ' B

            loiBX.MovePrevious: If loiBX.BOF Then Exit Do
         Loop
         'If i = 6 And rangM = 7 And t = t_test Then Stop
         'If t = 4 Then Stop

         'ici finalment pas de chox le choix se fera avec no_tetm et no_pos il faut effectuer le choix si il y en a 1 et le recopier dans
loiBA
         If loiBV.RecordCount > 0 Then 'ok on recopie
           If Fresc.Tag = 2 And (Abs(entre_2_zonM) = 30 Or Abs(entre_2_zonM) = 5) Then 'fresc.tag=2 le meilleur
              loiBV.MoveFirst
              A_best = -99
              Do Until loiBV.EOF
                If loiBV("A") > A_best Then A_best = loiBV("A"): okZ = loiBV("zonM60")
               loiBV.MoveNext
              Loop
              'Stop 'on verifie d'abord
              dbM.Execute "delete * from BUFFON_Verif WHERE [zonM60] <> " + Format(okZ)
           End If

           'pois_stat
            loiBV.MoveLast: Pdp = loiBV("no_A")

           loiBV.MoveFirst
           Do Until loiBV.EOF
              loiBA.AddNew
               loiBA("no_courbe") = i + 1
               loiBA("pasX") = i * sX
               loiBA("rangM") = rangM
               loiBA("zonM60") = loiBV("zonM60")
               loiBA("entre_2_zonM") = loiBV("entre_2_zonM")
               loiBA("Psi") = loiBV("Psi")
               loiBA("ALPHA_S2") = loiBV("ALPHA_S2")

               loiBA("tetM16") = t
               loiBA("tetM16_no_pos") = loiBV("no_A")
               loiBA("poids_stat") = 1 / Pdp

               loiBA("no_A") = A_compteur + loiBV("no_A")

               loiBA("Alin") = loiBV("Alin")
```

```
            loiBA("A") = loiBV("A")
            loiBA("A_N_interp") = loiBV("A_N_interp")
            loiBA("Ac_en_deg") = loiBV("Ac_en_deg")
            loiBA("Ac_en_deg_N_interp") = loiBV("Ac_en_deg_N_interp")
            loiBA("Ac_en_deg_mult_6point3") = loiBV("Ac_en_deg_mult_6point3")

            loiBA("sitA") = loiBV("sitA")

            loiBA("Beta_360_sitA") = loiBV("Beta_360_sitA")
            loiBA("Beta_PouM_60_sitA") = loiBV("Beta_PouM_60_sitA")
            loiBA("X_sitA") = loiBV("X_sitA")
            loiBA("X_ptA") = loiBV("X_ptA")
            loiBA("X_ptAlin") = loiBV("X_ptAlin")
           loiBA.Update

           loiBV.MoveNext
         Loop
         'If F <> loiBV("i_yt") Then Stop
         A_compteur = A_compteur + F
       Else
         If F > 0 Then Stop

       End If
    Next 'tetM


   If FrBUFF.Tag < 3 Then 'on analyse le resultat et on enregistre ds table_loiBN

    loiBN.AddNew
        loiBN("d_2tetM") = d_2tetM
        loiBN("no_courbe") = sN
        loiBN("rangM") = IIf(FrBUFF.Tag = 2, sN, no_Ligne(sN))
        loiBN("pasX") = i * sX

        loiBX.MoveFirst
        Do Until loiBX.EOF
          For t = 1 To N_tetM_Long
            vx = "tetM_" + Format(t)
            If IsNull(loiBX(vx)) = 0 Then
              sY = sY + 1
            End If
          Next
          loiBN("N_SB") = sY
          loiBX.MoveNext
        Loop

        sYY = sYY + sY 'SYY nbre total CB
    loiBN.Update

  End If


End Sub
```

#### Sub AAMM_BUFFON_calculs_NSB_1lig_16tetM_autour_1filA_OK()

```
Dim signX As Integer, pas_A As Single, i_yt As Integer, Rd_ou_Deg As Integer, teta_interpol As Single, A_N_interp As Integer

'RAPPEL: les angles sont calculées ds hsL mais ici les distances sont calculées ds hsR (donc inversion des angles et gaffe aux signes
 BeTArd = BeTA * Atn(1) / 45 'on passe en rd


  vd = "OK5" '"OK" si on veut editer ds Debug.print
  If i = 0 And rangM = 1 And t = t_test Then vd = "OK1"

 'X1 (bo1_teta) = position du startGP avt raccsst : = +60°dshsR
  X1 = AAMM_BUFFON_calcul_X_ptA_ds_hsR(bo1_rd) 'bo1_rd = teta2_hsR = +60°+30°

  If loiBX("X") <= X1 Then

      vd = "OK5" '"OK" si on veut editer ds Debug.print
      If i = 0 And rangM = 1 And t = t_test Then vd = "OK2"

     'X2 (bo2_teta) = positon endGP epres raccsst: = -40°dshsR
      X2 = AAMM_BUFFON_calcul_X_ptA_ds_hsR(bo2_rd) 'bo1_rd = - 17°-30°

      If loiBX("X") >= X2 Then 'ok c'est bon : X2_bo2_rd(finGP) <= XsitA_loiBX("X")<= X1_bo1_rd(debGP)

          ' F = compteur de possibilites de CB dans loiBV sachant qu'on en retiendra qu'une ds loiBA
          F = F + 1

         'etap 1: on interpole linéairement ? calcul + précis ?
         'ATTENTION ymi interploé linéairement en degré
         ymi = bo2_dg + (loiBX("X") - X2) * (bo1_dg - bo2_dg) / (X1 - X2) 'en deg

         'etap 2: on verifie et si pas bon on incrémente
         vd = "OK17" 'oK7 pour enregistrer la linearisation
        ' If t = 4 Then vd = "OK7"

         i_yt = 0
         For Rd_ou_Deg = 1 To 2 ' rd=1 deg=2

             yap = ymi * IIf(Rd_ou_Deg = 1, pi / 180, 1) '1=en rd 2=en deg et c'est la ou tombe sur les barres
             yt = AAMM_BUFFON_calcul_X_ptA_ds_hsR(yap) - loiBX("X") 'en nm

             If Abs(yt) > 0.1 Then ' tolerance 0.07 nm sx/10 correspondant approximativement à 0.5°
                signX = IIf(yt > 0, -1, 1)
                pas_A = 0.0007 * IIf(Rd_ou_Deg = 1, 1, 10) 'en rd

                For i_yt = 1 To 1000 '1°=0,01745 r d
                    yap = yap + pas_A * signX 'on incremente de pas_A
                    yt = AAMM_BUFFON_calcul_X_ptA_ds_hsR(yap) - loiBX("X")

                    '***********VERIF INTERPOLATION****************************************************
                    If vd = "OK7" Then 'on enregistre pour verifier
                       If Rd_ou_Deg = 1 Then
                         drap = 1
                       Else
                          loiBV.Seek "=", i + 1, rangM, F, i_yt
                          drap = IIf(loiBV.NoMatch, 1, 2)
                       End If

                       If drap = 1 Then 'on crée
                          loiBV.AddNew
                            loiBV("no_courbe") = i + 1
```

```
                loiBV("pasX") = i * sX
                loiBV("rangM") = rangM
                loiBV("zonM60") = zonM60(B)
                loiBV("entre_2_zonM") = entre_2_zonM
                loiBV("Psi") = xmi
                loiBV("ALPHA_S2") = ALPHA_S2 'par defaut
                loiBV("no_A") = F
                loiBV("i_yt") = i_yt

                loiBV("sitA") = loiBX("no")
                loiBV("tetM16") = t
                loiBV("Beta_360_sitA") = Int(yav * 10 + 0.5) / 10
                loiBV("Beta_PouM_60_sitA") = Int(BeTA * 10 + 0.5) / 10
                loiBV("X_sitA") = loiBX("X")
                loiBV("Alin") = Int(ymi * 10 + 0.5) / 10
                loiBV("X_ptAlin") = AAMM_BUFFON_calcul_X_ptA_ds_hsR(ymi * pi / 180) 'rappel ymi en deg


              Else 'deja crre
                loiBV.Edit

              End If

              'on complete
              If Rd_ou_Deg = 1 Then
                loiBV("A") = Int(yap * 180 / pi * 100 + 0.5) / 100 'rappel yap en rd donc on repssa en deg
                loiBV("A_N_interp") = i_yt
                loiBV("X_ptA") = AAMM_BUFFON_calcul_X_ptA_ds_hsR(yap)
                 loiBV("diffX_ptA") = loiBV("X_sitA") - loiBV("X_ptA")
              Else
                loiBV("Ac_en_deg") = Int(yap * 100 + 0.5) / 100 'yap en deg
                loiBV("Ac_en_deg_N_interp") = i_yt
                loiBV("X_ptAc_en_deg") = AAMM_BUFFON_calcul_X_ptA_ds_hsR(yap)
                loiBV("diffX_ptAc_en_deg") = loiBV("X_sitA") - loiBV("X_ptAc_en_deg")
              End If
             loiBV.Update
            End If 'ok7
            '***********FIN VERIF INTERPOLATION***************************************************

            If Abs(yt) < 0.1 Then Exit For 'i_yt
          Next 'i_yt

          'Debug.Print "k = " + Format(k), "Teta/ymi = " + Format(Int(ymi * 10 + 0.5) / 10) + "°",  "Teta/yap = " + Format(Int(yap *
IIf(Rd_ou_Deg = "deg", 1, 180 / pi) * 10 + 0.5) / 10) + "°", "yt = " + Format(yt) + " nm", "no_sitA=" +  Format(loiBX("no"))
          If i_yt > 1000 Then Stop

        End If

      'on memorise repassae en degré
      If Rd_ou_Deg = 1 Then 'rd
        teta_interpol = yap * 180 / pi
        A_N_interp = i_yt
      'else
      ' dans l'autre cas on garde yap tel quel
      End If
    Next 'Rd_ou_Deg

    If i_yt > 0 And vd = "OK7" Then Stop

      loiBV.AddNew
        loiBV("no_courbe") = i + 1
        loiBV("pasX") = i * sX
```

```
            loiBV("rangM") = rangM
            loiBV("zonM60") = zonM60(B)
            loiBV("entre_2_zonM") = entre_2_zonM
            loiBV("Psi") = xmi
            loiBV("ALPHA_S2") = ALPHA_S2 'par defaut
            loiBV("no_A") = F
            loiBV("i_yt") = 1

            loiBV("Alin") = Int(ymi * 100 + 0.5) / 100
            loiBV("A") = Int(teta_interpol * 100 + 0.5) / 100
            loiBV("A_N_interp") = A_N_interp
            loiBV("Ac_en_deg") = Int(yap * 100 + 0.5) / 100
            loiBV("Ac_en_deg_N_interp") = i_yt
            loiBV("Ac_en_deg_mult_6point3") = Int(yap * 100 / (2 * pi) + 0.5) / 100 '6.3=2*pi
            loiBV("sitA") = loiBX("no")
            loiBV("tetM16") = t
            loiBV("Beta_360_sitA") = Int(yav * 10 + 0.5) / 10
            loiBV("Beta_PouM_60_sitA") = Int(BeTA * 10 + 0.5) / 10
            loiBV("X_sitA") = loiBX("X")
            loiBV("X_ptA") = AAMM_BUFFON_calcul_X_ptA_ds_hsR(yap * pi / 180)
            loiBV("X_ptAlin") = AAMM_BUFFON_calcul_X_ptA_ds_hsR(ymi * pi / 180)
            loiBV("X_ptAc_en_deg") = AAMM_BUFFON_calcul_X_ptA_ds_hsR(ymi * pi / 180)

            loiBV("diffX_ptA") = loiBV("X_sitA") - loiBV("X_ptA")
            loiBV("diffX_ptAc_en_deg") = loiBV("X_sitA") - loiBV("X_ptAc_en_deg")
          loiBV.Update

    End If 'borne X2
End If 'borne X1


    End If



End Sub
```

#### Function AAMM_BUFFON_calcul_X_ptA_ds_hsR(angle_Teta_rd As Single) As Single

```
Dim yxx As Single
 LS2_p = Sqr(LS2 ^ 2 - ((r_filA + YS1a + LS1b * Cos(angle_Teta_rd)) * Sin(BeTArd) + r_filM * Sin(ALPHA_S2rd)) ^ 2)
 'dans le calcul de l'annexe l'angle Phi est negatif
 yxx = d_filAMc - (r_filA + YS1a + LS1b * Cos(angle_Teta_rd)) * Cos(BeTArd) - r_filM * Cos(ALPHA_S2rd) '>0

 FI1 = ASINr(-yxx / LS2_p) 'en rd
 AAMM_BUFFON_calcul_X_ptA_ds_hsR = Xd + XS1a + LS1b * Sin(angle_Teta_rd) + LS2_p * Cos(FI1)

End Function
```

#### Function ASINr(xt As Single) As Single 'donne angle en radient

```
ASINr = xt + (xt ^ 3) / 6 + (xt ^ 5) * 3 / 40 + (xt ^ 7) * 5 / 112 + (xt ^ 9) * 35 / 1152 + (xt ^ 11) * 63 / 2816 + (xt ^ 13) * 231 / 13312

End Function
```

#### Sub AAMM_BUFFON_angle_Teta_trace_Angle()

```
Dim presG As Integer, y_moyG As Single, fact_fG As Single
presG = 2 '1=presentation en col 2=presentation alternatic D ou G
comm = 0 '0=ss_comm // 1=av_comm

If loiBA.RecordCount = 0 Then 'atble pas remplit
   vd = MsgBox("Remplit d'abord la table", 0, "Triple Buse !")
   Obuff(3) = -1
   Exit Sub
 End If

'et maintenant on trace DX=DTeTa*R_WS*LS1b
' rappel : 11nm correspond à 70°soit 1nm = 6.4°

' la distributuion sur 50°soit 7.85 nm en fait 49°  soit 7.7 nm
 vx = "angle Téta (°)"
 'xt = 5
 'xn = Int(bo2_dg / xt - 0.01) * xt ' Teta2= -17.5°  Teta3=-37.4°
 'xx = Int(bo1_dg / xt - 0.01) * xt + xt 'Teta1=+30° Teta_S=55°
 xt = 10
 xn = -42
 xx = 28

If FrBUFF.Tag = 4 Then 'dénombrment
   vy = "N": yn = 0: yx = 7: yt = 1

Else 'gaussiennes

   Np4 = 100
   D = Val(TAM(17).Text) 'ecart-type SD=6.1°correspon d à une deplacement 1nm
   SDg = 9.5 '10 '°ecart-typr modele gaussien 22°p  our corrie 1999
   AAMM_BUFFON_angle_Teta_trace_Angle_caluls_AIRES

   'v = 45 / ((xx_IT - xn_IT) * Atn(1)) calculée dans  AAMM_BUFFON_angle_Teta_trace_Angle_caluls_AIRES
   'v=0.82 = dU :valeur loi uniforme sur 70°
   'v= 1.15 = dU :valeur loi uniforme sur 50°

   'loiBG.Seek "=", 13
   'y_moyG = loiBG("Moy_Aire")

   vy = "P": yn = 0: yt = 0.5
   yx = IIf(Frcond.Tag = 0 Or Frcond.Tag = 2, 2, 3)

   'fact_fG = v / y_moyG

End If

For i = 0 To 11 'i est le no de la courbe de 0 à 11

    loiBG.Seek "=", i
    fact_fG = loiBG("1/Aire")

    sYY = 0 'conpteur de WS ss pois stat pour chaque deplaclment
    sXY = 0 'conpteur de wS av poids stat pour chaque deplaclment
    AAMM_BUFFON_angle_Teta_trace_Angle_scale presG '1=en col 2=alternatic D ou G

    'titre
    With O
    If comm = 1 Then

         .FontSize = 14: .FontBold = -1
         .CurrentX = 80: .CurrentY = 115
        Select Case drap
        Case 1
           loiBO.MoveFirst
           O.Print Oesc(Fresc.Tag).Caption
        Case 2: O.Print "d_filAM = " + TAM(15).Text + " nm"
        Case 3: O.Print "ds hsR: bo1_Teta = " + TAM(6).Text + "° (Teta1 = " + Format(Int(Teta1 * 1800 /  pi + 0.5) / 10) + "°)"
        Case 4: O.Print "ds hsR: bo2_Teta = " + TAM(7).Text + "° (Teta2 = " + Format(Int(Teta2 * 1800 /  pi + 0.5) / 10) + "°)"
        Case 5: O.Print "d_bo = " + Format(Val(TAM(6).Text) - Val(TAM(7).Text)) + "° (dTeta =  " + Forma t(Int((Teta1 - Teta2) * 1800
/ pi + 0.5) / 10) + "°)"
```

```
      'Case 12  'total CB
      '    .CurrentX = -30: .CurrentY = 115
      '     O.Print "TOTAL CB pour les 9 lignes 16_tetM = " + Format(sXY)
      End Select
   End If

   .FontSize = 10: .FontBold = 0: .ForeColor = QBColor(0)
   .CurrentX = 30: .CurrentY = 117
   .FontBold = -1
   O.Print "RACCOURCISSEMENT de " + Format((i)) + " nm"  '  (TOTAL CB pour les 9 lignes 16_tetM = " + Format(sXY) + ")"
     If i = 1 Then
      .FontSize = 6: .CurrentX = 2: .CurrentY = 107
      O.Print "X0_ptD_depart =" + Format(X0_ptD_depart) + "nm"
    End If
    .FontBold = 0
   End With

   impenvi
   '2eme echelle
   'bo1_nm = LS1b * R_WS * (Teta0 - bo1_rd): bo1_nm = Int(bo1_nm * 10 + 0.5) / 10 'b=borne_X_preWS
   'bo2_nm = LS1b * R_WS * (Teta0 - bo2_rd): bo2_nm = Int(bo2_nm * 10 + 0.5) / 10
    npas = (xx - xn) / 10
   For z = xn / 10 To npas + xn / 10
      xav = (z * 10 - xn) * 100 / (xx - xn)
      O.Line (xav, 0)-(xav, -1)
   Next

   '                                    WS_end          WS_start        SB_start
   If comm = 1 And FrBUFF.Tag = 4 Then 'GROS TRAITS  Tetat3_HSR  Teta2_HSR Teta1_HSR  TetaF_HSR DS
TABLE8BUFFON8RWS
      O.DrawWidth = 6
      For z = 1 To 4 ' teta1 teta2 teta3 tetaF
         vd = IIf(z = 4, "F", Format(z)) 'F=First ou 1ere valeur d'attachement SB
         xav = (loiBR("teta" + vd + "_hsr") - xn) * 100 / (xx - xn)
         O.Line (xav, 10)-(xav, -10), QBColor(IIf(z = 1 Or z = 2, 2, 9)) '2=vert 9=bleu clair
      Next
   End If


   'on trace
    'petit police
   vd = O.FontName: O.FontName = Screen.Fonts(7): O.FontSize = 8 '7
   O.FontBold = -1
   If FraX.Tag < 3 Then FraX.Tag = 3 'au cas où
   ' pas de i ni t ni z
   If FrBUFF.Tag = 4 And FraX.Tag = 6 Then '1 seule ligne
       F = Val(TAM(16).Text) 'n°ligne
        AAMM_BUFFON_angle_Teta_trace_Angle2 F, "les 2"
   Else 'toutes les lignes
      For F = 1 To 9
         AAMM_BUFFON_angle_Teta_trace_Angle2 F, "les 2"
      Next
   End If

  O.FontName = vd

  If FrBUFF.Tag = 5 Then 'loi uniforme ?
     ' If Frcond.Tag = 1 Or Frcond.Tag = 3 Then 'le facteur sear donc Y *vlaen 21/valeur en 20
     '    loiBG.Seek "=", 20: r = v / loiBG("aire")
     ' End If

     'pt de départ
       loiBG.Seek "=", 0
       O.DrawWidth = 3
       O.CurrentX = (xn_IT - xn) * 100 / (xx - xn)
       yav = loiBG("sG_" + Format(i)) * fact_fG
       If Frcond.Tag = 1 Or Frcond.Tag = 3 Then 'on calul les ecats
          loiBG.Edit
            loiBG("Ecart_LU") = Abs(yav - v)
            loiBG("Ecart_LG") = Abs(yav - loiBG("LG") * r)
           loiBG.Update
```

```
 End If
 O.CurrentY = (yav - yn) * 100 / (yx - yn)

 For z = 1 To Np4 '100
    xap = xn_IT + z * (xx_IT - xn_IT) / Np4

    loiBG.Seek "=", z
    yav = loiBG("sG_" + Format(i)) * fact_fG
    If Frcond.Tag = 1 Or Frcond.Tag = 3 Then 'on calul les ecats
       loiBG.Edit
         loiBG("Ecart_LU") = Abs(yav - v)
         loiBG("Ecart_LG") = Abs(yav - loiBG("LG") * r)
       loiBG.Update
    End If

    xap = (xap - xn) * 100 / (xx - xn)
    yap = (yav - yn) * 100 / (yx - yn)
    O.Line -(xap, yap) ', coul

 Next 'z
'-------------------- on figonle en traçant la valeur moyenne
 O.DrawMode = 9
 O.ForeColor = QBColor(12)

If Frcond.Tag = 1 Or Frcond.Tag = 3 Then
   bo1_dg = Val(TAM(6).Text)
   bo2_dg = Val(TAM(7).Text)
Else
   bo1_dg = xn
   bo2_dg = xx
End If
xav = (bo1_dg - xn) * 100 / (xx - xn)
xap = (bo2_dg - xn) * 100 / (xx - xn)


'dU on ecrit densité mmoy vlaeur moy 0.573 et 1.273
If i = 20 Then
   O.CurrentX = 5: O.CurrentY = 100
   O.Print "p_moy = " + Format(v, "0.###")
   O.CurrentX = 5: O.CurrentY = 90
   loiBG.Seek "=", 14
   O.Print "p_SD = " + Format(loiBG("Moy_Aire_Norm"), "0.###")
End If

ymi = (v - yn) * 100 / (yx - yn)
       'O.DrawStyle = 0
O.DrawWidth = 2
O.Line (xav, 0)-(xav, ymi), QBColor(12)
O.Line (xap, 0)-(xap, ymi), QBColor(12)
O.DrawWidth = 5
O.Line (xav, ymi)-(xap, ymi), QBColor(12)

'd  aire rapporté à 100 en normalisant par rapport à densite moyenne
O.CurrentX = 85: O.CurrentY = 100
'loiBG.Seek "=", i
'ymi = loiBG("aire_Norm")
O.Print "1/A" + Format(i) + " = " + Format(fact_fG, "0.###")
ymi = (ymi - yn) * 100 / (yx - yn)
'O.DrawWidth = 1: O.DrawStyle = 2
'O.Line (xav, ymi)-(xap, ymi), QBColor(12)


If Frcond.Tag = 1 Or Frcond.Tag = 3 Then 'le facteur sear donc Y *vlaen 21/valeur en 20
   O.DrawWidth = 3

  ' loiBG.Seek "=", 21: r = loiBG("aire")
  loiBG.Seek "=", 20: r = v / loiBG("Moy_aire") 'facteur 1/aire

   O.CurrentX = (xn_IT - xn) * 100 / (xx - xn)
   loiBG.Seek "=", 0
   yav = loiBG("LG") * r
```

```
        O.CurrentY = (yav - yn) * 100 / (yx - yn)
        sX = loiBG("Ecart_LU")
        sY = loiBG("Ecart_LG")
        For z = 1 To Np4 '100
            xav = xn_IT + z * (xx_IT - xn_IT) / Np4
            loiBG.Seek "=", z
            yav = loiBG("LG") * r
            xap = (xav - xn) * 100 / (xx - xn)
            yap = (yav - yn) * 100 / (yx - yn)
            O.Line -(xap, yap), QBColor(10) 'QBColor(10)
            sX = sX + loiBG("Ecart_LU")
            sY = sY + loiBG("Ecart_LG")
         Next

       'on enregistre les ecarts
        loiBG.Seek "=", i
        loiBG.Edit
            loiBG("Moy_ect_LU") = sX / 12
           loiBG("Moy_ect_LG") = sY / 12
         loiBG.Update

      End If
  End If

  '*** BORNES ROUGES
   'on trace bo1_dg= Val(TAM(6).Text) et bo2_dg= Val(TAM(7).Text)
   O.DrawWidth = 4
   If FrBUFF.Tag = 4 Or Frcond.Tag = 1 Or Frcond.Tag = 3 Then
      bo1_dg = Val(TAM(6).Text)
      bo2_dg = Val(TAM(7).Text)
   Else
      bo1_dg = xn
      bo2_dg = xx
   End If
   xav = (bo1_dg - xn) * 100 / (xx - xn)
   xap = (bo2_dg - xn) * 100 / (xx - xn)


  '''O.DrawWidth = 4
  O.DrawWidth = 1: O.DrawStyle = 2
  'If FrBUFF.Tag = 5 And (Frcond.Tag = 1 Or Frcond.Tag = 3) Then 'Gaussiennes
  If FrBUFF.Tag = 5 Then 'Gaussiennes
     'on recupere l'aire sous les gaussiennes entre xn et xx
     'loiBG.Seek "=", i
     'ymi = loiBG("aire") * fact_fG   '* IIf(Frcond.Tag = 1 Or Frcond.Tag = 3, (xx - xn) / (bo1_dg - bo2_dg), 1)

     'on trace l'aire
     'O.Line (xav, 0)-(xav, ymi), QBColor(12)
     'O.Line (xap, 0)-(xap, ymi), QBColor(12)
     'O.Line (xav, ymi)-(xap, ymi), QBColor(12)

  ElseIf comm = 1 Then
      ymi = 60
      O.Line (xav, 0)-(xav, ymi), QBColor(12)
      O.Line (xap, 0)-(xap, ymi), QBColor(12)

  End If

 If FrBUFF.Tag = 4 Then
    With O
    .FontSize = 10: .ForeColor = QBColor(0): .FontBold = -1
    'nombre
     .CurrentX = 30: O.CurrentY = 103
     O.Print "TOTAL SB_9rangM = " + Format(sYY) + " ( "; Format(sXY, "##.#") + "_av pds stat)" 'poids stat pour tenir compte du
nombre de position pour chaque tetM
     S1 = S1 + sXY
     If FraX.Tag < 6 Then '% rappel 16 x 9 = 144
        .CurrentX = 30: O.CurrentY = 90
        O.Print " (% XS_9rangM = " + Format(sXY / 288, "0.##") + "_av pds stat)" 'poids stat pour tenir compte du nombre de position
pour chaque tetM
        S2 = S2 + sXY / 144
```

```
    End If
    .FontBold = 0
   End With
 End If

Next 'i deplcements

'TOUCHE FINALE
If FrBUFF.Tag = 4 Then 'on figonle en donnant val .moyenne
     O.ForeColor = QBColor(12)
     O.FontBold = -1
     O.CurrentX = 68
     O.CurrentY = 85
     O.Print "MOY_TOT_av pds stat = " + Format(S1 / 12, "##.#")
     If FraX.Tag < 6 Then '% rappel 16 x 9 = 144
        O.CurrentX = 68: O.CurrentY = 70
        O.Print "MOY_%SB_av pds stat = " + Format(S2 / 12, "0.###")
    End If

ElseIf FrBUFF.Tag = 5 And (Frcond.Tag = 1 Or Frcond.Tag = 3) Then
      sX = 0: sY = 0
      loiBG.MoveFirst
      Do While loiBG("pt") <= 11
         sX = sX + loiBG("Moy_Ect_LU") / 12
         sY = sY + loiBG("Moy_Ect_LG") / 12
         loiBG.MoveNext
      Loop

      loiBG.Seek "=", 13
      loiBG.Edit
          loiBG("Moy_Ect_LU") = sX
          loiBG("Moy_Ect_LG") = sY
      loiBG.Update

    O.CurrentX = 5: O.CurrentY = 100
    O.Print "SD_Gauss = " + Format(SDg) + "°"
   'loiBG.Seek "=", 13
    O.CurrentX = 5: O.CurrentY = 90
    O.Print "MOY ects/LU = " + Format(sX, "0.###")
    O.CurrentX = 5: O.CurrentY = 80
    O.Print "MOY ects/LG = " + Format(sY, "0.###")

End If

End Sub
```

### Sub AAMM_BUFFON_angle_Teta_trace_Angle_caluls_AIRES()

```
If FrBUFF.Tag <> 5 Then Stop
xn_IT = IIf(Frcond.Tag = 1 Or Frcond.Tag = 3, bo2_dg, xn)
xx_IT = IIf(Frcond.Tag = 1 Or Frcond.Tag = 3, bo1_dg, xx)
v = 45 / ((xx_IT - xn_IT) * Atn(1)) '1/D_tetamax ou 1/dtetasatrtXMAX

If loiBG.RecordCount > 0 Then dbM.Execute "DELETE * FROM " + zYb 'loiBA

For i = 0 To 11 'nm depalcments
  'etpe 0 : remise à 0
    For z = 0 To Np4: sG(z) = 0: Next

  'etape1 : calcul
    For F = 1 To 9 'Mrow

        loiBO.Seek ">=", Choose(Frcond.Tag + 1, 0, 1, 0, 1), i, F, -90 'i est le no de la courbe de 0 à 11 // F =n°rangM_16tetM // -90
valeur angle minimale
        If loiBO.NoMatch Then Exit Sub

        Do While loiBO("pasX") = i And loiBO("rangM") = F
            xav = loiBO("A_val")
            Pdp = IIf(Frcond.Tag < 2, 1, 1 / loiBO("sitA8"))
            ' rappel LG f(x)= exp[-0.5x(x-M)²/SD²] / [SD*sqr(2pi)]      'attention il faut tout calculer en radiant
            ' rappel :   D = Val(TAM(17).Text)         'ecart-type

            'pt de départ
             xmi = -0.5 * ((xn_IT - xav) / D) ^ 2  'rappel: xav = loiBO("A_val")
            ' la pas de pb en reste en nm  puis
             yap = Exp(xmi) / (D * Sqr(2 * pi))  'D=7°(= 0.9 nm) car variation angulaire d_teta= dx/(ls1 bxR_WS) mais on s'en fout car à
une constante pres que ce soit en °, en rad ou en n m
             sG(0) = sG(0) + Pdp * Exp(xmi) / (D * Sqr(2 * pi))

            For z = 1 To Np4 '100
                xap = xn_IT + z * (xx_IT - xn_IT) / Np4
                xmi = -0.5 * ((xap - xav) / D) ^ 2 'xav = loiBO("A_val") =val moy de la Gaussienne
                yap = Exp(xmi) / (D * Sqr(2 * pi)) ' D = Val(TAM(17).Text) 'ecart-type
                sG(z) = sG(z) + Pdp * Exp(xmi) / (D * Sqr(2 * pi)) 'Pdp = IIf(Frcond.Tag < 2, 1, loiBO("poids_stat"))
            Next
            loiBO.MoveNext
            If loiBO.EOF Then Exit Do
        Loop
    Next 'F

  'etape 2 : on stocke
    For z = 0 To Np4
        If i = 0 Then
           loiBG.AddNew
           loiBG("pt") = z
        Else
           loiBG.Seek "=", z
           loiBG.Edit
        End If
           loiBG("sG_" + Format(i)) = sG(z)
        loiBG.Update
    Next

   'etep 4: integer l'aire sous les gaussiennes entre xn et xx
      sX = 0: sY = sG(0)
      For z = 1 To Np4 '100
         sX = sX + (sG(z - 1) + sG(z)) / 2
         sY = sY + sG(z)
      Next
    'etape 5 on stocke
      loiBG.Seek "=", i
      loiBG.Edit
      loiBG("Moy_Aire") = sX / Np4
      loiBG("1/Aire") = (Np4 * v) / sX 'L=1/v car  v = 45 / ((xx_IT - xn_IT) * Atn(1)) = 1/L = 1/D_tetamax ou 1/dtetasatrtXMAX
      loiBG("Aire") = sX / (Np4 * v)
      loiBG("Moy_arith") = sY / Np4
      loiBG.Update
```

```
Next 'i

'derniere etape
sXY = 0: pT = 0
loiBG.MoveFirst
Do While loiBG("pt") <= 11
   sXY = sXY + loiBG("Moy_aire") / 12
   pT = pT + loiBG("Moy_arith") / 12
   loiBG.MoveNext
Loop
loiBG.Seek "=", 13
loiBG.Edit
    loiBG("Moy_Aire") = sXY
    loiBG("Moy_arith") = pT
loiBG.Update

'normalisée
sX = 0: sXX = 0: sY = 0: sYY = 0
loiBG.MoveFirst
Do While loiBG("pt") <= 11
   pT1 = loiBG("Moy_Aire") * v / sXY
   pTi = loiBG("MOy_arith") * v / pT
   loiBG.Edit
    loiBG("Moy_Aire_Norm") = pT1
    loiBG("Moy_arith_Norm") = pTi
   loiBG.Update
   'Aire
   sX = sX + pT1
   sXX = sXX + pT1 ^ 2
   'MOY
   sY = sY + pTi
   sYY = sYY + pTi ^ 2


   loiBG.MoveNext
Loop

'les moyennes
loiBG.Seek "=", 13 'on doit retrouber v
loiBG.Edit
    loiBG("Moy_Aire_Norm") = sX / 12
    loiBG("Moy_arith_Norm") = sY / 12
loiBG.Update

'les SD
loiBG.Seek "=", 14
loiBG.Edit
    loiBG("Moy_Aire_Norm") = Sqr(sXX - sX ^ 2 / 12) / Sqr(11)
    loiBG("Moy_arith_Norm") = Sqr(sYY - sY ^ 2 / 12) / Sqr(11)
loiBG.Update


'--------------------on teste la gaussienne ' rappel LG f(x)= exp[-0.5x(x-M)²/SD²] / [SD*sqr(2pi)]
 If Frcond.Tag = 1 Or Frcond.Tag = 3 Then
    'sdg = 20 'corrie 1999 22.1°
    Teta0 = (xx_IT + xn_IT) / 2
     For z = 0 To Np4 '100
         xav = xn_IT + z * (xx_IT - xn_IT) / Np4
         xmi = -0.5 * ((xav - Teta0) / SDg) ^ 2
         sG(z) = Exp(xmi) * 45 / (SDg * pi * Sqr(2 * pi))

         loiBG.Seek "=", z
           loiBG.Edit
           loiBG("LG") = sG(z)
        loiBG.Update
     Next

   'etep 4: integer l'aire sous les gaussiennes entre xn et xx
      sX = 0
      For z = 1 To Np4 '100
          sX = sX + (sG(z - 1) + sG(z)) / 2
```

```
      Next
    'etape 5 on stocke
      loiBG.Seek "=", 20
      loiBG.Edit
        loiBG("Moy_Aire") = sX / Np4
        loiBG("Aire") = sX / (Np4 * v)
         loiBG("1/Aire") = (Np4 * v) / sX
       loiBG.Update 'moyenne
       loiBG.Seek "=", 21
      loiBG.Edit: loiBG("Moy_Aire") = sX * (xx_IT - xn_IT) / Np4: loiBG.Update 'normalisé
      'le facteur sear donc Y *vlaen 21/valeur en 20

End If

End Sub
```

#### Sub AAMM_BUFFON_angle_Teta_trace_Angle_scale(iX As Integer)

```
drap = i + 1

If iX = 1 Then
   'de  0 1 2 3 4 5 nm sur col G + 'de 6 7 8 9 10 11 sur col D

   Select Case drap
      Case 1: impscale X0, Y0 + haut + hentete, X0 + 2 * larg, Y0 - 5 * haut - hpage
      Case 2: impscale X0, Y0 + 2 * haut + hentete, X0 + 2 * larg, Y0 - 4 * haut - hpage
      Case 3: impscale X0, Y0 + 3 * haut + hentete, X0 + 2 * larg, Y0 - 3 * haut - hpage
      Case 4: impscale X0, Y0 + 4 * haut + hentete, X0 + 2 * larg, Y0 - 2 * haut - hpage
      Case 5: impscale X0, Y0 + 5 * haut + hentete, X0 + 2 * larg, Y0 - haut - hpage
      Case 6: impscale X0, Y0 + 6 * haut + hentete, X0 + 2 * larg, Y0 - hpage

      Case 7: impscale X0 - larg, Y0 + haut + hentete, X0 + larg, Y0 - 5 * haut - hpage
      Case 8: impscale X0 - larg, Y0 + 2 * haut + hentete, X0 + larg, Y0 - 4 * haut - hpage
      Case 9: impscale X0 - larg, Y0 + 3 * haut + hentete, X0 + larg, Y0 - 3 * haut - hpage
      Case 10: impscale X0 - larg, Y0 + 4 * haut + hentete, X0 + larg, Y0 - 2 * haut - hpage
      Case 11: impscale X0 - larg, Y0 + 5 * haut + hentete, X0 + larg, Y0 - haut - hpage
      Case 12: impscale X0 - larg, Y0 + 6 * haut + hentete, X0 + larg, Y0 - hpage

    End Select

 Else
  'de  0 2 4 6 8 10 nm sur col G + 'de 1 3 5 7 9 11 nm sur col D

    Select Case drap

      Case 1: impscale X0, Y0 + haut + hentete, X0 + 2 * larg, Y0 - 5 * haut - hpage
      Case 3: impscale X0, Y0 + 2 * haut + hentete, X0 + 2 * larg, Y0 - 4 * haut - hpage
      Case 5: impscale X0, Y0 + 3 * haut + hentete, X0 + 2 * larg, Y0 - 3 * haut - hpage
      Case 7: impscale X0, Y0 + 4 * haut + hentete, X0 + 2 * larg, Y0 - 2 * haut - hpage
      Case 9: impscale X0, Y0 + 5 * haut + hentete, X0 + 2 * larg, Y0 - haut - hpage
      Case 11: impscale X0, Y0 + 6 * haut + hentete, X0 + 2 * larg, Y0 - hpage

      Case 2: impscale X0 - larg, Y0 + haut + hentete, X0 + larg, Y0 - 5 * haut - hpage
      Case 4: impscale X0 - larg, Y0 + 2 * haut + hentete, X0 + larg, Y0 - 4 * haut - hpage
      Case 6: impscale X0 - larg, Y0 + 3 * haut + hentete, X0 + larg, Y0 - 3 * haut - hpage
      Case 8: impscale X0 - larg, Y0 + 4 * haut + hentete, X0 + larg, Y0 - 2 * haut - hpage
      Case 10: impscale X0 - larg, Y0 + 5 * haut + hentete, X0 + larg, Y0 - haut - hpage
      Case 12: impscale X0 - larg, Y0 + 6 * haut + hentete, X0 + larg, Y0 - hpage

     End Select
End If

End Sub
```

#### Sub impscale(X1 As Integer, y1 As Integer, X2 As Integer, Y2 As Integer)

```
O.ScaleLeft = X1
O.ScaleTop = y1
O.ScaleWidth = X2 - X1
O.ScaleHeight = Y2 - y1
End Sub
```

#### Sub AAMM_BUFFON_angle_Teta_trace_Angle2(iR As Integer, G_D As String)

```
Dim fact_Gauss As Single

If FrBUFF.Tag = 5 Then '
   xn_IT = IIf(Frcond.Tag = 1 Or Frcond.Tag = 3, bo2_dg, xn)
   xx_IT = IIf(Frcond.Tag = 1 Or Frcond.Tag = 3, bo1_dg, xx)
End If
'                        1   2   3   4   5  6  7  8  9
'coul = QBColor(Choose(iR, 9, 13, 11, 12, 10, 8, 2, 4, 1))
coul = QBColor(Choose(iR, 10, 13, 11, 12, 9, 8, 2, 4, 1))

O.ForeColor = coul
loiBO.Seek ">=", Choose(Frcond.Tag + 1, 0, 1, 0, 1), i, iR, -90 'i est le no de la courbe de 0 à 11 // iR =n°rangM_16tetM // -90
valeur angle minimale

'loiBO.Seek "=", 0, i, loiBA("A")loiBO.Seek "=", 0, i, loiBA("A")

'rappel : la differencre entre xn=Teta_DOwn et xx=Teta_UP  et Teta1 et eta2 a ete fait donc sub_routine BUFFON_stocke
'           le calcue des gaussiennnese se fait entre les bornes xn=Teta_DOwn et xx=Teta_UP

If loiBO.NoMatch Then Exit Sub

'---bornes sur lesquelles on va travailler pour les conditions rappel : frcond.tag=0 tout 1cond
'bo1_Beta = Val(TAM(1).Text) 'bo1_Beta >=0
'bo2_Beta = Val(TAM(2).Text) 'bo2_Beta <60°
'bo1_Alpha = Val(TAM(3).Text) 'bo1_Alpha =0
'bo2_Alpha = Val(TAM(4).Text) 'bo2_Alpha > - 30°
'bo1_dg = Val(TAM(6).Text) 'borne_Teta1
'bo2_dg = Val(TAM(7).Text)'borne_Teta2
O.DrawWidth = 1.5
t = 0 'set à position les etiquettes des sitA
Do While loiBO("pasX") = i And loiBO("rangM") = iR

   If G_D = "les 2" Then  ' Or (G_D = "G" And loiBO("courb") < 7) Or (G_D = "D" And loiBO("courb") > 6) Then 'oK
      xav = loiBO("A_val")
      xap = (xav - xn) * 100 / (xx - xn)

      '**************
      If FrBUFF.Tag = 4 Then 'angle + etiquettes des sitA

        sN = loiBO("N_sitA")
        npas = 0

        For z = 1 To sN
           If comm = 1 Or FraX.Tag = 6 Then
              O.CurrentX = xap '+ IIf(Int(t / 2) = t / 2, -2, 0)
              O.CurrentY = (sN + z * 0.7 - yn) * 100 / (yx - yn)
              O.Print Format(loiBO("sitA" + Format(z)))
           End If
           npas = npas + 1

        Next

        'petite verif
        'If Frcond.Tag = 0 And npas <> sN Then Stop
        'trait si compteur >0
        If npas > 0 Then

         sYY = sYY + loiBO("N_sitA") 'ss poids stat
         'sXY = sXY + loiBO("N_sitA") * loiBO("poids_stat") 'poids stat pour tenir compte du nombre de position pour chaque tetM
         sXY = sXY + loiBO("N_sitA") / loiBO("sitA8")

         yap = (npas - yn) * 100 / (yx - yn)
           O.Line (xap, 0)-(xap, yap), coul
        End If

     '**********
     Else 'on trace les gaussiennes aturour de chaque position angulaire recensées et on en profite pour calculer la loi Uniforme
somme de toutes ces gaussiennes
     ' rappel LG f(x)= exp[-0.5x(x-M)²/SD²] / [SD*sqr(2pi)]
```

```
        'attention il faut tout calculer en radiant

        'Pdp = IIf(Frcond.Tag < 2, 1, loiBO("poids_stat"))
        Pdp = IIf(Frcond.Tag < 2, 1, 1 / loiBO("sitA8"))
        fact_Gauss = Pdp * 6 * IIf(Frcond.Tag = 0 Or Frcond.Tag = 2, 1, 7 / 5)
        '4

        'A_teta = moy au centre de LG
        yav = 0.5 * IIf(Frcond.Tag = 0 Or Frcond.Tag = 2, 1, 7 / 5)
        yap = (yav - yn) * 100 / (yx - yn)
        O.Line (xap, 0)-(xap, yap), coul

        'la gaussienne' rappel :  D = Val(TAM(17).Text) 'ecart-type
          'pt de départ
          loiBG.Seek "=", 0
          O.DrawWidth = 1
          O.CurrentX = (xn_IT - xn) * 100 / (xx - xn)
          xmi = -0.5 * ((xn_IT - xav) / D) ^ 2 'rappel: xav = loiBO("A_val")
          ' la pas de pb en reste en nm  puis
          yap = Exp(xmi) / (D * Sqr(2 * pi))  'D=7°(= 0.9 nm) car variation angulaire d_teta= dx/(ls1 bxR_WS) mais on s'en fout car à
une constante pres que ce soit en °, en rad ou en n m
          'sG(0) = sG(0) + Pdp * Exp(xmi) / (D * Sqr(2 * pi))
          O.CurrentY = (yap * fact_Gauss - yn) * 100 / (yx - yn)

         For z = 1 To Np4 '100
          xap = xn_IT + z * (xx_IT - xn_IT) / Np4
          xmi = -0.5 * ((xap - xav) / D) ^ 2 'xav = loiBO("A_val") =val moy de la Gaussienne
          yap = Exp(xmi) / (D * Sqr(2 * pi)) ' D = Val(TAM(17).Text) 'ecart-type
          'sG(z) = sG(z) + Pdp * Exp(xmi) / (D * Sqr(2 * pi)) 'Pdp = IIf(Frcond.Tag < 2, 1, loiBO("poids_stat"))

          xap = (xap - xn) * 100 / (xx - xn)
          yap = (yap * fact_Gauss - yn) * 100 / (yx - yn) 'on ajoutte un facteur 4 pour visulaiser les gaussiennes
          O.Line -(xap, yap), coul
         Next
      End If
      '*****************

      'no) ligne filM1
      O.CurrentX = 103
      O.CurrentY = 3 + iR * 10
      O.Print "rangM " + Format(iR)

   End If 'fin condition

   loiBO.MoveNext
   If loiBO.EOF Then Exit Do
   t = t + 1
Loop


End Sub
```
